# Simultaneous detection and estimation in olfactory sensing

**DOI:** 10.1101/2025.11.01.686013

**Authors:** Chen Jiang, Matthew Y. He, Venkatesh N. Murthy, Cengiz Pehlevan, Jacob A. Zavatone-Veth, Paul Masset

**Affiliations:** Department of Psychology, McGill University, Montréal, QC, H3A 1G1, Canada; Quantitative Life Sciences, McGill University, Montréal, QC, H3A 1G1, Canada; Center for Brain Science, Harvard University, Cambridge, MA, 02138, USA; Department of Molecular and Cellular Biology, Harvard University, Cambridge, MA, 02138, USA; Kempner Institute for the Study of Natural and Artificial Intelligence, Harvard University, Cambridge, MA, 02138, USA; John A. Paulson School of Engineering and Applied Sciences, Cambridge, MA, 02138, USA; Society of Fellows, Harvard University, Cambridge, MA, 02138, USA; Mila - Québec AI Institute, Montréal, QC, H2S 3H1, Canada

**Author notes:** CJ and MYH contributed equally to this work. JAZ-V and PM jointly supervised this work. Correspondence to or.

## Abstract

The mammalian olfactory system shows an exceptional ability for rapid and accurate decoding of both the identity and concentration of odorants. Previous works have used the theory of compressed sensing to elucidate the algorithmic basis for this capability: decoding odor information from the responses of a restricted repertoire of receptors is possible because only a few relevant odorants are present in any given sensory scene. However, existing circuit models for olfactory decoding still cannot contend with the complexity of naturalistic olfactory scenes; they are limited to detection of a handful of odorants. Here, we propose a model for olfactory compressed sensing inspired by simultaneous localization and mapping algorithms in navigation, in which the set of present odors and their concentrations are inferred separately. We implement this split inference in a biologically-plausible recurrent circuit by introducing separate dynamics matched to the distinct nature of presence and concentration, and drawing on the framework of Mirrored Langevin Dynamics. This model can accurately infer presence and concentration at scale. Moreover, its circuit structure can be mapped onto the primary cell types of the olfactory bulb, giving a possible normative account for functional differences between mitral and tufted cells. Our approach offers a general path towards circuit algorithms for probabilistic inference—in olfactory sensing and beyond—that both perform well in naturalistic environments and make experimentally-testable predictions for neural response dynamics.

## 1 Introduction

Animals sense the physical properties of their external world using specialized neural circuitry [1]. To enable rapid adaptation of behavior to the demands of a changing world, early stages of sensory processing leverage the statistical properties of sensed signals [2–5]. This evolutionary adaptation leaves an imprint on the structure of neural circuits [5–7].

The most important statistical analysis problem faced by many mammals is that of odorant detection, of analyzing the composition of a given mixture of scents. Mammals—particularly rodents—rely on olfaction to perform tasks essential for survival, such as detecting predators [8], recognizing conspecifics [9, 10], and locating food sources [11, 12]. Olfactory sensing is computationally challenging, as an animal has access only to intermittent samples of odorants wafted on turbulent plumes of air, from which it must determine which among a myriad of possible sources is present [11, 13–15]. This challenge is exacerbated by animals’ limited repertoire of olfactory receptor proteins, which range in number from ~300 in humans to ~1000 in mice and ~2000 in elephants (Figure 1a-b) [16].

**Figure 1:**
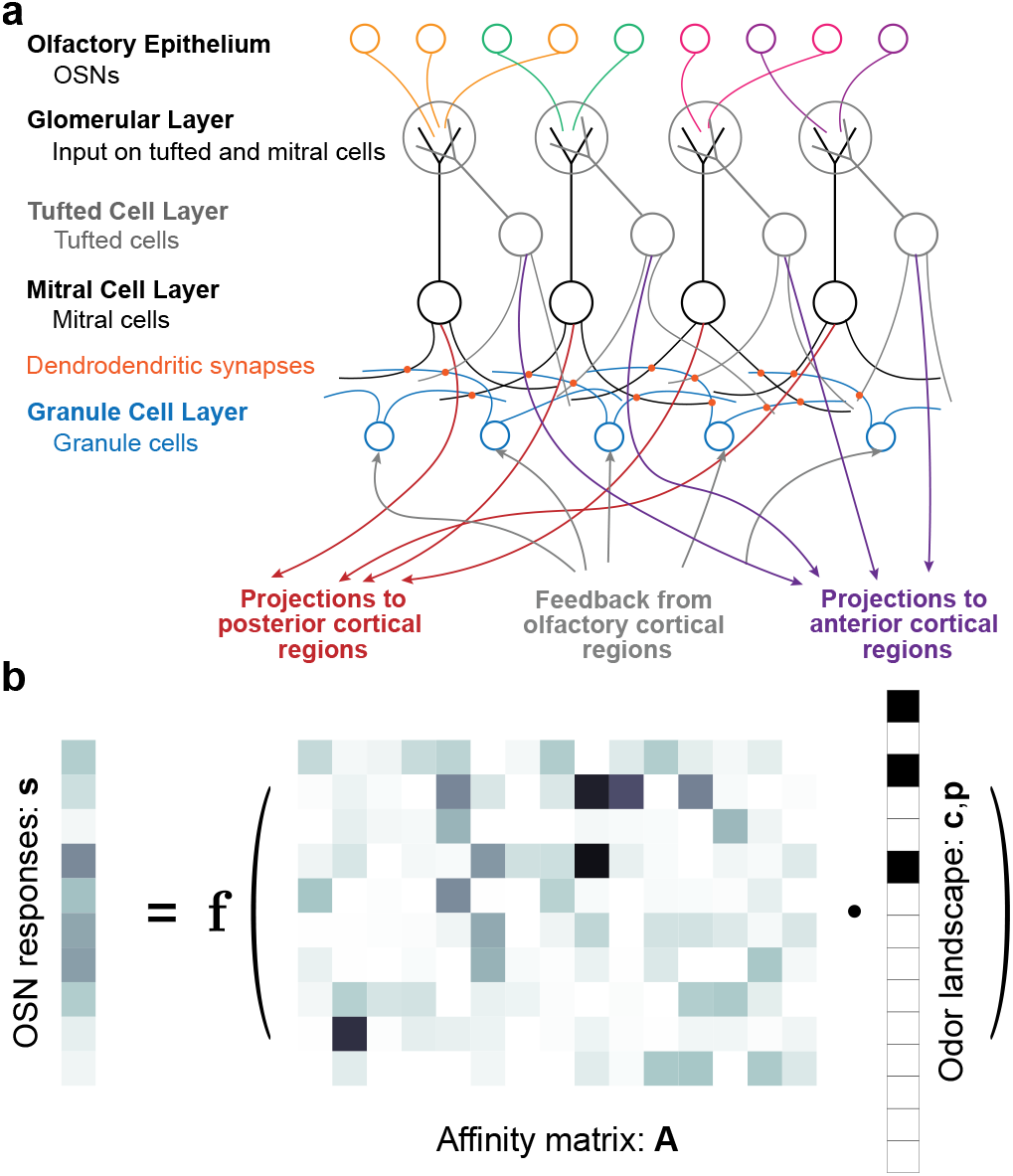
The structure of olfactory sensing. **a**. Outline of the anatomy of the mammalian olfactory bulb (OB). Volatile odorant molecules first bind to olfactory receptors on the surface of the olfactory sensory neurons (OSNs) that tile the olfactory epithelium. Each OSN expresses a single receptor type; humans have a repertoire of about 300 receptor types, while mice have about 1000 [9, 16, 42]. OSNs synapse onto the two primary excitatory projection neuron types of the OB—mitral and tufted cells—in clustered olfactory glomeruli containing axons of OSNs expressing the same, single receptor type [43, 44]. Mitral and tufted cells then project to higher areas, notably including the posterior piriform (primary olfactory) cortex and the anterior olfactory nucleus (AON) [22, 24, 45, 46]. Recurrent connectivity within the mitral and tufted cell layers is mediated via inhibitory granule cells, which make dendro-dendritic synapses with their excitatory partners. Feedback from higher areas to the bulb comes in the form of synapses onto the granule cells. **b**. Dimensionality of the sensing problem. The compressed OSN representation **s** of the olfactory world (*left*) is given by a (stochastic) function *f* (·) of the product of the matrix **A** of the affinities of each OSN to each odorant (*center*) with the sparse, high-dimensional odor scene vector giving the concentrations of which out of millions of possible odorants are present in a given scene (*right*). For illustrative purposes, we show 10 receptors and 15 odorants.

Despite this challenge, rodents can perform many olfactory sensing tasks with exquisite speed and accuracy, often within the hundred-millisecond timescale of a single sniff [12–14, 17–19]. Due to the difficulties inherent in generating tunable yet naturalistic olfactory stimuli in the laboratory, detailed probes of these capabilities have only become possible in recent years [12, 14, 18–20]. This growing body of work has begun to elucidate how the statistics of olfactory landscapes shape perception and detection. Moreover, it has revealed that the different cell types of the olfactory bulb (OB)—the locus of early olfactory processing in mammals (Figure 1a)—display diverse tuning and dynamics in response to odor stimuli [21–25].

How might these remarkable abilities arise? A growing body of theoretical work suggests that the answer may lie in the theory of compressed sensing [26, 27]: it is possible to infer what odorants are present given the noisy activity of olfactory sensory neurons (OSNs) because only a few odorants—tens or hundreds out of a million or so possibilities—are present in a given scene [3, 28–40]. Given a compressed sensing framework, the uniquely-structured recurrent circuits of the OB (Figure 1a) offer a potential substrate for the implementation of a simple inference algorithm.

However, the algorithms proposed by these past works—including our own [39]—have not shown how to simultaneously overcome two key challenges. The first is biological insight: a model must not only be implementable as a biologically-plausible recurrent neural network (RNN), but also have a clear biophysical interpretation, so as to account for cell types, provide understanding of the computation, and generate experimentally testable predictions under complex and realistic settings. For example, the circuit model of Grabska-Barwińska *et al*. [41] maps onto only a single projection-neuron population, collapsing the distinct response dynamics of mitral and tufted cells in the OB [21, 22, 25]. The second is maintaining strong performance when scaling to naturalistic olfactory scenes with a landscape of many rapidly-changing odorants.

Towards the second challenge, separate but coupled inference of interrelated quantities has proven successful in approaching complex inverse problems, as exemplified by simultaneous localization and mapping (SLAM) in robotic naviga-tion [47]. Similarly, in olfaction, odorant presence and concentration can be inferred separately [29, 36, 41]. We will refer to this approach as split-inference in the rest of the manuscript. The split-inference approach allows for the use of flexible, sparsity-promoting priors on presence, which can in principle enable more robust inference in large olfactory scenes. However, inferring presence requires solving a combinatorial problem, as it is inherently binary. This is difficult to implement in a rate-based recurrent circuit, an obstacle not overcome by previous split-inference models [29].

In this work, we propose a framework for olfactory sensing that solves these challenges. Starting from the split-inference formalism, we transform the problem into a form that can be solved using sampling-based probabilistic inference, implemented in a biologically-plausible RNN. To tackle the binary nature of presence, we represent it in two causally linked forms, each serving a different purpose: a probabilistic representation, realized as a continuous surrogate constrained to the interval [0, 1], and a binary representation, reconstructed from the surrogate, that restores a physically appropriate picture of the odor landscape and promotes robustness in concentration inference. The constrained surrogate can be inferred using gradient-based sampling thanks to the framework of Mirrored Langevin Dynamics [48]. Concentration is then inferred only over the odorants the reconstruction marks as present, yielding more robust estimates.

This treatment exploits the distinct physical and statistical nature of binary presence and non-negative concentration, tailoring a different representation and computation to each, and yields an interpretable model whose components align with experimental recordings from the OB. The resulting model enables faster and more robust inference of odorants compared to structurally-similar recurrent circuit models that do not split the inference problem. It can accurately analyze static scenes with hundreds of odorants present out of a repertoire of tens of thousands within the timescale of a single sniff, which significantly exceeds the detection capacity demonstrated by previous olfactory compressed sensing models. At the same time, our model is itself a recurrent circuit that demonstrates rich cell-type-specific neural dynamics in response to odor stimuli which qualitatively resemble those that have been measured in the mitral and tufted cells of the OB [21–25]. Thus, our work lays the groundwork for the development of performant circuit models that can generate detailed, experimentally-testable predictions about response dynamics in the olfactory bulb. Moreover, it introduces a framework for constrained sampling in biologically-plausible circuits that could be applied outside the context of olfaction.

## 2 Simultaneous detection and estimation in olfactory scenes

The starting point for our odorant recognition model is a generative model for olfactory scenes. An odor landscape is determined by which sources are present; in the Montréal cityscape one might imagine coffee alongside bagels pulled freshly from a wood-fired oven. The presence of these sources then implies the presence of a characteristic set of volatile odorants which they emit. These odorants are then transported by turbulent airflow to the nose, where they can at last bind to olfactory receptors, driving activity in OSNs and thus generating a perceivable smell.

We model the olfactory sensing problem as follows. At each physical time *t*, the odor scene is defined by a pair of vectors 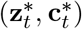, where 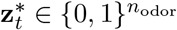 is the binary identity of which odorants are present and 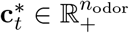 their concentrations. This scene is sensed by *n*_OSN_ olfactory sensory neurons through a receptor affinity matrix 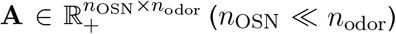, producing a stochastic observation 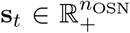 with 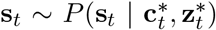. This is the only signal available to the model. The model’s task is to solve the inverse problem: to recover the true 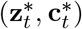 from **s**_*t*_ at each time *t*. To this end the model carries its own internal latents (**z**_*t*_, **c**_*t*_) and infers the posterior *P* (**z**_*t*_, **c**_*t*_ **s**_*t*_), from which it produces point estimates 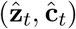 of the true presence and concentration. A full description of the problem statement is given in SI Appendix 2.

Importantly, odorant presence **z**_*t*_ and concentration **c**_*t*_ fluctuate with different statistics and on separate timescales. The presence of an odorant varies slowly, as sources appear and disappear, whereas the measured concentration at the olfactory sensory epithelium displays millisecond-timescale fluctuations over many orders of magnitude, driven both by the physical nature of turbulent transport in air before an odorant molecule arrives at the nose [14, 15, 49], and by the process of sniffing as a mammal actively inhales air into the nasal cavity [49]. As a result, a present odorant’s apparent concentration can fluctuate extensively, even vanishing in the blanks between whiffs, while a truly absent odorant stays absent until the sources change [15].

These physical and statistical distinctions suggest that it should be computationally advantageous to separately infer the presence and concentration of odorants in a given scene. This separation is reminiscent of the principles underlying simultaneous localization and mapping (SLAM) algorithms in robotic navigation, which separate the inference of an agent’s rapidly-changing position from that of a map of slowly-changing landmarks [47]. Probabilistically, it corresponds to treating separately the distribution over presence **z** and the conditional distribution of concentration **c** given **z**: *P* (**c, z**) = *P* (**c z**)*P* (**z**). This allows us to flexibly account for differences between the binary presence and non-negative concentration in both their generative statistical model and their temporal dynamics.

We now specify the two ingredients of this model: a likelihood *P* (**s c, z**) for OSN firing rates given the presence-masked concentration, and the priors *P* (**z**) and *P* (**c z**). For simplicity we take different odorants to be i.i.d. under the prior, a starting point that could be relaxed (see Discussion). Additionally, we do not introduce an explicit model for temporal dynamics in the priors *P* (**z**) and *P* (**c z**); we assume that the priors factorize in time (see SI Appendix 7 for an extension of the model to dynamically-structured priors).

To determine the likelihood, we model the mean activity of OSNs as a linear function of the concentration, with the receptor affinity matrix **A** and a baseline rate 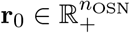. This is accurate within moderate concentration ranges [50] but neglects nonlinear effects known to be important in olfactory sensing; see the Discussion. Following past works [29, 39, 41], we use a Poisson noise model, with the OSNs firing independently, so that for the *j*^*th*^ OSN:

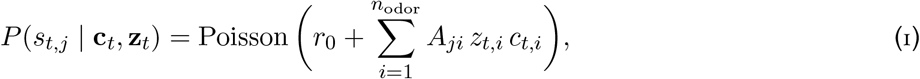

where the OSNs share a baseline rate *r*_0_ and the rate is driven by the presence-masked concentration *z*_*t,i*_ *c*_*t,i*_. We will compare several toy models for the distribution of affinities **A**. In defining **A**, we model all the OSNs expressing the same receptor type and converging on a single glomerulus as single input [9, 16, 42, 43].

For the prior, we use a spike-and-slab structure on the presence-masked concentration *z*_*i*_*c*_*i*_, which reflects the sparsity of natural odor scenes (Figure 2a): a ‘spike’ places most odorants at zero concentration, while a ‘slab’ captures the broad spread of the range of concentrations of the few present odorants. Masking makes this structure easy to impose, since a presence prior favoring directly produces the spike in *z_i_c_i_*. We take the presence prior to be Bernoulli,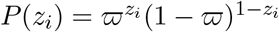, so that the expected number of present odorants is *E*[Σ_*i*_ *z*_*i*_] = *n*_odor_ *ϖ*, and the concentration of present odorants to be Gamma-distributed, *P* (*c*_*i*_ | *z*_*i*_=1) = Gamma(*c*_*i*_ | *α, β*), giving the slab. The parameters *α* and *β* control the shape and Gamma distribution, respectively.

We sample the posterior *P* (**c**_*t*_, **z**_*t*_ |**s**_*t*_) ∝ *P* (**s**_*t*_ |**c**_*t*_, **z**_*t*_)*P* (**c**_*t*_ | **z**_*t*_)*P* (**z**_*t*_) with Langevin-type dynamics realizable as a recurrent circuit (SI Appendix 4). Crucially, this sampler operates online: at each *t* it advances by a single step on the current observation **s**_*t*_, carrying its state forward rather than restarting, so that information accumulates across time even though each update is driven only by the present input.

**Figure 2:**
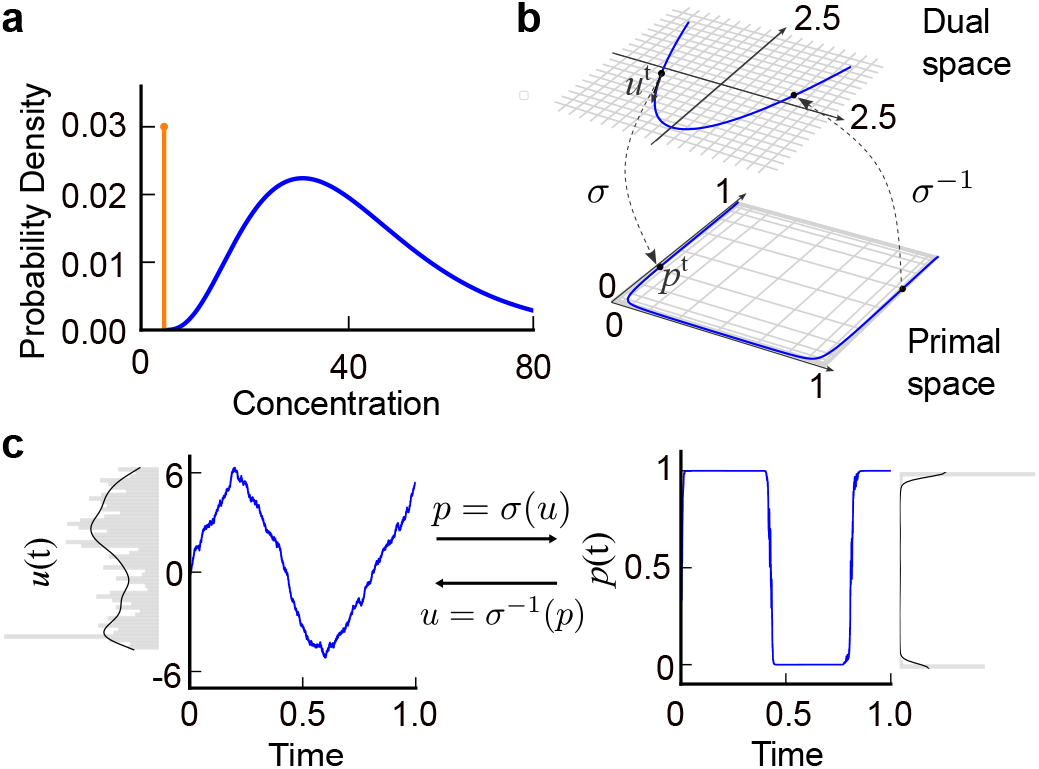
Spike-and-slab priors and Mirrored Langevin dynamics. **a**. Spike and slab prior over concentration, which is a mixture of a Dirac delta function (orange curve) and a Gamma distribution (blue curve). **b**. Mirror map illustrating the nonlinear transformation between dual and primal spaces with the mirror map *σ* and its inverse *σ*^*−*1^. Here, we set the gain of the sigmoidal mirror map *σ* to *γ* = 3. This transformation projects a bounded interval of primal space to an unconstrained dual space. The sampling is performed in the dual space and projected back to the primal space. **c**. Example dynamics of the dual variable **u**(*t*) and the corresponding primal variable **p**(*t*). The mirror map *σ* is a sigmoid with gain *γ* = 5. The distribution of samples in the primal space is concentrated near the boundaries at 0 and 1.

At the level of the computation, a true presence-dependent spike-and-slab cannot be sampled directly with a Langevin-type algorithm, as its point mass at zero is non-differentiable (but see Fang *et al*. [51] and SI Appendix 10). Thus, we adopt a unified Gamma prior on the latent *c*_*i*_ that is the same regardless of *z*_*i*_. This choice is harmless since the masked concentration *z*_*i*_*c*_*i*_ does not depend on *c*_*i*_ when *z*_*i*_ = 0 (but see SI Appendix 6 and Figure S3 for presence-dependent alternatives). In our main model, the spike-and-slab structure is instead recovered from the dynamics, through two coupled but differently gated pathways as will be described later.

## 3 Estimating presence through mirrored Langevin dynamics on a surrogate

Under the split-inference formalism, a model has to address the problem of sampling a binary presence variable. This is the key obstacle to a recurrent circuit sampler, as one cannot directly write down Langevin dynamics that sample a posterior supported on a discrete set [39, 52]. To resolve this issue, we sample the posterior over a continuous surrogate of the presence variable, denoted by 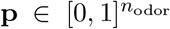. The sampler thus operates on a probabilistic representation of presence, as each component *p*_*i*_ is the posterior probability that odorant *i* is present, *p*_*i*_ ≈ Pr(*z*_*i*_ = 1| **s**).

To account for the fact that the surrogate **p** is constrained to lie inside the hypercube 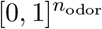, we leverage the framework of Mirrored Langevin Dynamics (MLD), which extends Langevin sampling to distributions subject to convex constraints [48]. We will not endeavor to provide mathematically rigorous guarantees for our models—indeed, not all of the relevant distributions are log-concave (SI Appendix 4)—and will rely on simulations.

The core idea of MLD is to map the constrained “primal” variable to an unconstrained “dual” variable through an invertible “mirror map”, and to run ordinary Langevin dynamics in the dual space [48, 53, 54] (Figure 2b, see SI Appendix 3 for a mathematical overview). To our knowledge this framework has not previously been applied in neuroscience.

Applying MLD to our problem, we express the surrogate 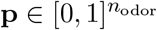 in terms of a dual variable 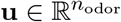 through a sigmoidal mirror map

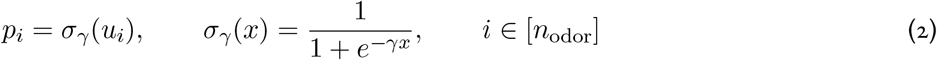

with gain *γ*, where [*n*_odor_] = {1, …, *n*_odor_}. Following the MLD recipe with this map yields a rate-based circuit that samples the posterior; we give the full derivation in SI Appendix 4 and state the resulting model here.

The sampling dynamics are driven by a ratiometric prediction error 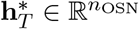, the ratio of the observed OSN activity to the rate predicted by the current estimate *c*_*i*_(*t*) *p*_*i*_(*t*) [39]. We interpret this as a population of projection neurons in the olfactory bulb, labeled *T* because its simulated responses resemble those of tufted cells (see Section 4):

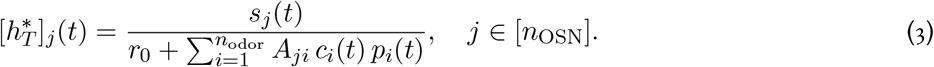

Here 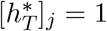 signals a perfect match between observed and predicted activity, so 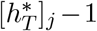 is the relative prediction error. Projecting this error back into odorant space through the affinities gives the bottom-up evidence *e*_*i*_ for the *i*^*th*^ odorant,

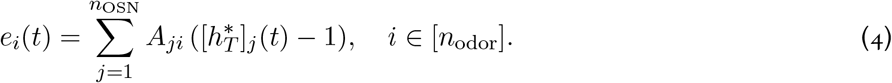

The dynamics of the dual presence variable **u**(*t*) and the latent concentration variable **c**(*t*) are then, for each *i* ∈ [*n*_odor_],

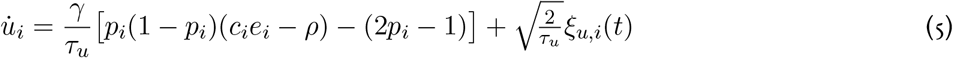

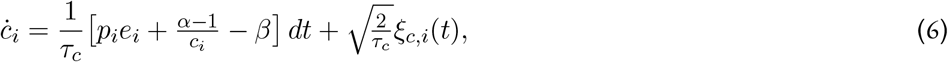

where 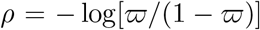 packages the presence prior. Here, ***ξ***_*u*_ and ***ξ***_*c*_ are independent isotropic Brownian noises. These noise terms can be interpreted as arising from various biophysical processes, including vesicle release and thermal fluctuations in membrane potential [29, 55, 56].

Each update couples evidence to prior: the bottom-up evidence *e*_*i*_ is scaled by concentration *c*_*i*_ in the presence update and gated by presence *p*_*i*_ in the concentration update, so the two estimates drive each other. The remaining terms in the **u** update arise from the mirror map (SI Appendix 4). Simulated with modestly large gain, the mirror map turns the smoothly-varying dual signal **u**(*t*) into a nearly binarized presence estimate **p**(*t*) (Figure 2c), so the relaxation yields an interpretable, near-binary readout of surrogate presence. The concentration prior also shapes the dynamics: for *α >* 1 the divisive 1*/c*_*i*_ term acts as a repulsive barrier that keeps concentration estimates from collapsing to zero.

## 4 Cell-type-specific computations and dynamics

Thus far, we have established a dynamical system that can implement split olfactory inference, converting the intractable binary problem into one a recurrent circuit can sample. But it does not yet exploit the key advantage of the split: the flexibility to treat presence and concentration differently. Doing so yields more robust inference and gives rise to distinct neuron populations that resemble mitral and tufted cells, the projection cell types of the olfactory bulb. Here, we outline the model architecture, deferring a full derivation to SI Appendix 4.

### 4.1 Hard gating and two projection neuron types

The physical concentration of an odorant should not be modulated by its presence probability. Gating the concentration estimate by the soft presence surrogate, as in (3), thus introduces a confound: fixing *p*_*i*_*c*_*i*_, if *p*_*i*_ is far from zero or one then *c*_*i*_ will not track the concentration. We therefore gate the concentration update by a binary readout of the soft presence, 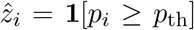, which can be interpreted as a reconstructed binary presence. We then introduce a second ratiometric prediction error signal 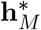, the hard-gated analog of 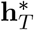:

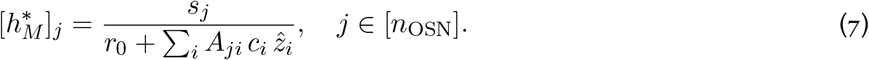

Through hard gating, 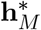 discards evidence from odorants with low presence probability. This improves the signal-to-noise ratio and restores the spike-and-slab structure that the soft model relaxes away.

The complete model thus carries two pathways: a soft-gated 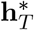 driving presence and a hard-gated 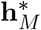 driving concen-tration. This amounts to inferring presence on a probabilistic representation and concentration on a reconstructed binary representation of presence. Replacing the soft evidence *e*_*i*_ in (6) with the hard evidence 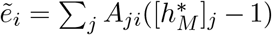, we have the new concentration update:

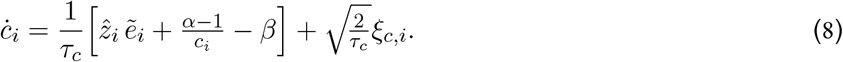

(5) and (8) constitute our complete algorithmic model. The hard-gating pathway is vital: without it the model fails to estimate small concentrations (Figure S4 in SI Appendix). We provide a more thorough discussion in SI Appendix 4.

Because this model performs simultaneous detection and estimation in olfactory scenes, we refer to it as “SDEO”. As shown in Figure 3a-c, our SDEO model accurately tracks the presence and concentration of changing odorants in a simple scene. Here, we simply model the sensitivity matrix **A** as a sparse binary matrix; see Figure S2 for a similar test with a different model for the affinity matrix. Fixing **p** = **1** and running Langevin sampling to infer **c**, we recover the model studied in our previous work [39] (non-separated model in Figure 3c). In Figure 3 we compare this non-separated model to the SDEO model developed above, and see that the SDEO model converges far more rapidly to a more accurate estimate of presence and concentration upon changes in the olfactory scene. This performance improvement, even in a relatively simple scene, is consistent with our conceptual motivations for considering split inference of presence and concentration.

**Figure 3:**
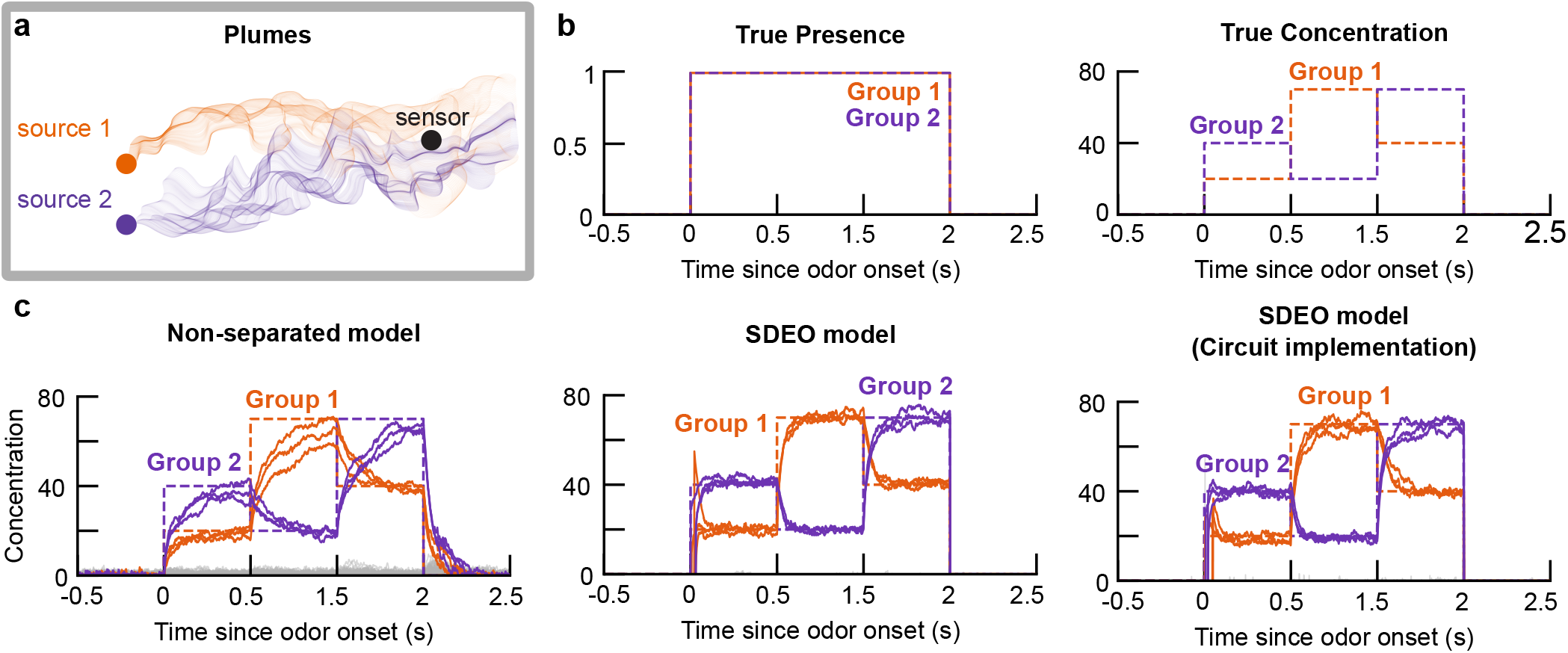
Dynamics of non-separated and SDEO models during estimation of odorants present in a slowly-changing scene. **a**. A static sensor downstream of two odor sources will measure an intermittent, variable mixture of the emitted odorants as they are transported by turbulent airflow. We illustrate this with a sketch inspired by the plume measurements of Nowotny and Szyszka [49]. **b**. In the laboratory, it is challenging to mimic the fast-timescale dynamics of a turbulent plume, but one can generate changing steps of concentration [14, 18, 19]. We model this scenario by selecting two groups of three random odorants each. Each group—intended to model the odor emitted by one source—has common concentration fluctuations, but remains present throughout the entire *in silico* experiment. Within each group, the concentrations of each odorant are identical, but the concentrations for the two groups change independently. We present this simulated odor scene to three models: non-separated (*left*; as in [39]), SDEO (*center*), and SDEO with circuit implementation (*right*). In each plot, the colored lines denote the estimated concentration for the presented odorants, while the gray lines represent those for the background (non-presented) odorants. The dashed line traces true concentration over time. In these simulations, we use a sparse binary affinity matrix. For details of our numerical methods and the corresponding plot using a dense gamma-distributed affinity matrix, see SI Appendix 9D.2 and Figures S1 and S2.

### 4.2 Linearizing the divisive projection neurons

The terms 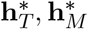 and the prior term (*α* −1)*/c*_*i*_ are divisive, and therefore hard to implement in a rate circuit [57]. Following [39, 58], we linearize them by promoting each division to a population whose fixed point computes it. Consequently, we have the projection neurons **h**_*T*_, **h**_*M*_ and an auxiliary population **f**, with dynamics

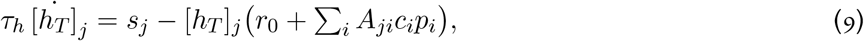

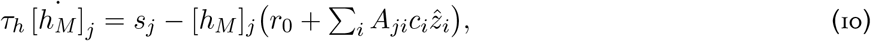

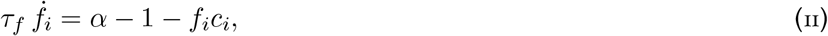

whose fixed points are 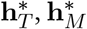, and (*α*−1)*/***c** respectively.

Substituting these populations for the divisions yields the full circuit dynamics, written directly in terms of the cell activities:

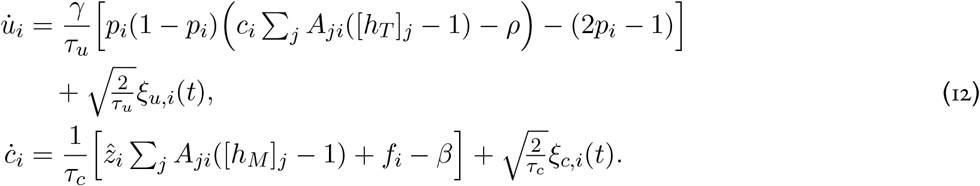

The presence update is driven by the soft-gated **h**_*T*_ and the concentration update by the hard-gated **h**_*M*_, with the prior division carried by **f**. For fast auxiliary timescales, this five-population circuit reproduces the algorithmic dynamics (Figure 3).

### 4.3 Mapping model circuitry onto the olfactory bulb

We can map these dynamics onto the circuit architecture of the olfactory bulb [7, 25]. As they are excited by the OSN input, we interpret **h**_*T*_ and **h**_*M*_ as the two principal classes of projection neurons in the OB (tufted and mitral cells respectively). Then, the concentration estimate **c** and presence estimate **p** are encoded by two classes of local interneurons (granule cells), which inhibit the projection neurons and gate each other’s dynamics via dendrodendritic synapses [25, 39, 59]. The **f** neurons required to linearize the prior can then be interpreted as a form of cortical feedback onto the granule cells [43]. As noted in our previous work [39], the multiplicative gating of projection neuron inhibition by granule cells is consistent with known physiology [60, 61].

With this coarse mapping in mind, we use time constants that are comparable to those measured in experiment for the major cell types of the olfactory bulb: we set 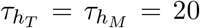 ms to roughly match the primary projection neurons of the bulb, the mitral and tufted cells [62], and we set *τ*_*u*_ = *τ*_*c*_ = 30 ms to roughly match the primary local interneurons of the bulb, the granule cells [63]. Although these timescales are not well-separated, inference remains reliable, consistent with other works using this linearization [39, 58]: the circuit closely tracks the algorithmic model (Figure 4) and still far outperforms the non-split model (Figure 3c, left).

**Figure 4:**
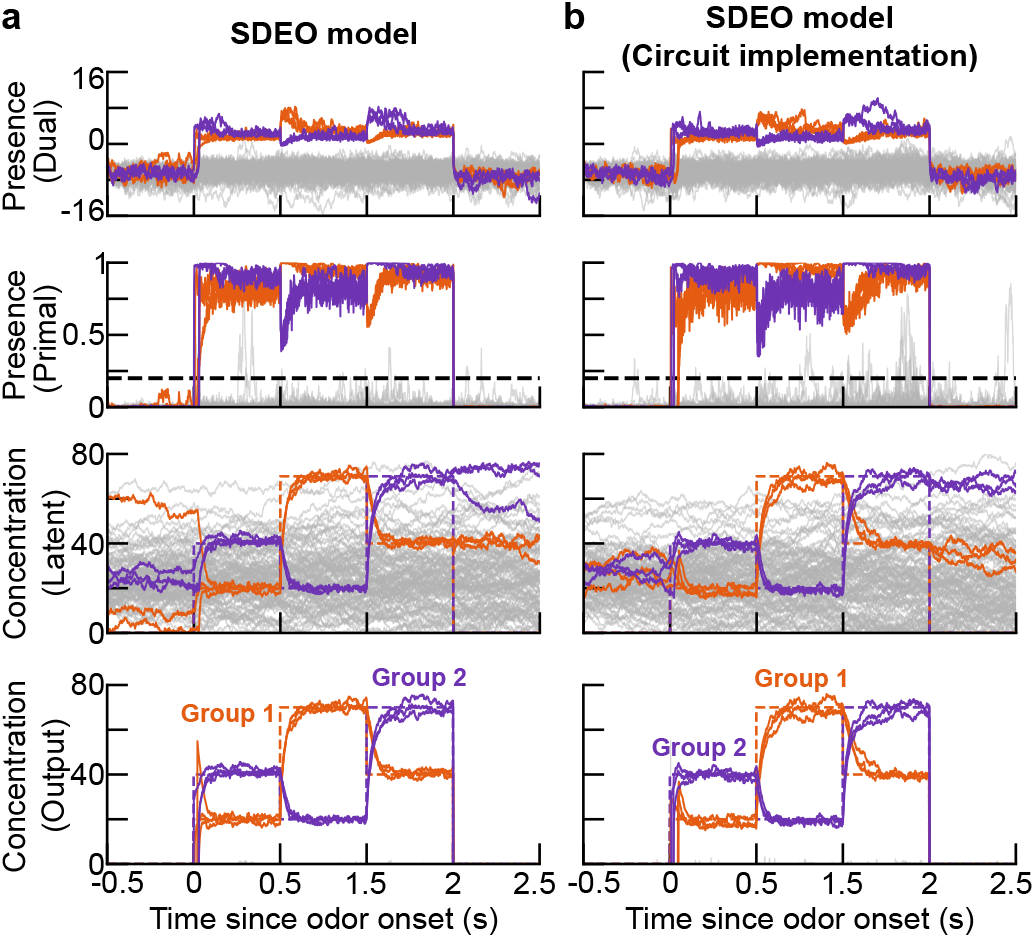
Internal dynamics of SDEO models during estimation of odorants present in the slowly-changing scene shown in Figure 3. In **a** and **b**, we show the basic SDEO model and its circuit implementation, respectively. The first row shows the estimated dual-space presence variable **u**, the second row shows the estimated presence in the [0, 1]-bounded primal space **p** (with a dashed line marking the threshold used to binarize presence during inference), the third row shows the latent concentration estimate **c** (which does not return to zero for non-present odors because of our choice of prior), and the fourth row shows the output concentration estimate *c*_*i*_*p*_*i*_. The panels in the fourth row are identical to the corresponding panels in Figure S2c. In each plot, the colored lines denote the estimated concentration for the presented odorants, while the gray lines represent those for the background (non-presented) odorants. Fluctuations in the primal space presence occur for non-present odors with low estimated concentrations, for which the presence can fluctuate without substantially changing the masked concentration *c*_*i*_*p*_*i*_. This is visible in the fact that the output concentrations for non-present odors remain near zero in the fourth row. In these simulations, we use a sparse binary affinity matrix. For details of our numerical methods and the corresponding plot using a dense gamma-distributed affinity matrix, see SI Appendix 9D.2 and Figures S1 and S2.

### 4.4 Dynamics and tuning of the hard- and soft-gated projection neurons

Previous computational models have usually ignored the distinction between the two dominant classes of projection neurons, the mitral and tufted cells [28, 29, 39, 41] (but see [64]). Our model possesses two classes of hard- and soft-gated neuron populations, which lets us ask how their responses compare to the known properties of mitral and tufted cells.

In simple scenes modeling the static odorant mixtures presented in experiments, we find that our model recapitulates the differences in response amplitude, concentration dependence, and duration characteristic of mitral and tufted cells. At the population level, both classes exhibit a transient burst of activity at odor onset (Figure 5). Soft-gated neurons then relax to a plateau above the baseline whose early amplitude and duration increase monotonically with concentration (Figure 5a-c,e), consistent with the response dynamics of the single projection class in our previous work [39] and with what has been observed experimentally in tufted cells [21, 22, 24]. In contrast, hard-gated neurons show a biphasic pattern: a steep suppression below baseline after the burst and then a slow rise to a second peak before returning to the baseline activity level. Higher concentrations evoke deeper initial drops and higher subsequent rise, i.e., a greater modulation depth (Figure 5a-b,d,f). This is consistent with the more complex dynamics of mitral cells [21, 22, 24].

**Figure 5:**
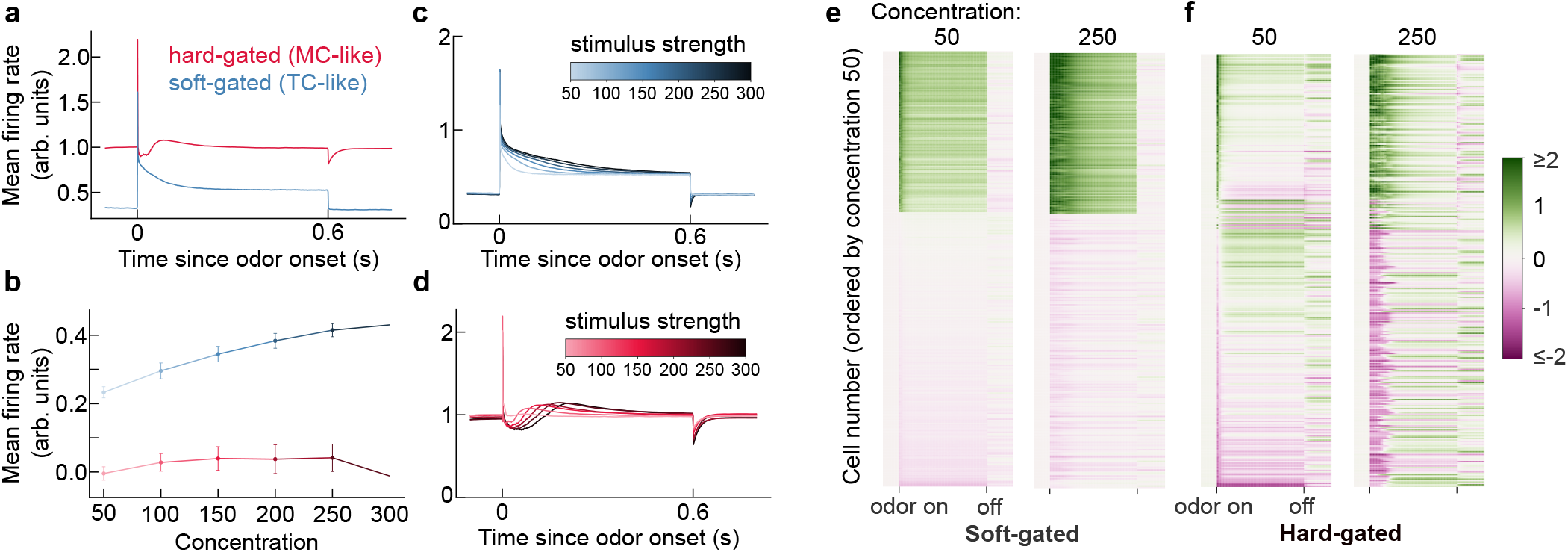
Comparison between putative mitral cells 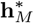 (hard-gated MC-like) and putative tufted cells 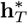(soft-gated TC-like). A set of fixed odor stimuli is present at 0s and withdrawn at 0.6s. We set the Gamma distribution as the concentration prior with shape parameter *α* = 5 and rate parameter *β* = 0.1, and the Bernoulli prior for presence with *ϖ* = 0.01. **a** Mean firing rate over time of two projection neurons. Upon stimulation, both the hard-gated neuron and the soft-gated neuron exhibit a transient burst of activity. After the burst, soft-gated neurons reduce to a plateau higher than baseline, while hard-gated neurons drop immediately, then rise again before settling into baseline activity. When the stimulation disappears, softgated neurons drop to baseline instantly, whereas hard-gated neurons exhibit a negative peak and then converge to baseline smoothly. **b**. Mean firing rate as a function of odor concentration, averaged over the first 100 ms after odor onset. Error bars indicate standard error across trials. The soft-gated neurons respond monotonically to concentration. **c**. Mean firing rate of soft-gated neurons under varying odor concentrations. Darker color indicates stronger stimulation. At odor onset, higher concentrations evoke longer-lasting responses, yet all activities converge to the same plateau. **d**. Mean firing rate of hard-gated neurons under varying odor concentrations. At odor onset, higher concentrations evoke deeper initial drops followed by a slower but higher rise. At odor offset, higher concentrations evoke lower negative peak, after which all responses converge back to baseline **e, f**. Baseline-subtracted activity of 300 soft-gated neurons and 300 hard-gated neurons, ordered by the response magnitude at concentration 50 (averaged over 50 ms after odor onset). A subset of soft-gated neurons and hard-gated neurons exhibits a stronger and loner peak at odor onset when concentration increases. The rest soft-gated neurons are stable, remain baseline or slightly below, whereas the remaining hard-gated neurons exhibit a negative peak, after which some recover to baseline and others rise above baseline. For additional single-neuron responses, see Figure S5.

Averaged over the first 100 ms after onset, soft-gated neurons (putative tufted cells) increase their response monotonically with odorant concentration, whereas hard-gated neurons (putative mitral cells) show no clear monotonic trend (Figure 5b)—consistent with experiment [24]. At the single-neuron level, both classes include cells that are activated and suppressed relative to baseline (Figure 5e-f and Figure S5). The activated soft-gated tufted-like neurons exhibit sustained activity during stimulation, while suppressed soft-gated tufted-like neurons are stable, remaining at or slightly below baseline. In contrast, activated hard-gated mitral-like neurons show transient bursts, which are strengthened and prolonged as concentration increases, while the rest show a negative peak, after which they follow one of three patterns: persistent suppression, return to baseline, or a rise above baseline. These contrasting motifs could be tested in future experiments.

The model’s response dynamics still deviate from experimental observations, particularly in onset latency, sparsity, and the response threshold [21, 22, 24, 65]. We will return to this issue, and discuss how well our model can be mapped to the circuitry of the olfactory bulb, in the Discussion. Nonetheless, our results illustrate the potential of our framework to generate testable predictions for cell-type-specific dynamics.

### 4.5 Sniff-modulated inference model

So far we tested the model dynamics under quasi-static odorant mixtures, where external odor concentration remains constant over long temporal intervals. In natural olfaction, the odor input is dynamically modulated by sniffing [66]. OSNs and principal projection neurons have been shown to be differentially aligned within the sniff phase [18, 21, 22]. To examine the phase response of the neuron classes in our model, we extend the model to incorporate sniff-modulated inference.

Although parameters of sniffing can vary drastically depending on an animal’s state [66], we use a sinusoidal oscillation to model the sniff cycle in order to keep the model simple and tractable. In addition, OSN discharge has been shown to lag behind inhalation onset, reflecting the time required for odorant–receptor binding and transduction. Because our main goal is to examine the phase-related differences in activity of the two principal neuron classes, we do not explicitly model the delay between inhalation onset and OSN discharge. Instead, we assume that OSN activation begins at inhalation onset, which we define as the sniff cycle onset.

Under this sniff-modulated stimulus, while the system-level concentration and odor presence estimates follow the sniff cycle (Figure 6a,b), the dynamics of the mitral-like and tufted-like neurons differ significantly. Both their mean firing rate (Figure 6c) and their phase of peak response (Figure 6d) are different, and such observations are robust to the sniffing frequency (Figure S6). These differences match experiment, including an offset in peak response phase across single cells of ~ *π/*2 (Figures 6d and S6d) [21, 22], and the greater stability and clearer mode of individual soft-gated tufted-like neurons (Figure S7).

**Figure 6:**
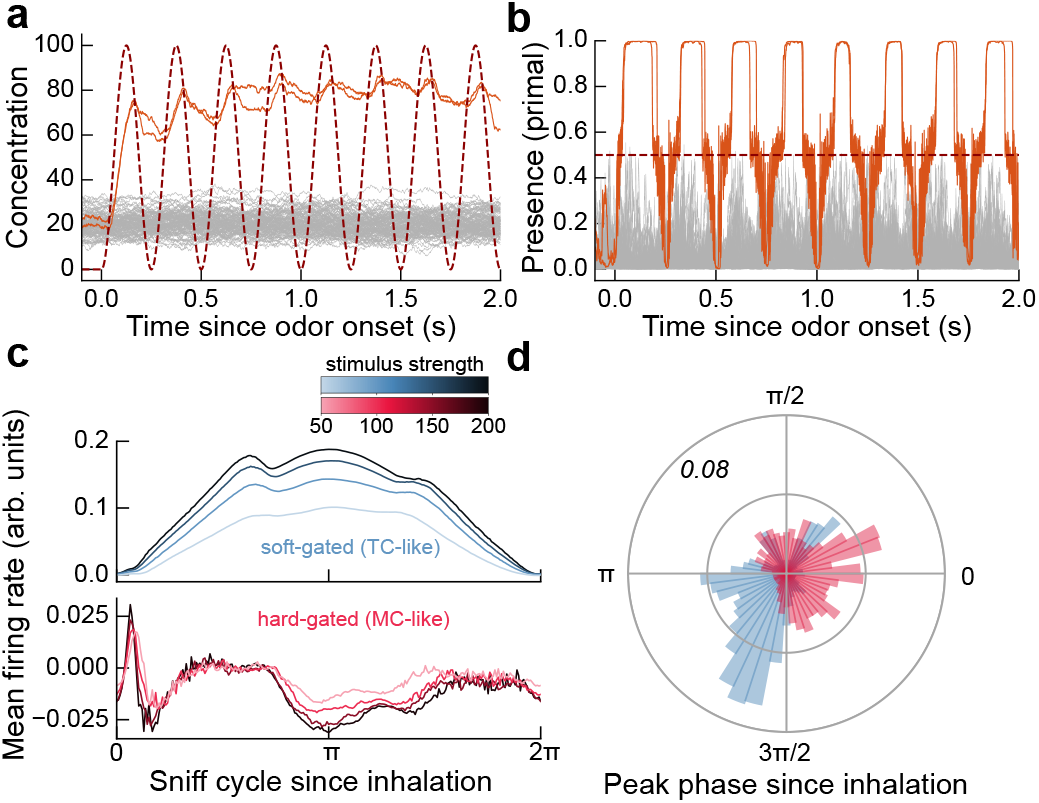
Sniff-coupled dynamics of the SDEO model. External odor stimuli are held fixed, but the input concentration received by OSNs is modulated by sniffing, modeled here as a sinusoidal oscillation. **a**. Example concentration estimates across odors. Orange traces indicate present odors, gray traces indicate absent odors, and the dark-red dashed trace shows the sniff-modulated input concentration received by OSNs. The inferred concentration approximately tracks the true input over a few successive sniff cycles. **b**. Inferred odor presence for the same simulation. Present odors rapidly cross the decision threshold, the dark red dashed line, at an early phase of the sniff cycle, whereas absent odors remain mostly below threshold. **c**. Baseline-subtracted mean firing rate in a sniff cycle of hard-gated projection neurons (red) and soft-gated projection neurons (blue) over concentration. The soft-gated neurons exhibit a clear monotonic trend with stimulus strength. While the activity of hard-gated neurons is distributed more broadly without a clear mode. **d**. Distribution of single cell peak response phases for soft- and hard-gated projection neurons, showing distinct phase preferences between the two types.

## 5 Scaling of olfactory inference

Beyond biological plausibility, an inference algorithm must be performant. We therefore evaluate our model with two scaling simulations: the first quantifies the model’s ability to estimate odorants on a timescale of hundreds of milliseconds, the second examines how the required number of OSNs scales with the size of the potential odorant dictionary.

As a baseline, we use the non-split model of [39]. Since it lacks explicit presence estimation, we approximate presence by binarizing its concentration estimates at half the true concentration. Our evaluation criteria are stricter than in [39], with a thorough discussion in SI Appendix 9.

### 5.1 Speed of inference

We first evaluate rapid inference of odor presence. With a potential odorant dictionary of size 1000 and 300 OSNs, the SDEO models can detect the concentration of ~40 simultaneously presented odorants with at least 80% accuracy and mean absolute error ≤25% · *c*_True_ (Figure 7b,d; see also Figure S9). For presence estimates we can also see a similar but slightly improved limit of detection at ~50 simultaneously presented odorants. On the other hand, the non-separated model can only detect concentration of less than 10 odorants reliably, and detect presence accurately for at most 25 odorants. Moreover, we can see that SDEO models have a significantly faster convergence rate where they reach steady states after ~100 milliseconds, while the non-separated model needs seconds to converge.

**Figure 7:**
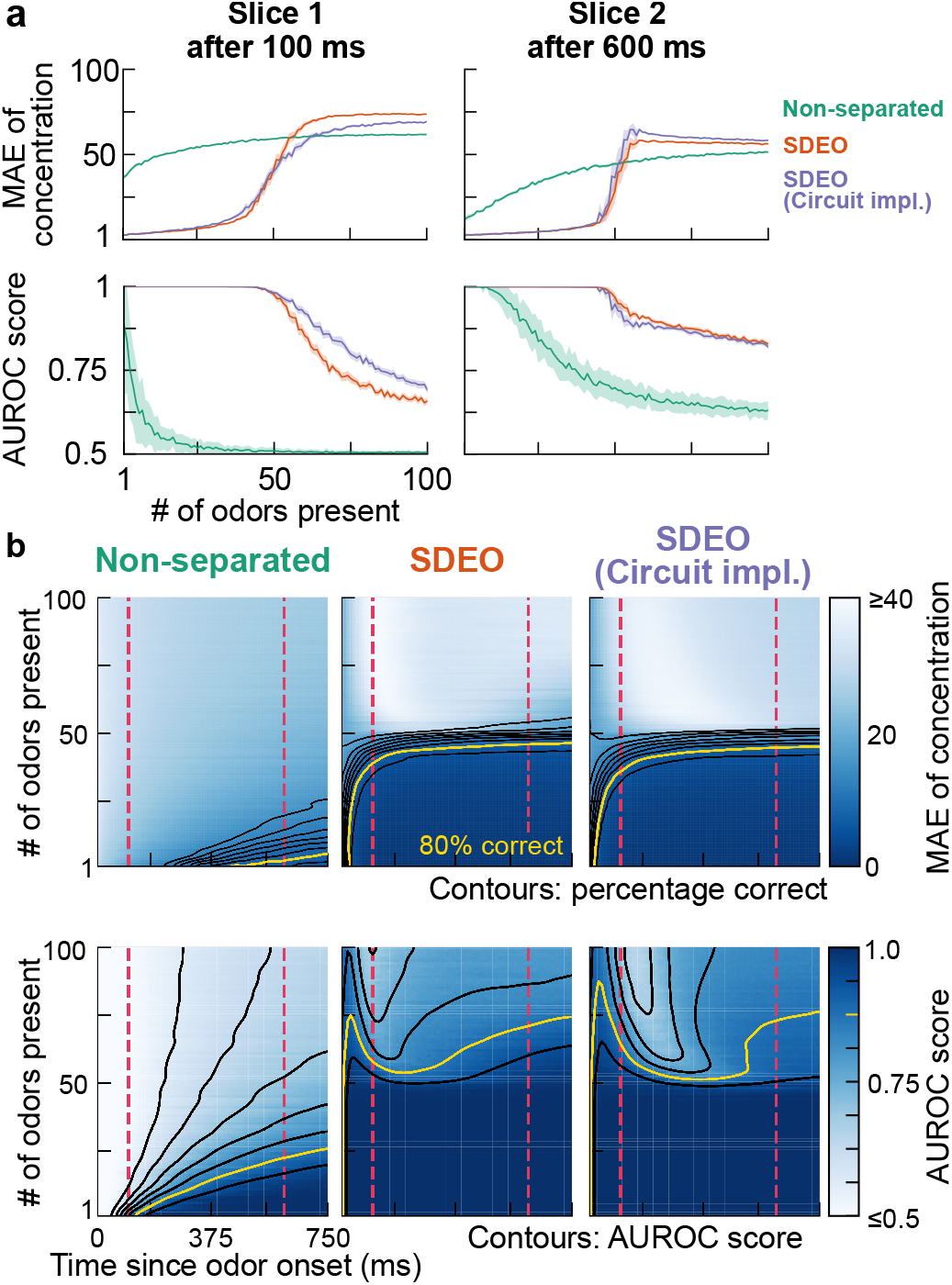
Improvement in fast detection of multiple odorants through separation of inference. We evaluate the same three models as in Figure 3 in a series of simulation where increasing numbers of odorants are simultaneously presented. In each run, in a set of 1000 odorants, a number of them are randomly selected and presented to the model for a duration of 0.75 s. We increase number of presented odors from 1 to 100, while repeat each setting for 40 times, compute the metrics and then take the average as final results. The shaded areas in **a** and **c** show ±1.96 · SEM (representing 95% C.I) over realizations throughout. The first and second rows assess the models’ performance in odorant concentration and presence estimation, respectively. **a**. Mean absolute error (MAE) of estimated concentration as a function of the number of odorants present at two timepoints after odor onset. **b**. Heatmap of mean absolute error over inference time and number of presented odorants, with smoothed contours of correct detection fraction overlaid. **c**. AUROC score as a function of the number of odors present at two timepoints after odor onset. **d**. Heatmap of AUROC score over inference time and number of presented odors, with smoothed contours overlaid. For details of implementations and corresponding figures using dense Gamma sensing matrices, see SI Appendix 9D.3 and Figure S9.

Notably, the SDEO models exhibit a clear phase transition. Although having higher capacity compared to the non-separated model, the SDEO models collapse dramatically once the number of presented odorants exceeds their capacity, indicating qualitative changes in the network dynamics (Figure 7b,d; see also Figure S9). In contrast, the performance of the non-separated model deteriorates more gradually as the number of presented odorants increases. Moreover, the transition occurs simultaneously at ~50 odorants for both concentration and presence, reflecting their coupling—the model exploits their joint dynamics rather than solving two independent tasks.

### 5.2 Detection capacity

From a compressed-sensing perspective, the key question when evaluating a model is how the number of sensors required to achieve a particular detection capacity scales with the size of the dictionary of potential odorants [26, 67]. In our work, we define detection capacity as the largest number of simultaneously present odorants a model detects with the desired accuracy (SI Appendix 9C.4).

In Figures 8 and S10, we examine the scaling of detection capacity with dictionary size and sensor repertoire across different combinations of models and sensing matrix. In the first two columns, we can see the SDEO model significantly outperforms the non-separated model across the entire heatmap. Thus, splitting the inference of presence and concentration improves the speed and accuracy of inference, yielding higher capacity. Upon closer inspection of the SDEO model’s dynamics, we find that the ultimate bottleneck of capacity arises from presence estimation, which drives the sampling of concentration.

**Figure 8:**
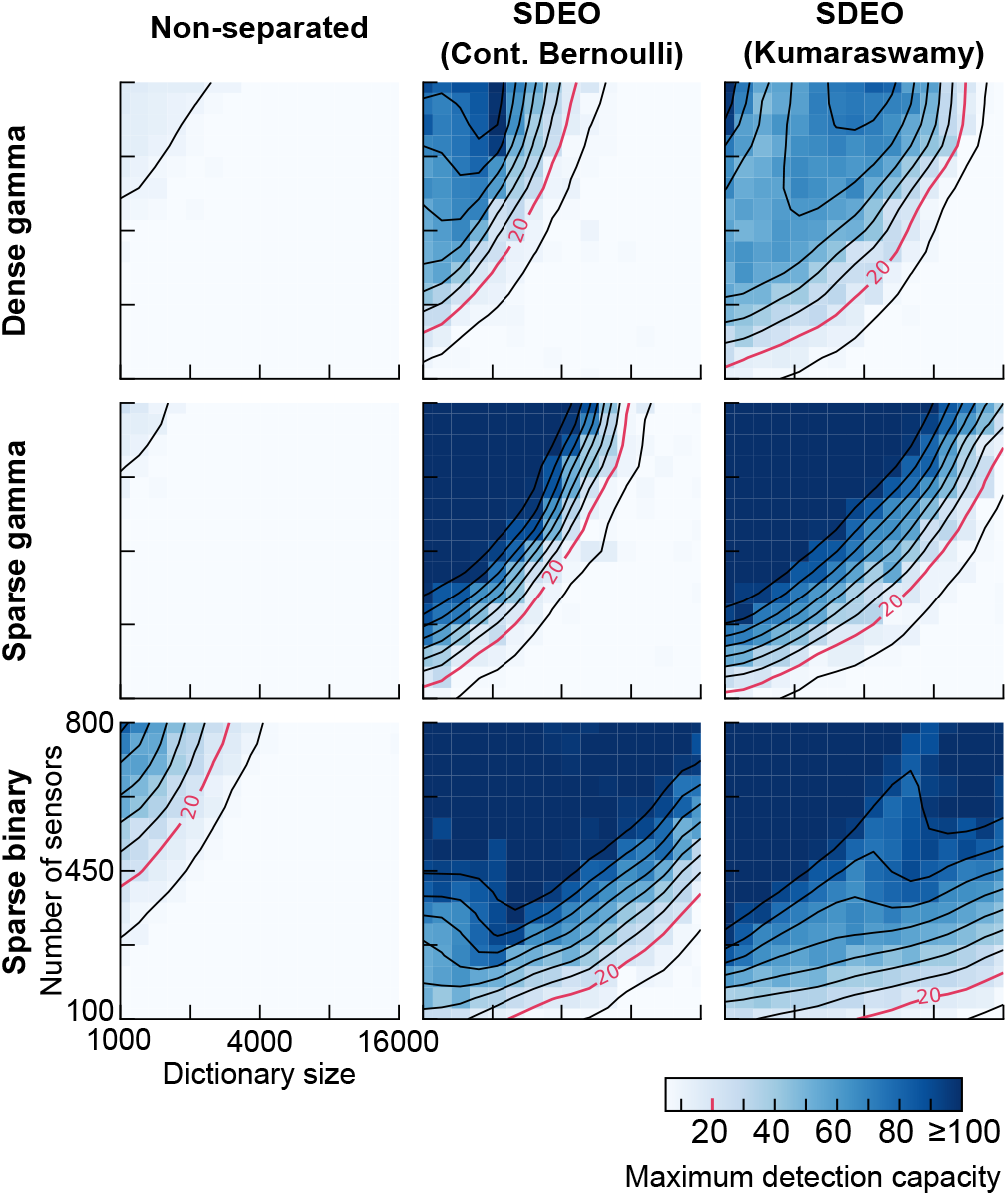
Scaling of detection capacity with dictionary size and sensor repertoire for different priors and sensing matrix models. The three columns correspond to three models—non-separated, SDEO and SDEO with truncated KS prior on the soft presence—and the three rows correspond to three types of affinity matrices. These are: dense gamma, whose entries are i.i.d. random variable following Gamma(0.37, 0.36); sparse gamma, obtained by applying a 0.1 sparsity mask to a dense gamma matrix; and sparse binary, whose entries are i.i.d. random variable following Bernoulli(0.1). Each heatmap shows the maximum detection capacity assessed by presence estimates for combinations of sensors counts (from 100 to 800, equally spaced linearly) and dictionary size (1000 to 16000 equally spaced on a log scale). The maximum detection capacity *κ*_AUROC_ is defined as the largest number of simultaneously presented number of odorants that the model can detect with a AUROC score ≥ 0.85. Smoothed contours are overlaid and can be interpreted as the required number of sensors to maintain a certain capacity as a function of dictionary size. The total inference time duration is 0.2 s for all runs, and the value in each cell of the heatmap is the average of three independent runs. For further discussion on affinity matrix and scaling capacity, see SI Appendix 8. For additional details of implementation and the corresponding figures using mean absolute error to compute maximum detection capacity, see SI Appendix 9D.4 and Figure S10.

### 5.3 Improving capacity by modifying the presence prior and sensing matrix

As detection capacity appeared to be bottlenecked by presence estimation, we sought to improve it by altering the presence prior. The Bernoulli prior we have used poorly reflects the bimodal nature of presence probability, which clusters near 0 and 1, whereas the continuous Bernoulli density is monotonic on [0, 1] for our parameters (Figure S11). We therefore seek a bimodal prior on [0, 1] with a tractable density and log-density gradient. The Beta distribution is the obvious candidate, but its density and log-density gradient are costly to compute; we instead introduce a truncated Kumaraswamy (KS) distribution [68–70] (for derivation of this variant of the model see SI Appendix 5).

The SDEO model with this prior outperforms the continuous-Bernoulli version (Figures 8, S10): for a given dictionary size, fewer sensors are needed to reach a given capacity, and the limiting dimensionality increases. This likely arises because the bimodal prior better captures the generating distribution of presence, facilitating sampling.

Sparsifying the affinity matrix yields similar gains, with capacity increasing from dense Gamma to sparse Gamma to sparse binary; we discuss the connection to compressed-sensing theory in the Discussion (see also SI Appendix 8 and Figure S12). In sum, with an appropriate prior and affinity matrix, SDEO detects and estimates the concentration of up to roughly a hundred odorants from a repertoire of tens of thousands (Figures 8, S10)—far exceeding our previous model and other olfactory compressed-sensing approaches.

## 6 Discussion

In this work, we combined insights from simultaneous localization and mapping in robotics [47], compressed sensing [26], and mirrored Langevin sampling algorithms [48] to propose a biologically-constrained model for odorant sensing (Figure 1-3). Building on these, our key step is to tailor the circuit’s representation and dynamics to the distinct statistics of presence and concentration. This step is essential for the model to work, and it directly leads the model to recapitulate experimentally-observed response properties of the two main projection neuron classes in the bulb, mitral and tufted cells (Figure 5-6). We further show that our approach scales to large sensory scenes, successfully detecting the presence and estimating the concentration of tens to hundreds of odorants amongst thousands of candidates (Figure 7-8).

Our model suggests a map between anatomy and computational function, with the tufted cells contributing to the odor presence detection and the mitral cells to the concentration estimation and with the dynamics of each cell type being broadly consistent with experimental observations [21, 22, 24] (Figures 5 and 6). In contrast to previous accounts of the mitral/tufted (M/T) split [64], our model proposes that the two cell types interact through indirect, reciprocal multiplicative gating rather than additive excitation—an effect that future experiments could probe. As in our previous work [39], the M/T cells encode a ratiometric prediction error between the current OSN activity and the prediction from the concentration estimate encoded in the granule cells. By introducing hard and soft gating, we form two types of prediction-error neuron populations, corresponding to two types of prediction errors—probabilistic versus reconstructed. Existing models for predictive coding in other sensory systems that incorporate multiple cell types generally assume Gaussian noise and encode subtractive prediction errors [71]. Contrasting subtractive versus ratiometric errors encoding formulations in modeling is a direction for future work.

Although our model further points to cell-type specific computations, we do not provide a biophysical interpretation of these computations and leave to future work a more detailed model of the granule cells and other cell types of the OB [39, 59]. Our model nevertheless provides avenues to understand the computational role of these cell types. For example, our model suggests two classes of granule cells that compute presence and concentration and connect to tufted and mitral cells respectively via dendrodentric synapses. This architecture is consistent with the distinct superficial (or type III) and deep (or type II) granule cells that preferentially target tufted and mitral cells respectively [25, 72, 73] and emerging experimental data pointing to distinct functional roles across these granule cell types [74]. However, we do not provide a full biophysical implementation for the granule cells as there is a lack of experimental data measuring their activity across large panels of odorants that would allow us to constrain their representations.

Our model also does not resolve how the information from the mitral and tufted cells is used by downstream cortical areas. The major cortical projection targets are distinct across the two cell types—with, roughly speaking, tufted cells projecting predominantly to anterior cortical regions like the AON and mitral cells projecting to more posterior regions like the piriform cortex—but there are secondary projections with substantial projection from mitral cells into the AON and some projection from tufted cells into piriform cortex [22, 45, 46]. This implies that the computational function of cortical areas is not readily mapped onto the presence and concentration axes. Representations in piriform cortex have been shown to carry both concentration and presence information [75, 76], consistent with inputs from both mitral and tufted cells. Piriform also carries contextual information such as location [77], which could play a role in adapting the odor-processing prior [78, 79]. Less is known about tuning to concentration in the AON, but experimental data is consistent with a role in detection of odor identity for generalization and categorization which would be mediated by the tufted cell inputs in our model [80–83]. The olfactory tubercle, which plays for the olfactory system a similar role to the striatum and mixes odor and valence representations [84, 85], is innervated by both mitral and tufted cells [22, 45]. Our model suggests that future experiments should carefully control and vary the statistics of odor concentration and presence across experimental conditions to disentangle how the mitral and tufted cell projections differentially impact olfactory cortical areas.

A similar probabilistic formulation of olfactory scene analysis was used in earlier work by Grabska-Barwińska *et al*. [29, 41]. Our model differs in how the inference is carried out, and as a result better accounts for the circuit dynamics in the OB [29] and provides insight into the functional roles of the distinct projection cell types [41]. Grabska-Barwińska *et al*. demonstrated two approaches: a Gibbs sampler for presence [29], which preserves exact binary estimates but has no rate-circuit realization, and a variational circuit [41] that maps onto only a single projection-neuron population. Neither was shown to scale to repertoires in the thousands. Our earlier non-separated circuit [39] likewise reached only tens of odorants out of ~8000. By splitting the inference and tailoring the circuit to presence and concentration, our model overcomes both the cell-type and scaling limitations. Approaches based on bottom-up morphologically-detailed biophysical models have provided precise biophysical predictions of mitral/tufted cell distinctions [86], but these models differ from our normative approach in that they do not scale computationally to perform inference of the composition of complex odor landscapes.

Experimental probes of cell-type-specific response dynamics in the OB have been limited by the difficulty in controlling odor stimuli. Unlike for vision or audition, it is challenging to deliver tunable olfactory stimuli that mimic the statistics of those encountered in the natural world. Typical experiments in psychology and neuroscience have delivered static odor flows at high concentrations over timescales of hundreds of milliseconds. It has only recently become possible to combine recordings of neural activity with rapid control of the physical stimulus, through rapidly switching air flows for odor delivery [14, 18, 87, 88], invasive approaches controlling breathing [50], or direct optogenetic activation of olfactory sensory neurons [89–92]. Thus, it is now possible to test predictions from a model like ours about the neural dynamics in response to temporally-structured stimuli.

Despite the richness of its predictions, our model provides a partial picture of olfactory bulb computation. The general SLAM framework offers flexibility we have not yet exploited; it would let our model account for the turbulent dynamics of the odor landscape. SI Appendix 7 sketches how the static prior could become dynamic, with a transition model that tracks the scene’s two timescales separately—slow changes in which odors are present, and fast concentration fluctuations from sniffing [18] and turbulent transport [15, 49]—realized as evolving cortical feedback. This adaptive feedback would enable richer cell-type-specific predictions [21, 24, 58] and better inference in complex scenes.

We also used a simplified model for OSN responses that assumes that receptor affinities are independent and identically distributed (i.i.d.), and that the mean firing rate of each OSN is linear in concentration. The i.i.d. assumption aids systematic analysis of the scaling properties of the model but is an idealization. Under it, sparser random affinities improved inference capacity, qualitatively consistent with compressed-sensing theory under Gaussian noise [67]. It also aligns with work showing that affinities optimized to maximize the mutual information between odor concentrations and OSN activity (under linear or simple nonlinear firing-rate models with Gaussian noise) are sparse [3, 4, 32], and with evidence that simple changes in receptor expression can improve coding efficiency [93]. In these optimized models the nonzero affinities are broadly distributed—which our sparse Gamma model roughly captures—though individual elements carry structure beyond long tails [4]. Investigating how optimized sensitivity matrices and non-linear OSNs affect our results is left for future work, as the optimization procedures used in past works are too computationally expensive for the scaling analysis performed here.

Our model does not incorporate correlations between the presence of different odorants, which naturally arise between odorants that share a common source. Such environmental correlations can shape the distribution of optimal receptor affinities [4], and Tootoonian and Schaefer [40] have recently proposed a circuit basis for the encoding of correlated odorant priors in the olfactory bulb. Their construction leverages sister mitral cells, which receive input from the same receptors but are connected to different granule cells [89]. Our model does not leverage this structure, and incorporating structure and hierarchies into the prior will be important for testing its ability to parse realistically-structured scenes.

In the present work we consider a non-distributed coding scheme, in which each granule cell represents the estimate of the presence or concentration of a specific odorant, whereas previous studies have shown distributed coding in granule cells [7, 80]. In our previous work on the non-split version of this inference model [39], we showed that linearly distributing the odorant code—writing the estimated concentration vector as **c** = **Γg** for model granule cell activities **g** and a decoding matrix **Γ**—improved performance if one chose **Γ** to cancel correlations in OSN responses induced by a dense sensing matrix. It is possible that taking advantage of this geometry would reduce the gaps between different sensing matrix ensembles which we observed in Figure 8. However, the simple linear mixing we used in [39] will not suffice here, as the additional nonlinearities in the model dynamics fix a preferred basis. Naïvely following the linear mixing approach leads to a model without a clear circuit interpretation. Determining how to construct a split inference model with fully distributed coding will be an important task for future work, and a prerequisite for more direct comparison of cell-type-specific dynamics in models with those measured in the OB.

More generally, our work points to compositionality of neural representations, as has been studied across other modalities and neural circuits [94–96]. Decoupling concentration estimation and odor detection would allow extensions of our model to learn complex priors over these two domains separately. For example the dependence on context of the structure of odor presence and of the dynamics of concentration fluctuations due to air flow are likely to be only loosely correlated, given their disparate physical origins.

In closing, the approach we developed here extends to a broad array of constrained inference problems that arise in sensory modalities beyond olfaction. At its core, when inferring variables of different character—discrete and continuous, slow and fast—we can tailor the representation to each, and couple them through distinct computational pathways. Mirrored Langevin dynamics supplies the sampling substrate for this recipe, allowing us to build general recurrent circuits that solve constrained probabilistic inference problems. For example, inference under Poisson sensing noise occurs in many technological problems such as in low-light imaging, microscopy or astronomy [97], or in the growing field of event-based visual sensing [98]. Mirror descent—which is to optimization as MLD is to sampling—has been applied to study learning under biological constraints on synaptic weights [99]. However, the application of this toolkit to neural dynamics is, to our knowledge, novel. As MLD has recently gained popularity in machine learning for sampling from constrained distributions using diffusion generative models [100, 101], our approach holds much promise as a foundation for building circuit models that can rapidly process information in naturalistic and complex stimuli.

## Acknowledgements

We thank Dmitri Chklovskii, Farhad Pashakhanloo, and Sina Tootoonian for inspiring conversations. Moreover, we thank Juan Carlos Fernández del Castillo, Siddharth Jayakumar, and Farhad Pashakhanloo for helpful comments on a previous version of this manuscript.

C.P. is supported by an NSF CAREER Award (IIS-2239780), DARPA grants DIAL-FP-038 and AIQ-HR00112520041, the Simons Collaboration on the Physics of Learning and Neural Computation, and the William F. Milton Fund from Harvard University. This work has been made possible in part by a gift from the Chan Zuckerberg Initiative Foundation to establish the Kempner Institute for the Study of Natural and Artificial Intelligence. JAZV was supported by the Office of the Director of the NIH under Award DP5OD037354, and by a Junior Fellowship from the Harvard Society of Fellows. This research was carried out in part thanks to funding from the Canada First Research Excellence Fund, awarded to PM through the Healthy Brains, Healthy Lives initiative at McGill University. PM acknowledges the support of the Natural Sciences and Engineering Research Council of Canada (RGPIN-2025-05676 and DGECR-2025-00255), and of the Fonds de Recherche du Québec (CB-365865). PM was further supported by a Sloan Research Fellowship. This research was enabled in part by support provided by Calcul Québec and the Digital Research Alliance of Canada.

## Author contributions

PM and JAZ-V conceptualized this work, with input from VNM and CP. CJ and MYH contributed equally to coding, investigation, and figure preparation, with input from JAZ-V and PM. CJ, MYH, JAZ-V, and PM wrote an initial draft of the paper. All authors contributed to review and editing of the manuscript. JAZ-V and PM contributed equally to supervision.

## Data and code availability

Code to reproduce all figures is available on GitHub at: https://github.com/labmasset/Simultaneous-detection-and-estimation-in-olfactory-sensing.

## Supplemental Information

### A Notational conventions

We note that we will use vector notation, with ⊙ representing element-wise multiplication and ⊘ representing element-wise division, *i*.*e*., for two vectors **a** and **b** we write (**a** ⊙ **b**)_*i*_ = *a*_*i*_*b*_*i*_ and (**a** ⊘ **b**)_*i*_ = *a*_*i*_*/b*_*i*_.

### B Formulation of the problem

#### B.1 Modeling of the physical problem

We model the physical odor landscape with *n*_odor_ distinct types of odorants using a time series of vector pairs 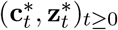, where 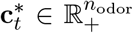 contains the concentration of each odorant and 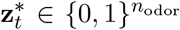 is a binary vector indicating which odorants are present, i.e. the odor identity.

This odor landscape is sensed by a population of *n*_OSN_ olfactory sensory neurons (OSNs). Each OSN is broadly tuned, responding to many different odorants with a sensitivity that varies from one odorant to another. We record these sensitivities in an affinity matrix 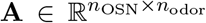, whose entry *A*_*ij*_ is the sensitivity of the *i*-th OSN to the *j*-th odorant. Through **A**, the OSNs’ activity reflects the combined contribution of all present odorants, and their firing rates form a time series (**s**_*t*_)_*t≥*0_. At each time *t*, the firing rate **s**_*t*_ is a sensory encoding of the underlying odor 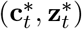, and is the only signal available to the downstream model. This encoding is stochastic, as OSN firing inherits the noisy nature of spike generation: for a given odor, **s**_*t*_ is governed by a probability distribution rather than a deterministic map, and the observed (**s**_*t*_)_*t≥*0_ is a single realization of this process — the only realization the model has access to at each time. We make the dependence of **s**_*t*_ on 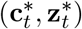 through **A** precise when we define the likelihood *p*(**s**_*t*_ | **c**_*t*_, **z**_*t*_) in Section B.2.

The problem of encoding 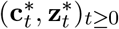 in (**s**_*t*_)_*t≥*0_ is called the forward problem in physics literature. Here, we focus on the other direction, i.e. inferring the physical information 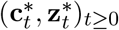 solely from the OSN activity (**s**_*t*_)_*t≥*0_. This is often referred to as the inverse problem. In particular, we frame this inverse problem in the Bayesian statistics framework.

#### B.2 Bayesian inference and posterior sampling

For a fixed *t* ≥ 0, we treat the physical truth 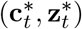 as the fixed unobserved target, while the realized observation **s**_*t*_ is the only information available to the model. To represent the physical world, our model carries its own internal latent random variables (**c**_*t*_, **z**_*t*_). The ultimate goal is to produce point estimates 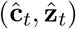 of the physical truth 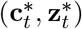 from the observation **s**_*t*_. Under a Bayesian perspective, all statistics of the truth 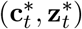 can be derived from the posterior *p*(**c**_*t*_, **z**_*t*_ **s**_*t*_).

We define the posterior starting from the joint distribution, which factorizes as

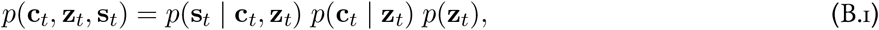

where the likelihood is

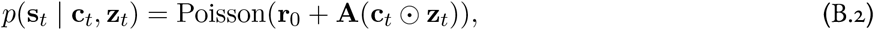

and the prior distributions *p*(**c**_*t*_ | **z**_*t*_) and *p*(**z**_*t*_) are specified later. Bayes’ rule then gives the posterior

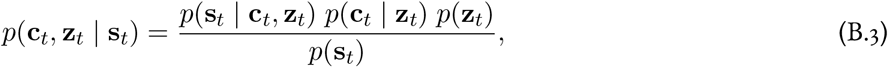

where *p*(**s**_*t*_) = ∫*p*(**s**_*t*_ |**c**_*t*_, **z**_*t*_) *p*(**c**_*t*_ |**z**_*t*_) *p*(**z**_*t*_) *d***c**_*t*_ *d***z**_*t*_ is the marginal likelihood. The exact *p*(**s**_*t*_) is intractable, due to the combinatorial sum involved in integrating over the binary **z**_*t*_, so the normalized posterior cannot be computed in closed form. Sampling the posterior, however, only requires it up to this constant, which bypasses the intractable *p*(**s**_*t*_) (further discussed in Section D). We therefore define the point estimators 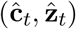 based on samples drawn from the posterior. With sufficiently many samples, these empirical estimators converge to the corresponding posterior quantities. Suppose we have *N* samples 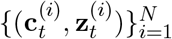 from (B.3). For the concentration vector, we take the point estimate to be the empirical meanof these samples,

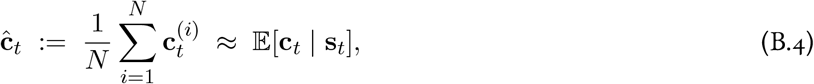

which approximates the posterior mean. For the binary identity vector 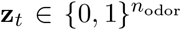, we likewise form the empirical mean of the samples,

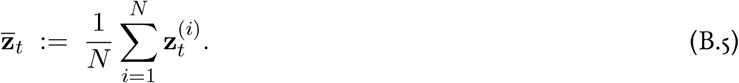

Because each sample 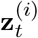 is binary, this average is simply the fraction of samples in which each odorant appears as present, and it approximates the marginal posterior probability of presence,

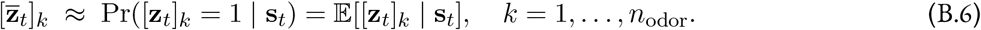

This marginal probability cannot itself serve as a binary estimate, since it does not lie in 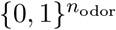. We therefore convert 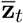 into a binary point estimate by thresholding componentwise at a level *p*_th_ ∈ (0, 1),

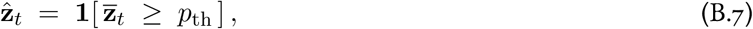

where the indicator is applied componentwise.

The choice of *p*_th_ implicitly specifies a cost structure on classification errors. Consider the asymmetric 0–1 loss for each component^1^

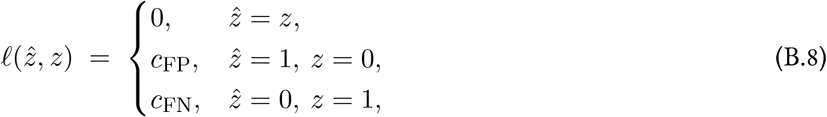

where *z* ∈ {0, 1} is the true label and 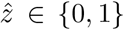 is the decision. The Bayes-optimal decision minimizes the posterior expected loss 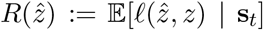, which sets 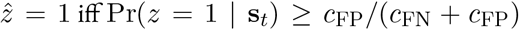. Matching this rule with our componentwise thresholding, we have that thresholding at *p*_th_ is the Bayes-optimal decision under the loss defined above, with a cost ratio

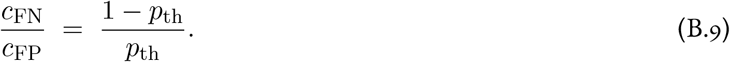

It follows that setting *p*_th_ = 1*/*2 corresponds to *c*_FN_ = *c*_FP_ and recovers the marginal maximum a posteriori (MAP) estimate of each [**z**_*t*_]_*k*_. Moreover, lowering *p*_th_ raises the implicit cost ratio, treating missed detections as more costly than false positive and thus trading precision for recall, which is appropriate when odorant presence is sparse. Unless stated otherwise, we use *p*_th_ = 0.2, corresponding to a cost ratio of 4:1.

Together, 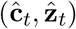 constitute the point estimate of the physical truth 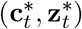 recovered from **s**_*t*_, and they are the outputs of our model.

#### B.3 The time-varying setting and online sampling

In the previous section, we considered the problem at a single fixed time *t*, with a single observation **s**_*t*_. In reality, however, the inference problem is time-varying: at each *t* a new observation **s**_*t*_ arrives, and the underlying truth 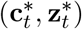 may have changed. The model must therefore produce a time series of estimates 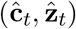 that reflects the changing odor landscape.

A central consideration here is the timescale separation between the sampling dynamics and the rate at which the OSN firing rate **s**_*t*_ changes. On one hand, one might assume that the sampling dynamics are much faster than the OSNs activity, in which case the algorithm can draw sufficiently many samples and reach equilibrium before the next observation arrives. On the other hand, one might assume that there is no such separation, so that for each observation the algorithm can draw only a single sample.

We adopt the latter assumption in our formulation. It is useful to distinguish two times: the *physical* (stimulus) time *t*, which indexes the observations **s**_*t*_ and the underlying truth 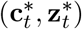, and the *sampling* (algorithmic) time *τ*, in which the Langevin/MLD dynamics derived below evolve — so every sampling stochastic differential equation (SDE) is written *d* **u**(*τ*), *d***c**(*τ*), … with Brownian motions **B**(*τ*). In principle the two are separate: one could run the *τ*-dynamics toward equilibrium for each fixed observation **s**_*t*_. Under the single-sample assumption, however, there is no separation — at each physical time *t* the sampler advances its *τ*-dynamics by exactly one step on the current observation **s**_*t*_, so *τ* and *t* advance in lockstep and we identify them in the deployed and simulated model. Namely, at each time *t*, the sampler draws one sample based on the observation **s**_*t*_, and iterates as time progresses. (B.4) and (B.5) in Section B.2 therefore collapse to the case *N* = 1, which implies the estimate at each *t* is just a single sample. In this case, the law of large numbers guarantee on convergence no longer applies. What compensates for the absence of averaging is the integration of information through time in the dynamical trajectory of the algorithm. From a high level, as time progresses, our sampler carries its state from the previous timestep through its dynamics instead of restarting. Therefore, information from observations is carried and integrated over time, even though each update is driven solely by the presented observation **s**_*t*_.

This setting emphasizes the core challenge we aim to address: detecting odors in a changing environment. It is the more demanding regime, yet remains compatible with the other: a model that works under the single-sample assumption should work at least as well when many samples are available, since the extra samples can only reduce the variance of the estimate instead of fundamentally changing the dynamics of the model. Additional details will be discussed when we introduce the design of our algorithm.

#### B.4 Conceptual overview

The following diagram summarizes our physical and statistical formulation of the olfactory sensing problem:

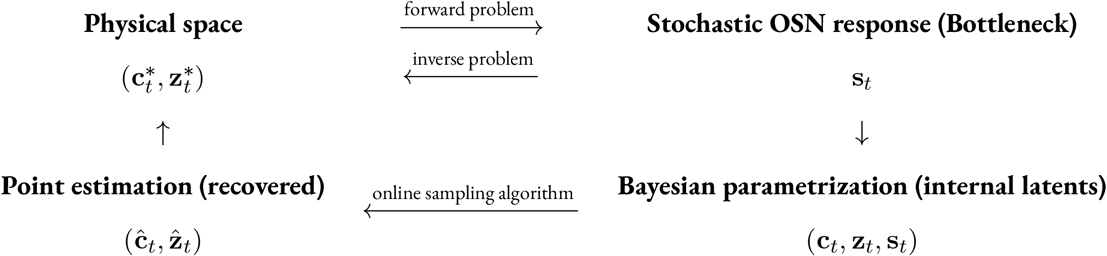

In the following sections, we derive our model as an online sampling algorithm for the time-varying posterior (B.3).

### C Review of Mirrored Langevin Dynamics

Our algorithm to sample the posterior (B.3) is built on the Mirrored Langevin Dynamics (MLD) framework introduced by Hsieh *et al*. [48]. Thus, before diving into the full derivation, we give a brief review of the MLD framework in this section. Our review presents a less general case of MLD that suffices for our application, as the formulation under the most general setting requires more sophisticated mathematical tools. In general, suppose that we wanted to sample a distribution *p*(**x**) over a random variable **x** ∈ℝ^*n*^ that is subject to some constraint, i.e., we have **x**∈ *X* for some convex set *X* ⊂ ℝ^*n*^ with a nonempty interior. Such a constraint prevents the direct application of standard Langevin dynamics. Instead, MLD prescribes that we can create an intermediate variable **y** in an unconstrained space, whose value and probability density are related to **x** in a particular way such that samples of **y** correspond to samples of **x** ~ *p*. Specifically, MLD can be established by first defining a smooth strictly convex function *ϕ* : *X* → ℝsuch that it transforms the constrained variable **x** into an unconstrained variable **y** ∈ ℝ^*n*^ via the “mirror map” defined by the gradient of *ϕ*, **∇**_**x**_*ϕ* : X → ℝ^*n*^:

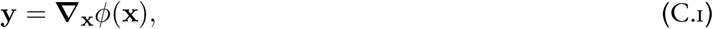

Following the convention in the optimization literature, we call **x** the primal variable and **y** the dual variable. The dual variable **y** can be mapped back to the primal variable **x** via an inverse mirror map, defined by the gradient of the Fenchel conjugate of *ϕ*, **∇**_**y**_*ϕ*^*\**^ : ℝ^*n*^ → *X*:

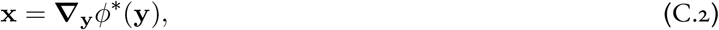

where the Fenchel conjugate *ϕ*^*\**^ is given by:

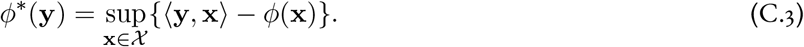

With appropriate choices of mirror map, this framework can handle many classes of constraints; the smoothness and strict convexity assumptions on *ϕ* can also be relaxed [48]. Since **y** is unconstrained, we can run naïve Langevin dynamics on **y**. Denoting the time in the dynamics by *τ*, we have:

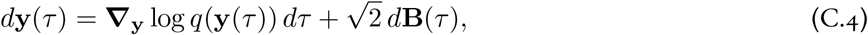

where **B**(*τ*) is a standard Brownian motion in the unconstrained dual space and *q*(**y**) is a probability distribution on **y** that we are free to choose. To build a valid MLD system, where samples of **y** actually correspond to samples from the target *p*(**x**), we must choose *q* such that if **y** ~ *q*, then **x**(**y**) = **∇**_**y**_*ϕ*^*\**^(**y**) ~*p*. Thus, the last step of establishing the MLD framework is to derive such *q*(**y**), which follows from applying the change-of-variables formula to the inverse mirror map **x**(**y**) = **∇**_**y**_*ϕ*^*\**^(**y**). By strict convexity of *ϕ*, the Jacobian of the mirror map **y** = **∇**_**x**_*ϕ*(**x**) is the Hessian of *ϕ*, which is positive definite:

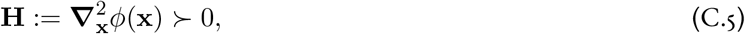

From the inverse function theorem, we have the Jacobian of the inverse mirror map **x**(**y**) is the inverse of the Jacobian of the mirror map:

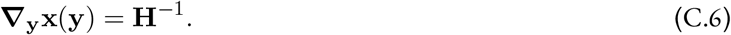

Applying the change-of-variables formula, the density of **y** corresponding to **x** ~ *p* is

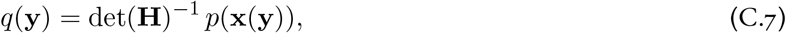

where the absolute value around the determinant is omitted since det **H** *>* 0. By the chain rule, the gradient (also called score) driving the Langevin dynamics in the unconstrained dual space ℝ^*n*^ is

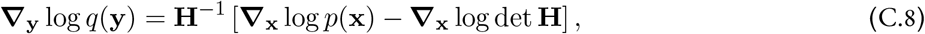

where the right-hand side is to be evaluated at **x** = **x**(**y**). Substituting (C.8) into (C.4) yields the mirrored Langevin dynamics:

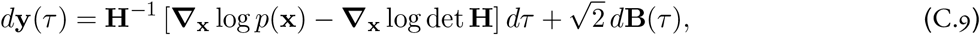

again with the right-hand side evaluated at **x** = **x**(**y**). To generate samples of the target *p*, one simulates this SDE for **y** and maps each sample back via the inverse mirror map **x**(**y**) = ***T***_**y**_*ϕ*^*\**^(**y**).

### D Derivation of the full model

Thus far, we have specified the inference target at the statistics level and reviewed the general MLD framework. We now put the two together and demonstrate the derivation of the full model. The remaining design choices fall into two levels: optimization and implementation. At the optimization level, we will address the technical difficulty of sampling the posterior *p*(**c**_*T*_, **z**_*T*_ | **s**_*T*_): through a deliberate continuous relaxation of **z**_*T*_ into a surrogate, followed by the application of the MLD framework, we derive a continuous-time dynamical system. We then discuss how such a dynamical system can be implemented: by introducing a few further modifications, we map the model onto a biologically plausible circuit and assign a biological interpretation to each component of the computation.

#### D.1 Sampling target and failure of vanilla Langevin dynamics

For a fixed *T*, we recall that our goal is to sample the Bayes posterior

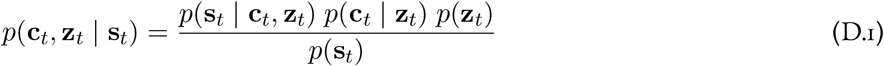

over odor presence 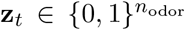 and concentration 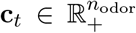 given a realized 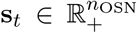 of olfactory sensory neuron (OSN) activity. Defining the potential of this posterior as

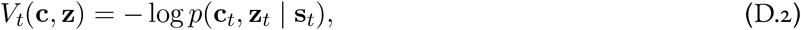

a Langevin-type sampler would run a diffusion driven by its gradient, of the form

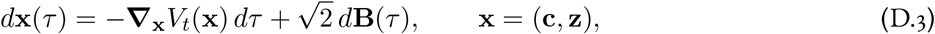

where the only state-dependent term is the gradient of the potential and **B**(*τ*) denotes a standard Brownian motion. We write (D.3) only for illustration: as it stands it is ill-defined, because both latent variables are constrained—**z** to the discrete set 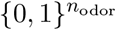 and **c** to the nonnegative orthant 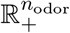 —and naive Langevin dynamics cannot be directly applied under such constraints. The core of the optimization-level design of the model is to turn (D.3) into a valid sampler. Nonetheless, since we have

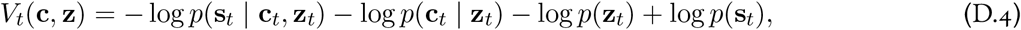

taking the gradient with respect to **c** and **z** eliminates the constant term log *p*(**s**_*T*_); the intractable marginal likelihood *p*(**s**_*T*_) therefore never enters the dynamics, as first noted in Section B. Moreover, considering **x** = (**c, z**) as a concatenation, (D.3) splits into

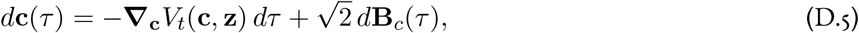

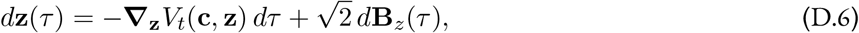

with independent Brownian motions **B**_*c*_ and **B**_*z*_, which lets us derive the inference dynamics for (**c, z**) separately.

Since **c** is continuous, (D.5) is already a valid Euclidean diffusion. The only issue is that it does not guarantee that **c** remains nonnegative. It turns out we can maintain positivity through the concentration prior rather than the sampler geometry, which will be discussed after we resolve the challenge of sampling **z**_*T*_.

(D.6) as it’s written has a bigger problem, since **z** is discrete, neither the gradient ***T*** _**z**_*V*_*T*_ nor a Brownian motion is defined. The immediate work to be done is to make it well-defined. As demonstrated in the next section, we achieve this by relaxing **z** into a continuous surrogate and applying the MLD framework discussed previously in Section C.

#### D.2 Continuous relaxation: presence probability as a surrogate

Recall from Section C that, for the MLD framework to be applicable, the variables must be constrained to a convex set with nonempty interior. Although the concentration **c**_*T*_ already meets this requirement, the presence **z**_*T*_ does not, as the binary set 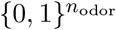 is neither convex nor has a nonempty interior. We therefore cannot apply MLD to **z**_*T*_ directly. A natural choice of relaxation for the binary constraint is its convex hull, the hypercube 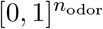. That is, we create a continuous variable 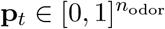 supported on the hypercube, as a surrogate to replace the binary **z**_*T*_.

Having fixed the domain, it remains to define how this surrogate relates to the binary **z**_*T*_. In particular, we want the surrogate to be chosen so that solving the relaxed problem also solves the original reasonably well. Among the candidates, we give **p**_*T*_ a definite meaning: each component [**p**_*T*_]_*k*_ is read as the estimated probability that the *k*-th odorant is present, a posterior quantity conditioned on the realized observation. In fact, it is the marginal of **z**_*T*_ given **s**_*T*_. Therefore, as an internal latent variable of the model,

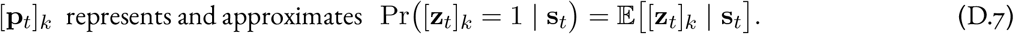

Partially substituting **p**_*T*_ for **z**_*T*_ in the likelihood and the presence prior of *V*_*T*_, we obtain the surrogate potential

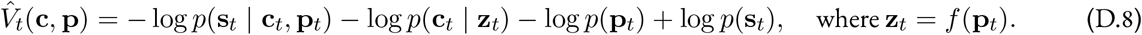

Under this surrogate, the dynamics is fundamentally about **c** and **p**, while we recover **z** through a defined relation **z**_*T*_ = *f* (**p**_*T*_). In our case, *f* is simply an elementwise gating at some threshold *p*_th_, carrying the same Bayes-decision semantics discussed in Section B for the thresholding of the estimator. We note that the concentration prior *p*(**c**_*T*_ | **z**_*T*_) is deliberately left unchanged, as in the original *V*_*T*_. From a high-level perspective, 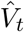 is a smooth interpolation of *V*_*T*_ over the hypercube 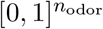, which gives us solid ground to establish the MLD framework and eventually derive a valid sampling dynamics.

Although the surrogate 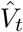 does not equal the original potential *V*_*T*_ and its design is heuristic, we can nonetheless rationalize the choice of surrogate by examining how far it deviates from the original and what semantics it carries.

##### Surrogate gap

First, the surrogate contains the original at the corners of the hypercube: when 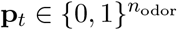, we have 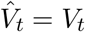, up to an additive constant from the prior normalization that does not affect the dynamics after taking the gradient.

Second, away from the corners, we can derive an approximate expression for the gap. The only state-dependent deviation between 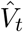 and the original is in the Poisson likelihood, since the remaining terms are inert: the concentration prior is unchanged and the marginal-likelihood term log *p*(**s**_*T*_) is constant. The surrogate evaluates the likelihood at the rate ***λ***(**p**) = **r**_0_ +**A**(**c**⊙ **p**), whereas the original uses ***λ***(**z**) = **r**_0_ +**A**(**c** ⊙**z**). Since the rate is affine in the presence and **p**_*T*_ = E[**z**_*T*_ **s**_*T*_], the surrogate rate is exactly the expected rate, ***λ***(**p**) = E_**z**_[***λ***|(**z**)]. Writing the log-likelihood (dropping the constant log **s**!) as

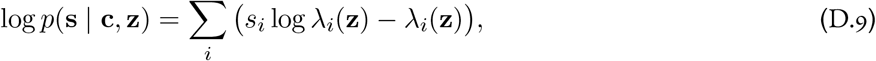

we compare the surrogate to the expected original by taking their difference, denoted *G*:

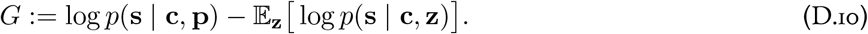

The linear term −*λ*_*i*_ has expectation −*λ*_*i*_(**p**) and cancels, leaving only the contribution of the logarithm,

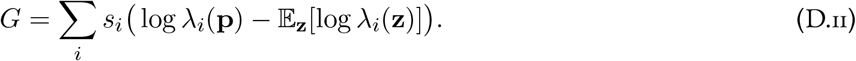

By the concavity of the logarithm (Jensen’s inequality), we have that *G* ≥ 0 is nonnegative. Semantically, this implies the surrogate always over-estimates the expected true log-likelihood: it is optimistic about how well a given (**c, p**) explains the data, relative to averaging over the binary configurations of **z**. On its own, however, this inequality is in the wrong direction: it provides a lower bound instead of an upper bound of the gap. Instead, we use a second-order expansion to estimate the gap and show how it depends on **p**:

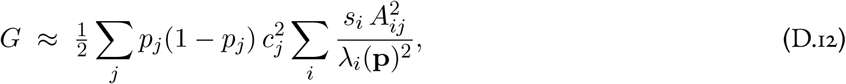

which is controlled by the Bernoulli variance *p*_*j*_(1 −*p*_*j*_) of each presence variable: it vanishes as *p*_*j*_ →0 or 1, and is largest at *p*_*j*_ = 1*/*2. The surrogate likelihood therefore agrees with the original at the corners and stays close whenever the presence estimate is confident—precisely the regime the dynamics are driven toward. Empirically, we observe in our simulations that **p** clusters near the endpoints 0 and 1 and crosses 1*/*2 rapidly, indicating that the surrogate operates almost always in a high-fidelity regime. Combining this analytical estimate with the empirical dynamics, we conclude that our choice of surrogate is numerically sound.

##### Semantic meaning

This surrogate also carries a clear semantic meaning; by construction, **p**_*T*_ is the marginal posterior probability of presence. Thus, this implies our sampling dynamics operates on a probabilistic representation of odor identity, and the binary identity is obtained as a readout through the relation **z**_*T*_ = *f* (**p**_*T*_). Such representation also inherently encodes the uncertainty, in other words, confidence of the predictions, which is not the case in a direct estimation of **z**_*T*_.

Given its semantic meaning, we also call **p**_*T*_ the presence estimate onward.

#### D.3 Deriving the Mirrored Langevin Dynamics

We now apply the MLD framework to derive the dynamics for the joint inference of presence and concentration, based on the surrogate potential 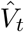 in (D.8).

##### Mirror map for presence p_*T*_

Our surrogate presence 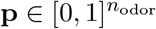 is constrained to a hypercube, to sample which we adopt the MLD framework introduced in Section C. The first step is to define a bijective mirror map between the presence **p** 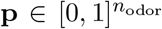 and an unconstrained variable 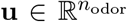. A natural smooth, bijective map between ℝ and [0, 1] is the logistic sigmoid, so we define the relation

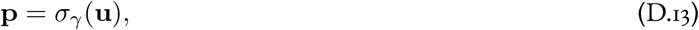

where for any 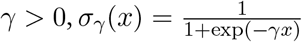 is the logistic sigmoid with gain *γ*, taken to act elementwise; the gain *γ* controls its steepness. To apply the MLD recipe, our goal is to construct a convex function 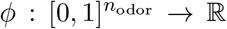 whose Fenchel conjugate 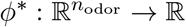 has gradient equal to this map, i.e.

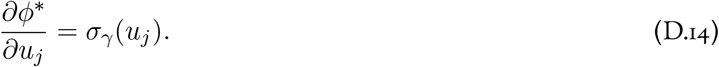

This implies that we should have

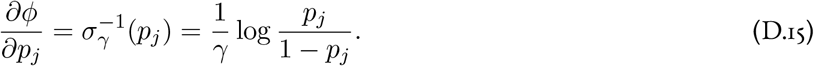

The natural choice—as discussed by Hsieh *et al*. [48]—is then to take

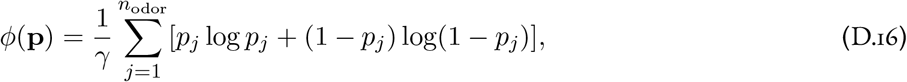

whose Fenchel conjugate, obtained from the definition (C.3), is

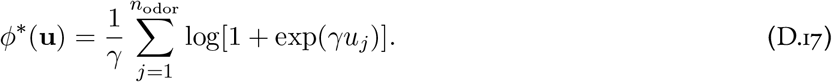

Here, we use the usual information-theoretic convention that 0 log 0 = 0.

Instantiating the general MLD dynamics (C.9) for the surrogate potential 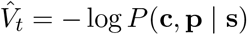, which we abbreviate 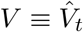 from here on, the dynamics for the presence are

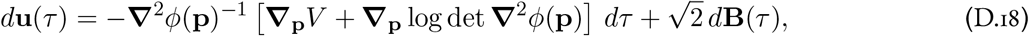

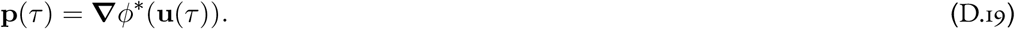

We now compute the inverse Hessian **∇**^2^*ϕ*(**p**)^*−*1^ and the term **∇**_**p**_ log det **∇**^2^*ϕ*(**p**) appearing in (D.18). As *ϕ* is additively separable, its Hessian is diagonal,

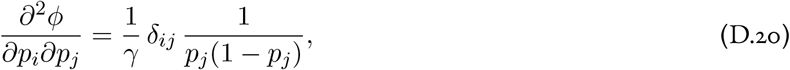

where *δ*_*ij*_ = 1 if *i* = *j* and 0 otherwise. Since the Hessian is diagonal, its determinant is the product of its diagonal entries, so the log-determinant is the sum of their logarithms,

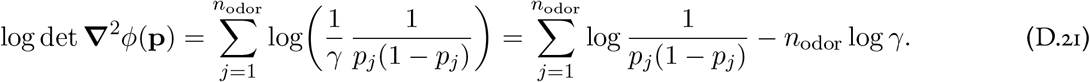

To differentiate with respect to a single coordinate *p*_*j*_, note that only the *j*-th term depends on *p*_*j*_, and that 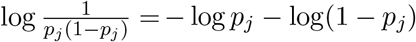. Hence

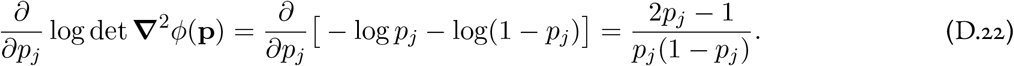

Because the Hessian is diagonal, its inverse is also diagonal, with entries

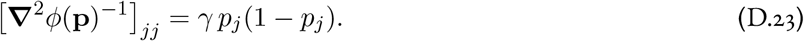

Substituting this inverse and the log-determinant gradient into (D.18) and reading off the *j*-th coordinate,

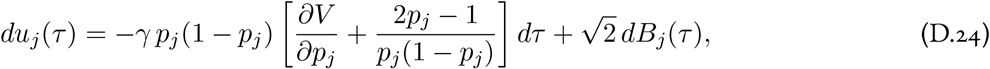

in which the factor *p*_*j*_(1 − *p*_*j*_) cancels against the denominator of the second term, leaving

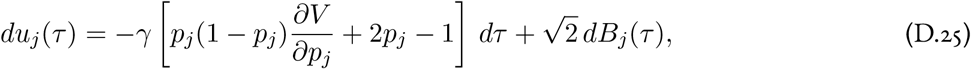

or, in vector form,

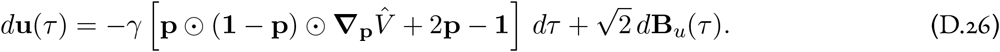

Meanwhile, we can directly use standard Langevin dynamics to sample **c**,

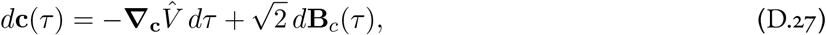

for an independent Brownian motion **B**_*c*_. To be fully formal, we can also introduce a trivial identity mirror map for **c**, which casts the joint sampling problem within the MLD framework under a combined mirror map induced by

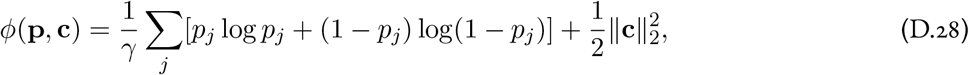

which handles the constraint on **p** while leaving **c** intact. Although **c** is restricted to the nonnegative orthant 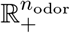, the trivial mirror map leaves it to ordinary Euclidean Langevin dynamics, which does not by itself keep **c** positive. We do not enforce positivity through the mirror geometry; instead, it is maintained by the concentration prior. For the Gamma prior we adopt below (with shape *α >* 1), the score **∇**_**c**_ log *P* (**c** | **p**) = (*α* − 1)**1** ⊘ **c** − *β***1** diverges to +∞ as any component *c*_*i*_ → 0^+^, acting as a logarithmic barrier that repels the dynamics from the boundary. This soft barrier suffices in continuous time; numerically, we additionally floor **c** at a small *ε >* 0 to guarantee positivity under finite step sizes.

We remark in passing that not all of the distributions of interest are log-concave in the dual space (which is the relevant notion for MLD [48]). Guarantees for the convergence of stochastic gradient Langevin dynamics given certain sufficient conditions when sampling on non-log-concave target distributions are weaker in general than those known in the log-concave case, but theoretical progress has in recent years been rapid [102, 103]. Since we are anyway relying on numerics rather than rigorous proofs for fast convergence, we will not dwell on this issue further.

In summary, applying the MLD framework to the surrogate potential 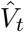 yields a continuous-time dynamical system that jointly samples concentration and presence. The presence is handled in the dual space: an unconstrained variable **u** evolves under the mirrored dynamics (D.26), and the presence is read out as **p** = *σ*_*γ*_(**u**). The concentration **c** evolves directly under ordinary Langevin dynamics (D.27). Both are driven by the gradients of the surrogate potential, 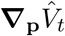 and 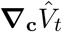, and their stationary distribution is the surrogate posterior over (**c, p**). It remains only to specify these gradients, which we expand into a likelihood and prior terms in the next section and complete by fixing the priors thereafter.

#### D.4 Simultaneous inference of concentration and presence identity: Coupling mechanism and readout

Our model infers the concentration **c** and the binary presence identity **z** of odors simultaneously by running the two dynamics in (D.26) and (D.27) together. Here, we address how these two dynamics are coupled and how they produce the estimation readout.

Firstly, although the presence dynamics evolve the dual variable **u**, we can consider them as equivalent in terms of **p**, since **p** = *σ*_*γ*_(**u**) is a bijection and the two carry the same information.

The dynamics of **c** and **u** are then linked through a reciprocal coupling. Specifically, **c** and **u** enter each other’s dynamics through the gradient terms _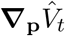_ and _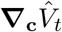_, which will be shown once we derive their exact expressions.

It follows that the two outputs **ĉ** and 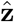 are produced as readouts from the system.

For the output 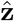, we simply let it be the iteratively updated readout **z**_*T*_ := *f* (**p**_*T*_) as in (D.8), where *f* is the threshold gating mechanism. The output **ĉ**, however, requires a more deliberate choice. In our specific setting, the internal representation of **c** is not directly hard gated by **z**, which means **c** can be positive when 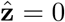. To avoid this and produce a physically meaningful estimation of concentration at the output level, we must therefore further hard gate the concentration estimate by the binary identity of presence, namely **ĉ** = **c** ⊙ **z**. This ensures that concentration estimations become 0 whenever the odor is not detected, while allowing the model to still possess a continuous internal representation of concentration for running the Langevin dynamics.

In fact, running estimation on a continuous internal **c** and masking afterward is necessary. We could not instead directly sample a gated quantity such as **c** ⊙ **z**, because the Langevin dynamics are not well-defined when the concentration is suppressed to zero through a gating mechanism.

We summarize the coupling and the readout in the diagram below.

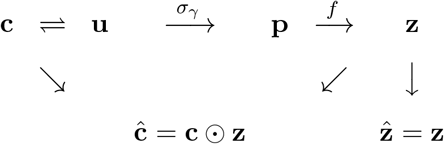

##### Plotting convention

For empirical results, we plot **p**_*T*_ instead of **z**_*T*_ because it demonstrates richer dynamics, while the reader can still easily infer **z**_*T*_ given the gating threshold plotted. For a similar reason, we also plot **c**_*T*_ to show more detailed dynamics.

#### D.5 Gradients of the posterior

We now explicitly compute the gradients of the surrogate potential 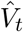 that drive the MLD. Recall that we have,

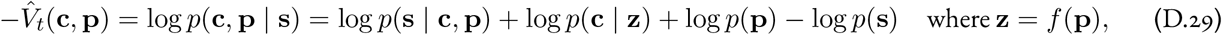

where *f* is an elementwise threshold. Thus, the required gradients are

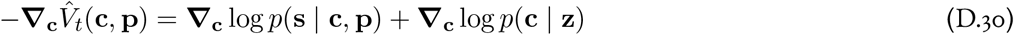

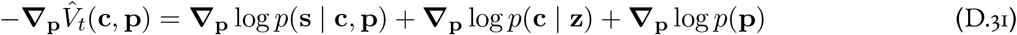

where the constant log *p*(**s**) drops out under the gradient. For a Poisson likelihood with rate 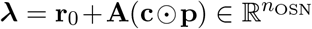,we have:

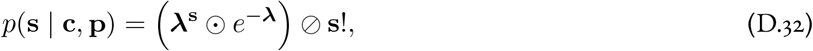

where ***λ***^**s**^ and *e*^*−****λ***^ are elementwise operations. Notably, although the Poisson probability mass function (PMF) is discontinuous with respect to **s**, it is continuous and hence differentiable with respect to **c** and **p**. Taking logarithms, we have

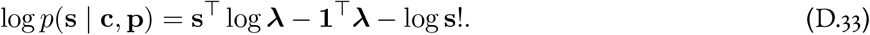

We let **h**_*T*_ := **s**⊘ ***λ*** = **s** ⊘ (**r**_0_ + **A**(**c**⊙ **p**)) and interpret it as the activity of one specific population of neurons. Empirically, we found the behavior of this population is similar to that of tufted cells; hence we label them as tufted-like cells with subscript *T*.

Mathematically, **h**_*T*_ is the ratio of the observed OSN rate to the rate predicted by the current estimate. Because the rate ***λ*** = **r**_0_ + **A**(**c** ⊙ **p**) is gated by the soft presence **p, h**_*T*_ encodes the soft-gated estimation error. It follows that 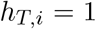 signals a perfect match and **h**_*T*_ − **1** is the relative prediction error.

Since 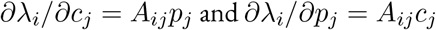, by the chain rule we have

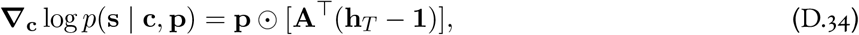

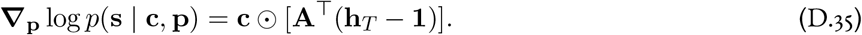

Substituting (D.34) and (D.35) back into (D.30) and (D.31), we have

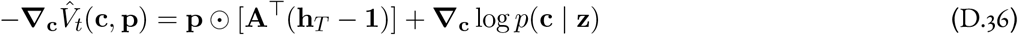

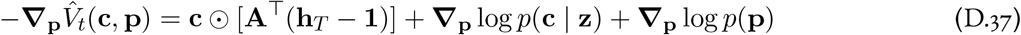

We leave the prior terms unspecified, as we will subsequently explore different choices of prior.

#### D.6 Choices of prior discussion: spike and slab prior

There are two prior distributions in the potential 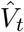 in (D.8): *p*(**c** |**z**) and *p*(**p**). Both of them directly show up later in (D.36) and (D.37).

While the prior on **p** does not raise this question, there is more physical nuance when choosing the prior *p*(**c**| **z**). Specifically, whether we should let this prior depend on **z**.

At the level of the physical model, the natural choice is a spike-and-slab prior on the presence-masked concentration [**c**_*T*_]_*i*_[**z**_*T*_]_*i*_: a point mass at zero (the spike) when the odorant is absent, [**z**_*T*_]_*i*_ = 0, and a Gamma density (the slab) when it is present, [**z**_*T*_]_*i*_ = 1. This is the most realistic choice, as it aligns with the sparse nature of the physical odor landscape, where most odorants are absent with few being present. On the masked quantity, this structure is clean: masking alone produces the spike, since [**c**_*T*_]_*i*_[**z**_*T*_]_*i*_ = 0 whenever [**z**_*T*_]_*i*_ = 0, regardless of [**c**_*T*_]_*i*_.

The difficulty is that our sampler does not operate on the masked quantity [**c**_*T*_]_*i*_[**z**_*T*_]_*i*_, but on the internal continuous concentration **c**, and on this variable the spike-and-slab is problematic for two reasons. First, the point mass at zero is not differentiable, so we cannot run Langevin dynamics on a concentration drawn from it. Second, even setting this aside, it lacks biophysical plausibility, as it is unclear how it could be implemented in biological neurons.

We therefore separate the two levels. In the following sections, we first derive our model using a presence-independent Gamma prior on the internal **c**—differentiable and benign for the dynamics—and then restore the spike-and-slab structure at the level of the computation, through the introduction of a hard-gated (mitral-like) neuron population. In effect, the spike-and-slab that is natural on the masked concentration is recovered not as a prior but as a gating operation in the algorithm.

#### D.7 Model equations for a continuous Bernoulli prior on presence and a Gamma prior on concentration (Softgated-only)

We now derive the version of our model that uses a continuous Bernoulli prior on presence and a Gamma prior on concentration. We assume that different odorants are independent and identically distributed under the prior. This could be relaxed, but would require more notation.

We first define a continuous Bernoulli prior distribution for the presence **p**, namely the *p*(**p**) term in (D.8). Element-wise, we have:

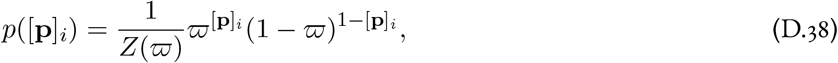

where *ϖ* ∈ (0, 1) is a parameter, and *Z*(***ϖ***) is a normalization constant with:

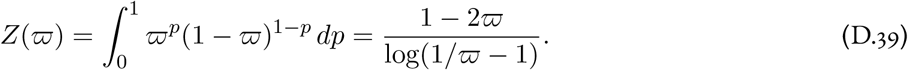

However, as only the score 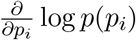 appears in the dynamics, showing this normalization constant is mainly for formal rigor.

For the prior on concentration, namely the *p*(**c**_*T*_| **z**_*T*_) term in (D.8), we take the straightforward route flagged in the prior discussion above and choose [**c**]_*i*_ | [**z**]_*i*_ to be the same regardless of the value of [**z**]_*i*_. This presence-independent choice is the most straightforward way to sidestep the non-differentiability and biophysical implementation issues of a true spike-and-slab prior, at the cost of leaving the latent **c** ungated internally; we restore the spike-and-slab structure in the next section. The same unified choice was also made by Grabska-Barwińska *et al*. [29]. Following those authors, we use a Gamma distribution with parameters *α* and *β*:

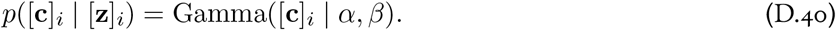

Now we consider the two components of the prior. Clearly, ***T***_**c**_ log *p*(**p**) = **0**, while

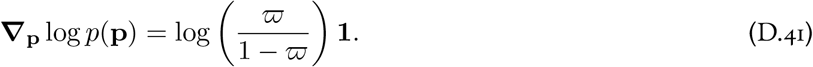

The log-prior on concentration is

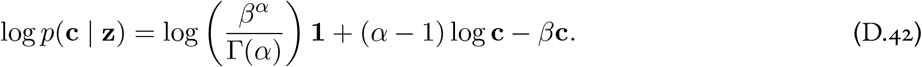

Because of our simplifying choice that the concentration prior does not depend on presence, we have

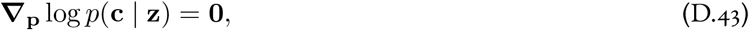

while the derivative with respect to **c** yields

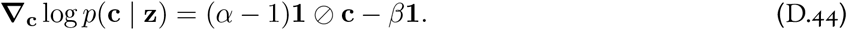

Substituting (D.41), (D.43) and (D.44) into (D.36) and (D.37), we then have

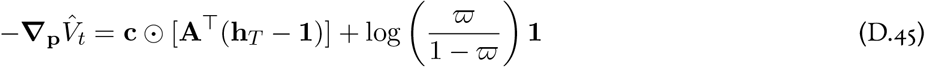

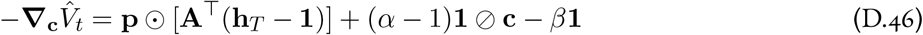

Substituting (D.45) and (D.46) into the mirrored Langevin dynamics (D.26) and (D.27) gives the soft-gated-only model.

Before we lay out the explicit equations for the soft-gated-only dynamical system, we first group a few quantities by their semantic meaning. We write ***λ*** = **r**_0_ + **A**(**c** ⊙ **p**) for the OSN rate predicted under the soft presence **p**. Recall that **h**_*T*_ := **s** ⊘ ***λ***, we can define ***ε*** as the corresponding normalized OSN prediction error:

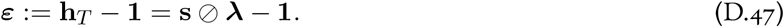

We have that ***ε*** is zero when the prediction matches the data. It follows that the error ***ε*** backprojected into odorant space is:

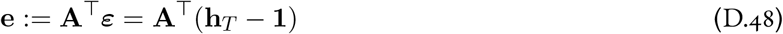

which is in fact the bottom-up evidence on each odorant.

Letting 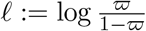, we have log **∇**_*p*_(**p**) = *l* **1**, and the dynamics of the soft-gated version of the model become,

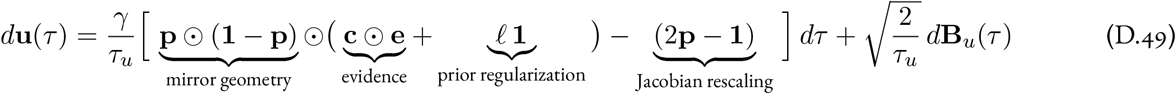

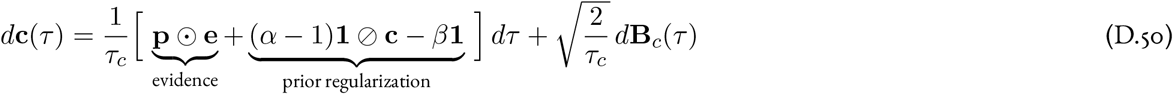

Such high-level formulation allow us to observe the structure of the dynamics. In the dynamics for **c** and **u**, the evidence term pulls the estimate toward explaining the OSN response, a prior term, coming from the gradient of the prior, pulls the estimate toward the high density value in the prior and hence serves as regularization.

Due to the dynamics of **u** involving a mirror map, we have two additional terms: the gain rescaling **p** ⊙ (**1**− **p**) shaped by the mirror map, which vanishes at the boundary {0, 1}, and the Jacobian rescaling 2**p 1** coming from the determinant of Jacobian in the change-of-variable formula. The Jacobian rescaling shows up as an additive term because we are operating on the log-density.

With **p** = *σ*_*γ*_(**u**), the model then outputs the estimator 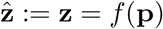 and **ĉ** = **c**⊙**z**, where *f* is the component-wise threshold at *p*_th_: *z*_*i*_ = 1 when *p*_*i*_ ≥*p*_th_ and *z*_*i*_ = 0 otherwise, encoded by 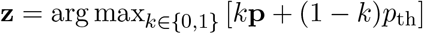.

Moreover, we now have explicitly shown that the dynamics of **c** and **u** coupled together through gating and rescaling on the evidence term. Specifically, in the presence estimation, the evidence is rescaled by the concentration, while in the concentration estimation, the evidence is gated by the presence.

Now, in detailed form, we expand the evidence down to the cell activity **h**_*T*_, the same dynamics then read,

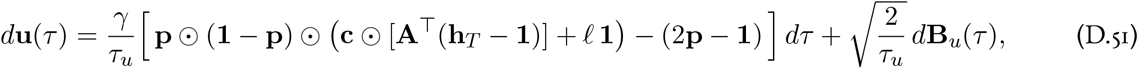

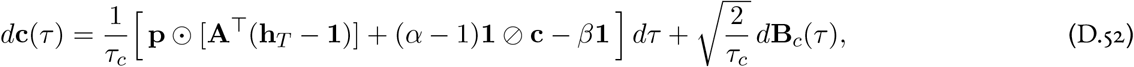

#### D.8 Introducing the hard-gated Mitral-like cell

In Section D.7, instead of using a spike-and-slab prior on concentration, we used a presence-independent Gamma; as a way to bypass the non-differentiable spike that makes running Langevin dynamics impossible. However, such a presence-independent prior does not realistically reflect the sparse nature of the physical odor landscape and consequently harms the inference performance.

Given that a direct implementation of the spike-and-slab prior is intractable, we restore its spirit through a special design at the level of the computation: the introduction of hard-gated cells.

We first introduce another surrogate potential in addition to 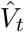 in (D.8):

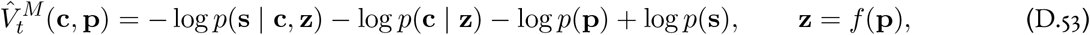

The difference between 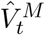 and 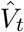 is in the likelihood term. While 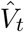 uses a surrogate likelihood 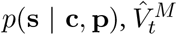 incor-porate a reconstructed likelihood *p*(**s** | **c, z**):

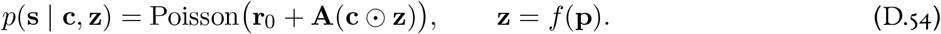

The reconstruction is in the sense that, the likelihood has the exact same form as the physical true likelihood, supported on binary **z** instead of a surrogate, and how we compute it is through a reconstruction of physical identity from **z** = *f* (**p**). Thus, while the main deviation from the exact potential *V*_*T*_ and 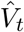 comes from the surrogate gap, the deviation for 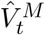 appears as a reconstruction loss.

We then derive our concentration dynamics using potential 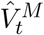 in (D.53) while keeping the presence dynamics derived from 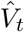 in (D.8).

For the gradient, since *∂λ*_*i*_*/∂c*_*j*_ = *A*_*ij*_*z*_*j*_ under (D.54), we have:

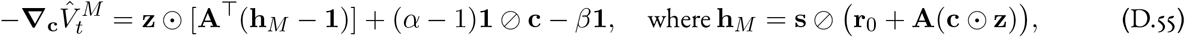

Here we interpret **h**_*M*_ as a new population of neurons. Empirically we find their activities are similar to mitral cells, and hence we denote it by subscript *M*. Mathematically, it encodes the prediction-error ratio under the reconstructed binary odor.

Plugging this gradient into (D.27), we have:

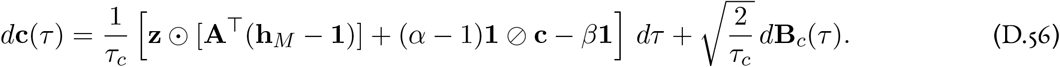

Combining this with the unchanged presence update from (D.49), we have the system:

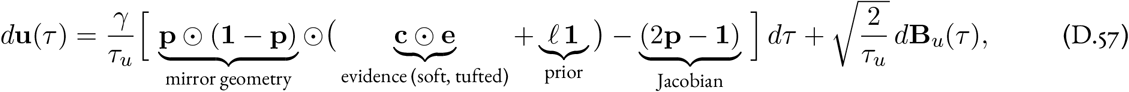

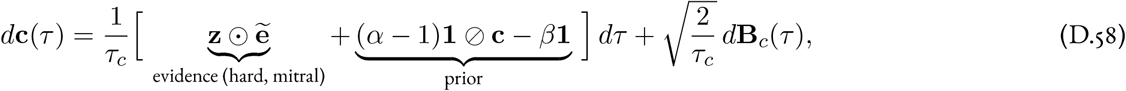

with soft evidence **e** = **A**^*T*^(**h**_*T*_ − **1**) and hard evidence 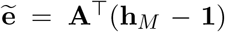. Expanding both evidences down to the cell activities gives the detailed form

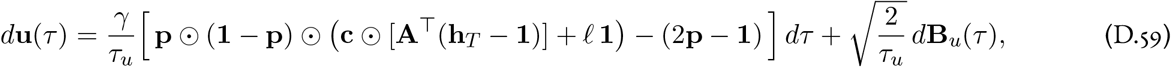

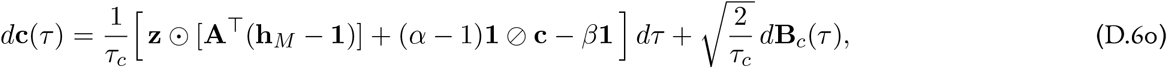

with the soft (tufted) and hard (mitral) cell activities

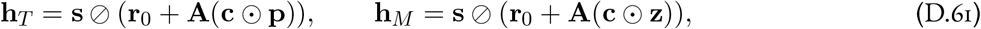

and **p** = σ_γ_(**u**), **z** = *f* (p)

In summary, the presence update is driven by the soft-gated tufted-like cells and the concentration update is driven by the hard-gated mitral-like cells. They correspond to a surrogate probabilistic and a reconstructed representation respectively.

Moreover, the coupling and readout now form a closed loop. Unlike the soft-gated-only model of Section D.7, where the concentration update was driven by the soft presence **p**, the concentration dynamics here are gated by the reconstructed binary readout **z** = *f* (**p**): the readout feeds back into the dynamics through the mitral channel **h**_*M*_, closing the cycle **c** → **u** → **p** → **z** → **c**.

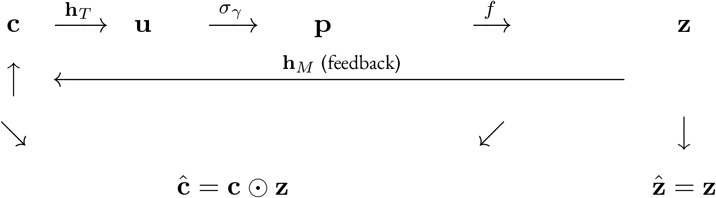

In Section D.9 we discuss how this two-channel structure can be implemented at the circuit level. Before that, we provide some additional discussion of the hard-gated dynamics.

As noted in the main text, performing concentration inference on a binary representation aligns better with the physical nature of the problem. The concentration of an odorant is a property of the odorant itself and should not be modulated by our confidence in its presence; the model should instead commit to which odorants are present and actively infer concentration only for those. This is also what the generative process dictates: an odorant reaches the OSNs only when it is actually present, so the rate that generated **s** is gated by the binary identity **z**, not the soft surrogate **p**. The soft presence is the right substrate for inferring presence, since it is differentiable and can be sampled, but for concentration the binary representation is the appropriate one. We therefore read out the binary representation by hard gating the soft presence.

This hard gating recovers the spike-and-slab structure at the level of the computation rather than the prior. The latent **c** still evolves under the smooth Gamma prior, but the masked output **ĉ** = **c** ⊙ **z** carries the structure: for an inferred-absent odorant (*z*_*i*_ = 0) the hard evidence vanishes and the mask reports exactly zero, reproducing the spike, while for an inferred-present odorant (*z*_*i*_ = 1) the mitral-like error **h**_*M*_ drives *c*_*i*_ toward the Gamma slab.

Numerically, this pathway is vital for stabilizing the system: an ablation that drops the hard-gated pathway fails when the true concentration falls below the prior expectation (Fig. S4). This may be because the hard-gating pathway makes concentration inference a better-posed subproblem: the dynamics run on a reconstructed support that is sparse, with most elements suppressed to exactly zero. Without hard gating, the evidence for absent odorants is only softly suppressed by their small presence probabilities, so across the large repertoire substantial noise leaks into the concentration dynamics. Effectively, the reconstructed representation constrains the sampler to an active subspace within the high-dimensional ambient space.

This also explains why the failure appears specifically at small concentrations. The Gamma prior contributes a repulsive term (*α* − 1)*/c*_*i*_ that pushes each *c*_*i*_ up toward the prior mode; only full-strength, support-conditioned evidence can overcome this barrier and pull *c*_*i*_ down to a genuinely small value. The diluted soft evidence cannot, so without hard gating the estimate is trapped near the prior expectation and systematically inflates true low concentrations.

#### D.9 Circuit implementation of the model

To implement the coupled dynamics in a biologically plausible way, we linearize their nonlinear terms by introducing auxiliary neural populations whose fixed points compute those terms, following Chalk *et al*. [58] and our previous work [39].

Here, we overload the notation: we let **h**_*T*_ and **h**_*M*_ in previous section become 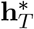 and 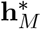, and let **h**_*T*_ and **h**_*M*_ be the activity of neuron that evolve around 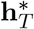 and 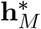 as fixed point.

Thus, we first have **h**_*T*_ in (D.59) becomes

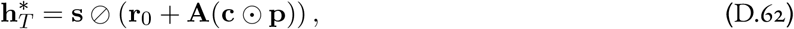

and create a relaxation dynamics that evolve around 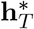 as fixed point:

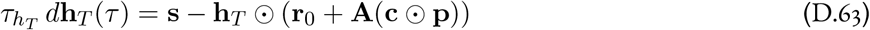

Similarly, we have **h**_*M*_ in (D.60) becomes

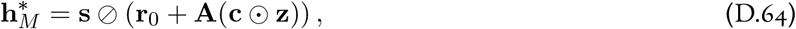

and we create the relaxation dynamics:

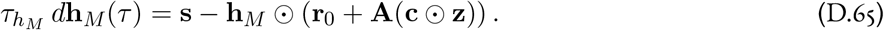

Finally, the Gamma prior contributes the term (*α* − 1)**1** ⊘ **c** to the concentration dynamics, we let this be a fixed point

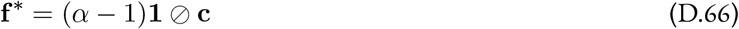

and create the relaxation dynamics **f** around it:

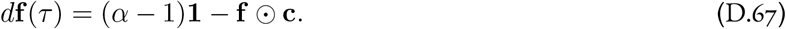

Putting everything together, the full dynamics are

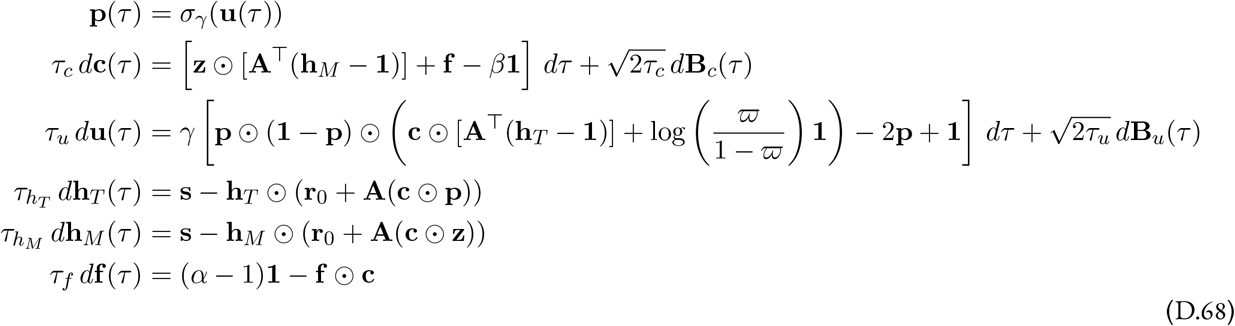

We can map these dynamics onto the circuit architecture of the olfactory bulb [7, 25]. Excited by the OSN input, the two populations **h**_*M*_ and **h**_*T*_ resemble the two classes of projection neurons in the OB, the mitral and tufted cells respectively. The concentration estimate **c** and presence estimate **p** are carried by local interneurons (granule cells), which inhibit the projection neurons and gate one another’s dynamics. The feedback population **f**, which linearizes the prior, can then be interpreted as a form of cortical feedback onto the granule cells.

### E Kumaraswamy Prior on Presence

In our olfactory sensing model, a prior distribution that reflects the underlying structure of the natural odorants landscape yields a more informative posterior and is expected to promote more efficient inference. Earlier, we used a unimodal continuous Bernoulli (CB) distribution as the prior on presence. However, the nature of the presence variable is bimodal, where *p*_*i*_ tends to be either 0 or 1. Concretely, the natural density of *p*_*i*_ should not be monotonically decreasing near the upper boundary 1, whereas in a Bernoulli prior the density taper off as *p*_*i*_ → 1 (Figure S11a). Therefore, compared to a unimodal prior, a bimodal prior is more realistic, and this motivates us to adopt a bimodal prior distribution. A commonly used bimodal distribution is the Beta distribution with a correct choice of parameters. However, practically it is hard to adopt the Beta distribution because its density and cumulative distribution function involve non-elementary functions. Fortunately, there is a Beta-type distribution that is easier to work with: the Kumaraswamy (KS) distribution, originally proposed by Kumaraswamy in 1980 [69]. **The original KS distribution**. A random variable *X* has the KS distribution if *X* has density function

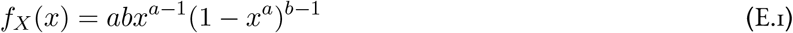

and cumulative distribution function (CDF)

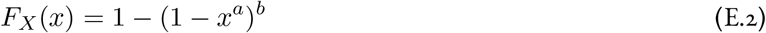

where *a, b >* 0 are shape parameters. The KS distribution, similar to the Beta distribution, can be bimodal, when *a <* 1 and *b <* 1, as visualized in Figure S11a. On the other hand, the KS distribution is also much easier to work with than the Beta distribution for several reasons. One of them is that it only involves simple functions in its density function [68]. **Truncation of the KS distribution**. The support of the original KS distribution is (0, 1); our presence variable *p*_*i*_ of each odorant, however, is defined on [0, 1]. Hence, we must transform variables to ensure that the support of the prior distribution matches the domain of *p*_*i*_. To eliminate the asymptotic behavior of the KS distribution near 0 and 1, we first restrict the support of KS distribution to the close interval *C* = [0 + *ϵ*, 1 − *ϵ*] for a small *ϵ >* 0.

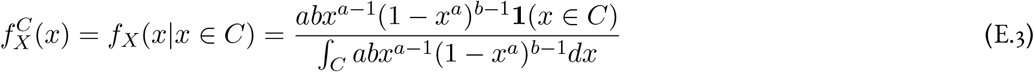

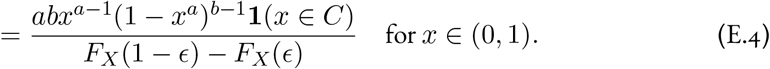

The restricted KS distribution has compact (closed and bounded) support [0 + *ϵ*, 1 − *ϵ*] and Lipschitz smooth density function. We then remap the restricted KS distribution to the full interval [0, 1]. We define a function 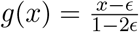 where *ϵ* ∈ (0, 0.5). Given *X* is a random variable with density function 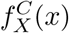, we define a new random variable *Y* as *Y* = *g*(*X*). Since the function *g* is injective, differentiable and has positive derivative, we can use the change of variables formula to get the probability density function of 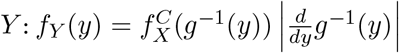. Since

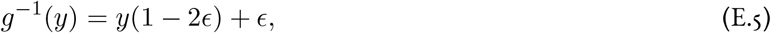

we have 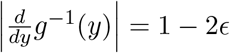, and so

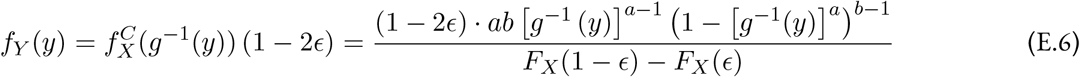

where *y* [0, 1]. Denoting the normalization factor 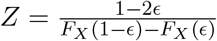, we have:

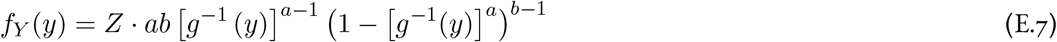

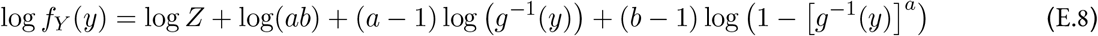

The density function *f*_*Y*_ (*y*) defines a new probability distribution that is well-defined on [0, 1]. and we call this distribution the transformed-KS (TKS) distribution. It follows that if we let *ξ* = *g*^*−*1^(*y*) = *y*(1−2*ϵ*)+*ϵ*, the first and second derivative of the log distribution (E.8) are:

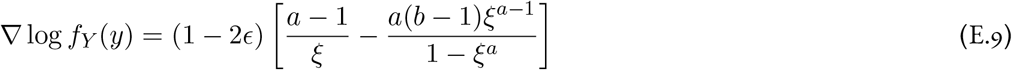

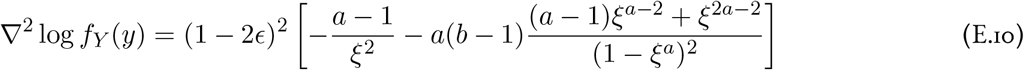

The first and second derivative of the log distribution are visualized in the third and fourth columns (left-to-right) in Figure S11a. We now choose a particular TKS distribution with the set of parameters *a, b* and *ϵ* to be the prior on presence. As before, we denote the prior distribution density function as *p*(**p**) in vector form. The gradient of the log-prior with respect to **c** is 0, while the derivative with respect to **p** is

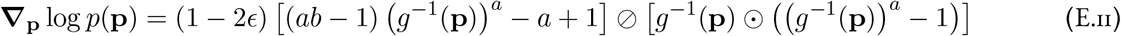

where *g*(**p**) = (**p** − *ϵ*) ⊘ (1 2*ϵ*) is the remapping function. The above equation (E.11) directly follows from (E.8). Using (E.9), the gradient of the energy function with respect to **p** is

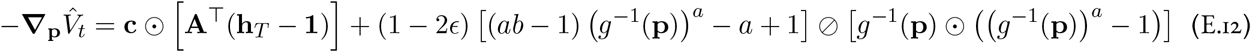

while the gradient with respect to **c** is unchanged:

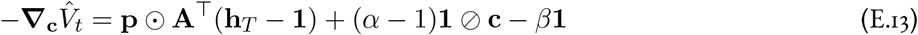

Thus, exactly as in the basic model (Section D.7), we hard-gate the concentration update through **z** = *f* (**p**) while the presence update keeps the soft tufted error; only the presence prior changes. In evidence-wrapped form the full dynamics are

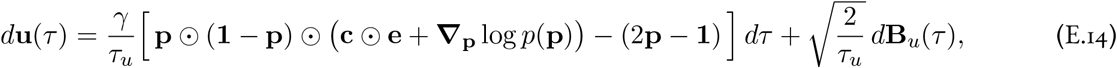

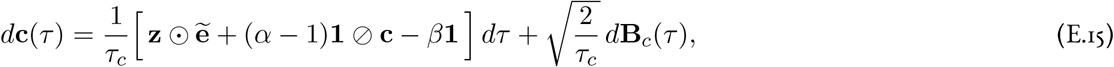

where **e** = **A**^*T*^(**h** − ^*T*^**1**) is the soft (tufted) evidence driving presence, 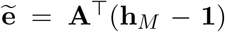 is the hard (mitral) evidence driving concentration, and **∇**_**p**_ log *p*(**p**) is the truncated-Kumaraswamy log-prior gradient (E.11). Expanding both evidences down to the cell activities gives the detailed form

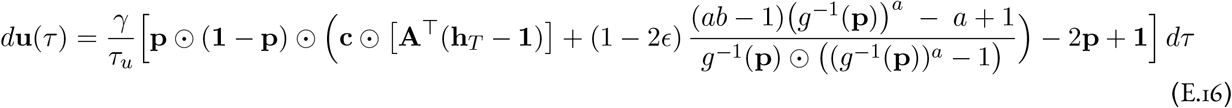

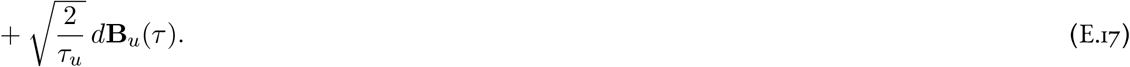

and

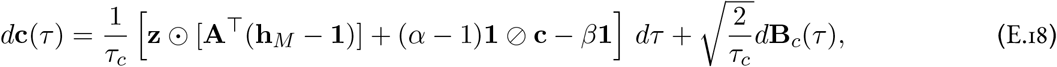

with the soft (tufted) and hard (mitral) cell activities **h**_*T*_ = **s**⊘ (**r**_0_ + **A**(**c**⊙ **p**)) and **h**_*M*_ = **s**⊘ (**r**_0_ + **A**(**c**⊙ **z**)). The gating structure is identical to the basic model of Section D.7 — soft tufted error for presence, hard mitral error for concentration — with only the presence prior replaced by the truncated Kumaraswamy. We now try to gain some more intuition for the behavior of the prior over **u** resulting from the truncated KS prior over **p**. Importantly, it is clear that it is not log-concave. As the prior is factorized over odorants, consider the *j*-th odorant, for which we have:

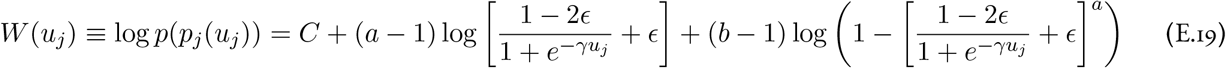

where *C* is a normalizing constant. We can also derive *W*^*′*^(*u*_*j*_) and *W*^*′′*^(*u*_*j*_) as following:

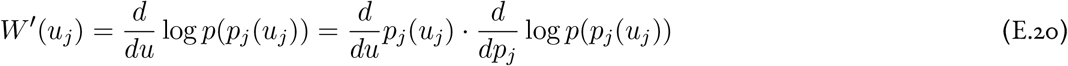

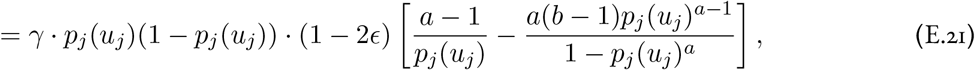

where we get 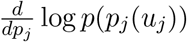 from (E.9).

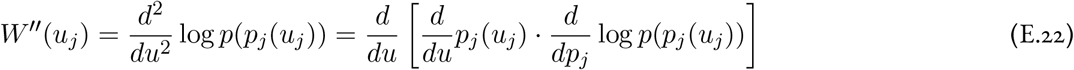

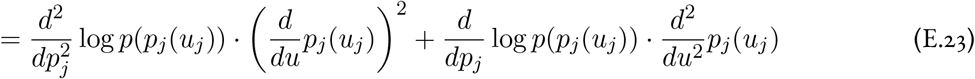

We can then substitute (E.9) and (E.10) in and get the second derivative w.r.t. *u*_*j*_. For any positive small 1 ≫*ϵ >* 0, *W* (*u*_*j*_) is roughly constant for any *u*_*j*_ of even modestly large absolute value, with a transition region between. In particular, for *u*_*j*_ → −∞ we have

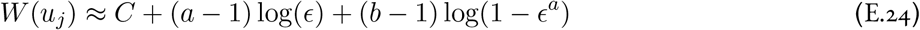

while for *u*_*j*_ → ∞ we have

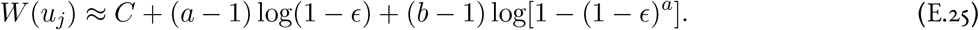

In between, we note that

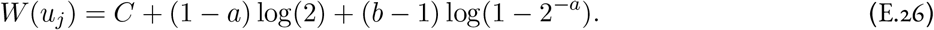

What remains is to figure out the behavior of the function in the transition region, as well as the width of that region. By direct computation, we find that the only stationary point 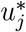 of *W* (*u*_*j*_) (that is, the solution to 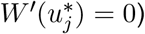 is

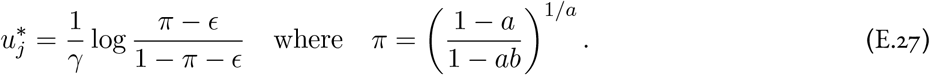

Using Mathematica, we can verify that the Hessian 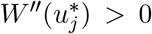 at this point, meaning that it is a local minimum. Moreover, we can see that *W*^*′*^(*u*_*j*_) *>* 0 for 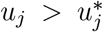, while *W*^*′*^(*u*_*j*_) *<* 0 for 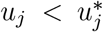. Thus, the two plateaus are separated by a low-probability well. By inspection, and from Figure S11c, we can see also that the Hessian *W*^*′′*^(*u*_*j*_) does not have definite sign, meaning that the prior is not log-concave. In addition, the necessity to truncate the original distribution becomes obvious as we compare between Figure S11c&d. The log prior distribution as a function of *u*_*j*_ is unbounded on ℝ, causing numerical instability in implementation. **Re-parametrization of the original KS distribution**. To select the desired parameters *a, b*, we introduce a reparameterization of the original KS distribution PDF. Specifically, we reparameterize the PDF w.r.t *a, r* instead of the original *a, b*. We start from the CDF of the original KS distribution (E.2) and set *F*_*X*_(0.5) = *P* (*X* ≤ 0.5) = *r*:

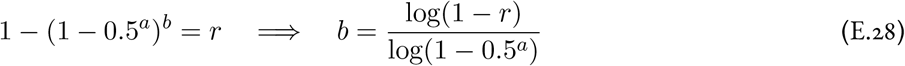

Hence we have 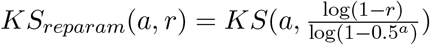. Under this parameterization, we interpret *a* as a shape parameter and *r* as a bias parameter that balances the density near the two end points. This parameterization is much easier to interpret and useful when finding a desired distribution. Although we lose the exact relation that *P* (*X*≤ 0.5) = *r* after the truncation process, the qualitative properties of the distribution is mostly unchanged. Hence, this reparameterization is still helpful for selecting parameter for truncated KS distribution. Particularly, in the simulations, we let *a* = 0.055 and *r* = 0.75, which yields (*a, b*) = (0.055, 0.422) in the original expression.

### F Presence-dependent concentration priors

For most of this paper, we have imposed the simplifying assumption that the conditional prior on concentration is the same irrespective of whether or not an odorant is present. However, such a formalism yields an artifact that our latent concentration estimates *c*_*i*_ does not goes to 0 when *p*_*i*_ = 0. To address this, we introduce a presence-dependent concentration prior that favors having *c*_*i*_ = 0 when *p*_*i*_ = 0. We design a differentiable log-prior that interpolates between a sparsity-encouraging exponential distribution (that is, an *L*_1_ penalty) with rate *λ* at *p*_*i*_ = 0, and a Gamma distribution with parameters *α* and *β* at *p*_*i*_ = 1. Writing the two component log-densities as log Exp(*c*_*i*_ | *λ*) = log *λ* −*λc*_*i*_ and log Gamma(*c*_*i*_ | *α, β*) = log(*β*^*α*^*/*Γ(*α*)) + (*α* −1) log *c*_*i*_ − *βc*_*i*_, we define the log-prior as their presence-weighted combination, so that *p*_*i*_ = 0 recovers the exponential and *p*_*i*_ = 1 the Gamma:

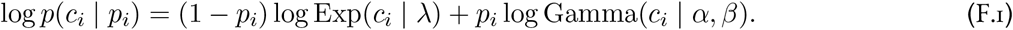

Equivalently *p*(*c*_*i*_ |*p*_*i*_) ∝ Exp(*c*_*i*_| *λ*)^1*−pi*^ Gamma(*c*_*i*_ | *α, β*)^*pi*^, a geometric (rather than arithmetic) interpolation, which keeps the log-prior — and hence the gradients below — linear in *p*_*i*_. In vector form,

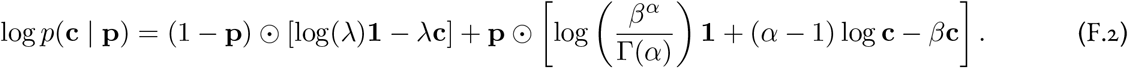

Its derivatives with respect to **p** and **c** are

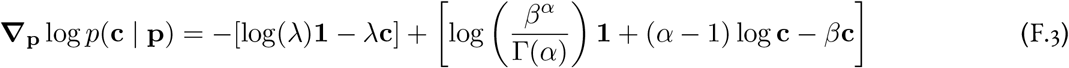

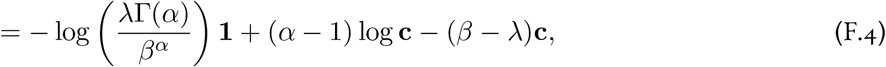

and

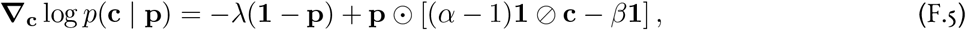

respectively. With the same definition of the energy function as before, we have

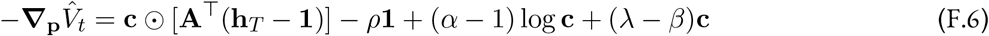

and

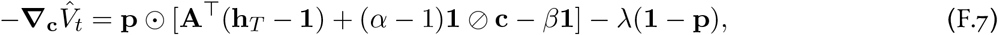

where we now let

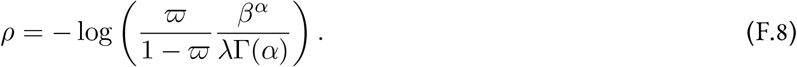

Thus, the full SDEO model dynamics with this prior are, in evidence-wrapped form,

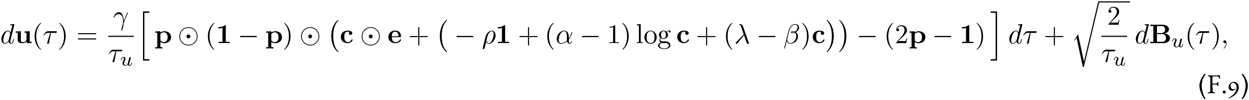

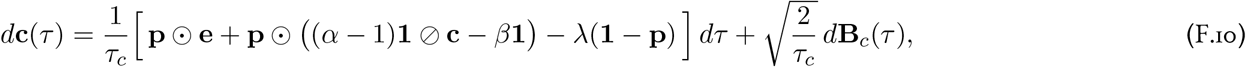

with soft (tufted) evidence **e** = **A**^*T*^(**h**_*T*_ − **1**). Expanding the evidence down to the cell activity **h**_*T*_ gives the detailed form

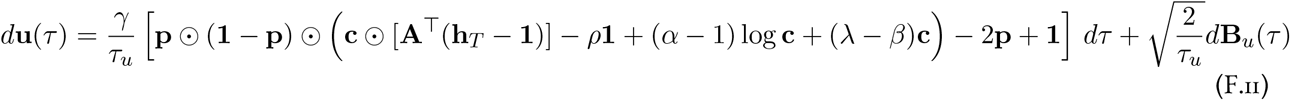

and

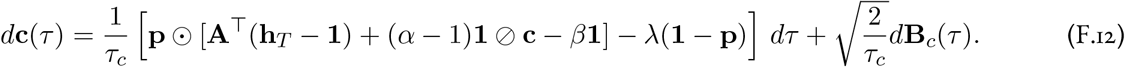

This model is soft-gated by design — the presence-dependent prior couples **c** to the continuous **p** — so both updates use the tufted error **h**_*T*_ = **s**⊘ (**r**_0_ + **A**(**c ⊙ p**)), and no hard mitral channel enters here. We show an example simulation of these dynamics in Figure S3. In contrast to Figures 3, S1, and S2, we see that with this choice of prior the latent concentration estimates for non-present odors decay to zero once the presence estimate indicates that the odor is absent. Moreover, in the dynamics above, we see that for *α >* 1 the update to the presence estimate coming from the conditional prior on **c** | **p** has an interesting effect: it will tend to push low-**c** odors towards being recognized as absent, as (*α* − 1) log **c** will become strongly negative.

### G Adaptive prior on presence

In the previous section, we introduced the Bernoulli and Kumaraswamy distributions for modeling odor presence during the inference. These priors encode sparsity in odor scene but they both assume a static distribution. The same prior distribution is used at every time point during inference, independent of previous sensory evidence. This assumption is suitable for snapshot inference, but it neglects an important temporal structure of natural olfactory scenes. In real-world scenarios, odor concentration can fluctuate quickly, but odor presence usually has temporal persistence. So the model should not treat each time point as an independent static inference problem. Instead, it should carry forward a belief about which odors are likely to remain present over time.

This extension also has a more natural circuit interpretation. In our framework, the prior over odor presence can be viewed as a form of cortical feedback to the olfactory bulb. Real corticobulbar feedback is unlikely to provide a fixed, time-invariant prior over external stimuli. Instead, it may convey adaptive and context-dependent expectations shaped by recent sensory history, behavioral state, or environmental context. Motivated by this interpretation, we extend the static presence prior to a time-dependent prior whose parameters are updated according to the model’s evolving belief about odor presence. We refer to this belief-driven prior as a dynamic prior on odor presence.

Note that the prior over concentration could also be time-dependent by imposing a dynamical rule on concentration. However, because odor concentration is naturally highly fluctuating, we keep the Gamma prior over concentration time-independent and introduce temporal structure only through the prior over odor presence. Importantly, this enables the timescales of concentration inference and presence inference to be separated. Since concentration is not constrained by a temporal transition rule, it can vary rapidly from one inference window to the next. In contrast, odor presence is governed by a dynamical transition rule. Therefore, when the switching rates are small, the inferred presence state evolves slowly over time.

We keep the assumptions of the static SLAM model except that odor presence evolves in time according to a two-state Markov process.

Before receiving the next sensory input, the system should already have a prediction about which odors are likely to remain present. Then, when new receptor activity arrives, the system corrects this prediction based on the current evidence. This can be achieved using recursive Bayesian filtering, as commonly used in SLAM. The inference is decomposed into a time update, which carry forward prior belief over presence of odors, and a measurement update, which corrects the belief using new sensory observations.

The time update is based on a “prior” of transition rule over how the state evolves from one time point to the next.

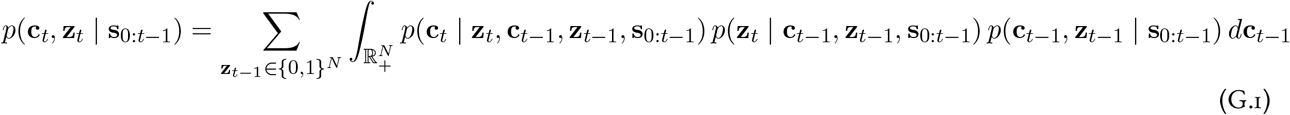

We assume the system satisfies the Markov property. The current state (**c**_*T*_, **z**_*T*_) is conditionally independent of the observation history given the previous state, so the transition factors no longer depend on **s**_0:*t−*1_:

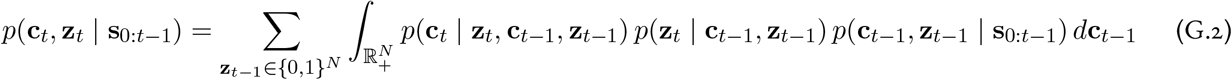

We also assume **c**_*T*_ is conditionally independent of last state (**c**_*t−*1_, **z**_*t−*1_) given **z**_*T*_, and **z**_*T*_ is independent of **c**_*t−*1_. We consider each *z*_*T,i*_ ∈ {0, 1} is its own 2-state Markov chain

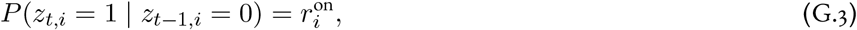

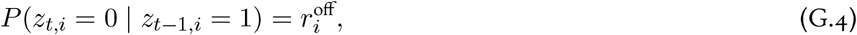

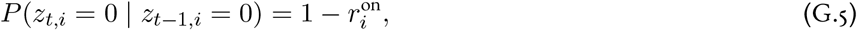

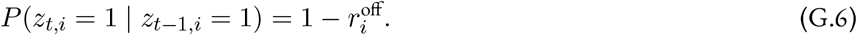

We further assume *z*_*T,i*_ are independent across *i* given **z**_*t−*1_. Then we have

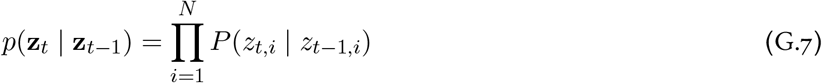

Then the time update can be simplified to

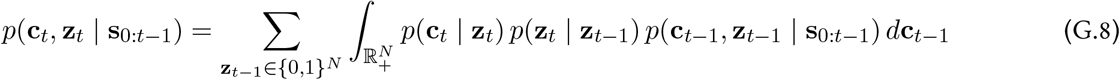

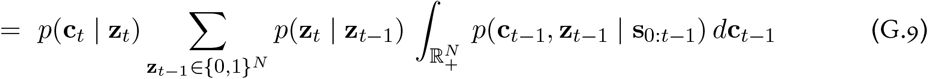

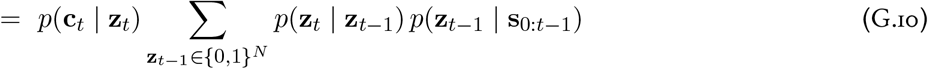

The measurement update follows the same logic as the original static inference framework, but the “prior” is predictive instead of a static distribution.

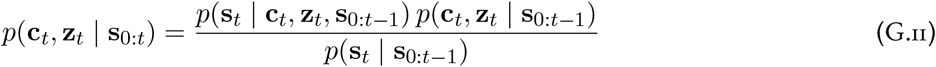

Under the conditional independence assumption for the observation

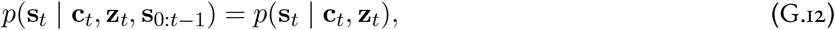

the measurement update can be simplified to

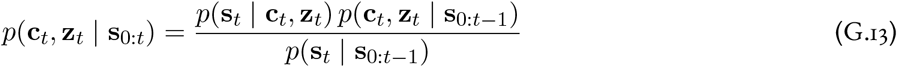

Because our model estimates states through sampling, maintaining the full predictive distribution during the time-update step is not straightforward. Instead, we let the time update track the expected value of the prediction, and then use this predicted expectation as the parameter of a Bernoulli distribution. This Bernoulli distribution serves as the predictive prior in the subsequent measurement-update step. We define

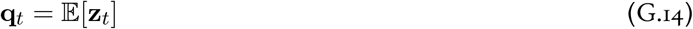

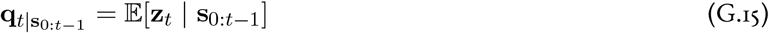

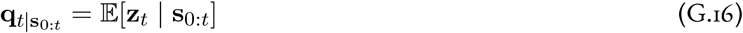

From the time update we have

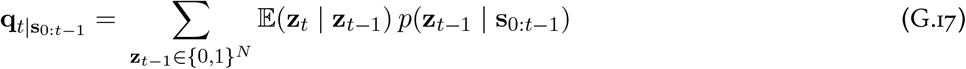

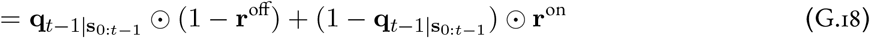

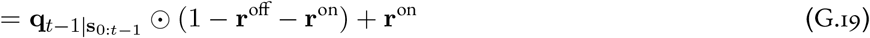

The dynamics can be interpreted as a form of cortical feedback. Rather than being fixed, this feedback can have its own dynamics and evolve according to the recent inferred state of the odor scene.

To perform sampling from the measurement update, we use the same MLD framework from the previous derivation by defining a relaxed variable **p**. In practice, we use the relaxed **p** from the measurement update to approximate 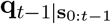. We also introduced a constant hyperparameter *w* on the prior term to control the extent to which the prior influences the inference. Thus the dynamics are

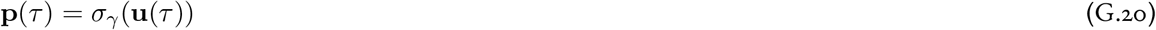

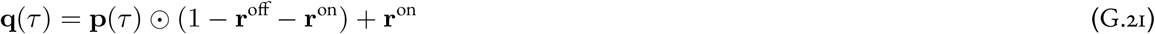

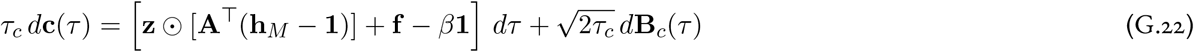

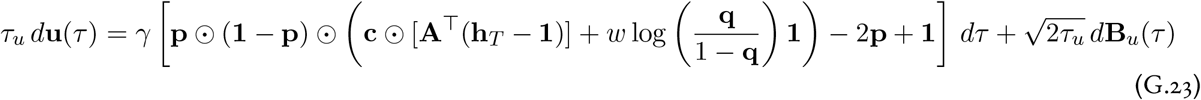

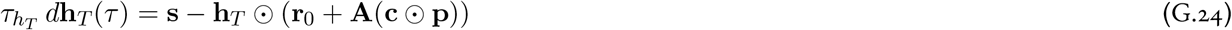

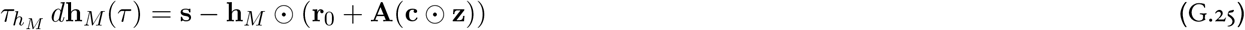

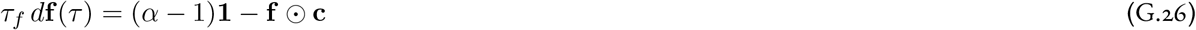

This detailed (cell-activity) system differs from the basic circuit model of Section D.9 only in the presence prior: the static log-odds 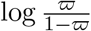 is replaced by the belief-driven 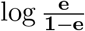, with the predictive presence **e** carried by the time-update equation. In evidence-wrapped form — writing the hard (mitral) evidence 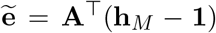 for the concentration channel and keeping the soft (tufted) evidence **A**^*T*^(**h**_*T*_ **1**) explicit (here the bare symbol **e** is reserved for the predictive presence) —the two estimation channels read

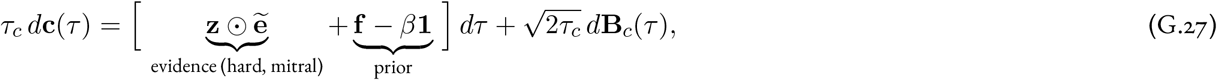

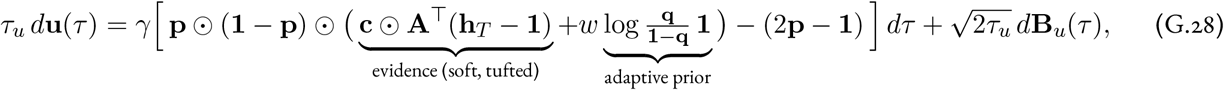

with cell activities **h**_*T*_ = **s**⊘ (**r**_0_ +**A**(**c ⊙ p**)) and **h**_*M*_ = **s**⊘ (**r**_0_ +**A**(**c ⊙ z**)), the feedback population **f** → (*α* − 1)**1 ⊘ c** of Section D.9, and **e** evolving by *τ*_*e*_ *d***e** = **p**⊙ (1 −**b** −**a**) + **a**.

The dynamics using adaptive presence prior is shown in Figure S8. Compared with static presence priors, such as Bernoulli distribution and Kumaraswamy distribution, the adaptive presence prior yields a more stable estimate of odor presence and allows the concentration estimate to more closely track the oscillations. This is particularly important in natural odor sensing, where odor concentration is highly fluctuating.

### H Mutual coherence of affinity matrix and scaling capacity

In this section we will discuss our model from a pure compressive sensing perspective, focusing on how the properties of the random sensing matrix—in our case, the affinity matrix—affect the scaling capacity. The key property of interest is the *mutual coherence* of the sensing matrix, which is one measure of how closely the measurement approximates an isometry. It is important to note that compressed sensing with Poisson noise is not as well-understood as the standard case of additive Gaussian noise [27, 104]. However, many of the same desiderata for the sensing matrix carry over. Namely, the mutual coherence should ideally be small.

The mutual coherence is a measure of the worst-case similarity between the columns of a projection matrix ***A***. It is defined as:

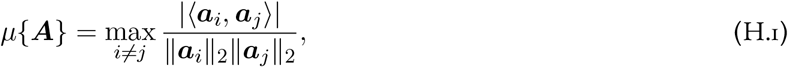

where ***a***_*i*_ is the *i*-th column of **A**. Equivalently, the mutual coherence is the maximum absolute value of the off-diagonal elements of the normalized Gram matrix

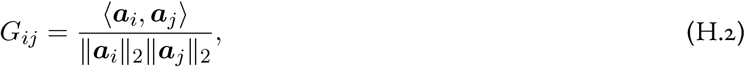

*i.e., µ*{*A*} = max_*i*≠*j*_ |*G*_*ij*_| To achieve good compressed sensing performance, the mutual coherence should be small.

However, the mutual coherence is a worst-case measure, and thus can give extremely pessimistic predictions relative to the performance of a particular sensing matrix in practice. On these grounds, Elad [67] argued that the *t-averaged mutual coherence*

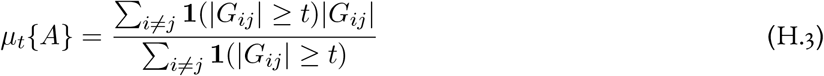

can provide a more informative measure. Here, **1**(·) is an indicator function which is equal to one when the predicate in its argument is true, and zero otherwise.

Using these two measures, we can evaluate the quality of an affinity matrix ***A***. Particularly, we will use the *t-averaged mutual coherence*, where the *T* value is set to be the 80% quantile of the Gram matrix, which gives us the average of top 20% worst cases.

We are interested in three ensembles of random affinity matrices:

1. **Dense gamma affinity matrix**: each element of the affinity matrix follows a Gamma distribution, and the matrix is normalized by its largest element;
2. **Sparse binary affinity matrix**: each element of the affinity matrix follows a Bernoulli distribution, being 1 with a probability *p* and 0 otherwise;
3. **Sparse gamma affinity matrix**: each element of the affinity matrix follows Gamma distribution with probability *p* and 0 otherwise.

We plot examples of these three types of sensing matrices in Figure S12a.

In Figures 8 and S10, we saw that sparse binary affinity matrices seem in general to achieve the highest capacity, followed by sparse gamma and then dense gamma. Examining the mutual coherence distributions for each of these ensembles, we see that a moderate system size (dictionary of 5000 odorants, 600 sensors), we notice that the sparse Gamma sensing matrix has the lowest average mutual coherence *µ*_*t*_ but the sparse binary sensing matrix has the lowest mutual coherence *µ* (Figure S12b). The average mutual coherence varies with the dictionary size and with the number of sensors, but this ordering remains roughly consistent across scales (Figure S12c).

In the sparse sensing ensembles, a crucial parameter governing the mutual coherence is the mean sparsity (average fraction of non-zero elements) *p* [27]; sparser matrices have lower average mutual coherence (Figure S12d). As we would expect, this results in a scaling capacity that decreases with increasing *p* across a variety of dictionary and sensor repertoire sizes (Figure S12e-f). Therefore, our empirical results are consistent with the broad conclusion that lower average mutual coherence enables higher compressive sensing capacity.

### I Numerical methods and additional scaling results

#### I.1 General numerical method pipeline

This section illustrates the general numerical analysis pipeline used in this paper. As an example, we showed how we numerically simulate the SDEs (D.49), (D.56) and (E.16). These equations describe the full model dynamics under the assumption of 1-on-1 coding in granule cells. They use either the Bernoulli prior (D.49) or the Kumaraswamy prior (E.16) on the presence. Because the single-sample online regime advances the sampling time *τ* by one step per physical step (Section B.3), here the sampling time *τ* and the physical time *t* coincide; there is no timescale separation, so no separate Δ*τ* is needed and we discretize directly in physical time *t*. We use a step size Δ*t* = 10^*−*5^, resulting in a total of 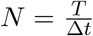 timesteps over an infer-ence window of physical duration *T*. Afterward, applying the Euler-Maruyama method, along with additional treatments to stabilize the numerical computation, we obtained a discrete-time Markov chain (*X*_*n*_ : 0≤ *n* ≤*N*) representing the evolution of the estimated **c** and **u** over time. Since the subscript *n* is already used to denote the time step, we use **p**_*n*_[*i*] to denote the *i*-th element of the vector **p**_*n*_ in this section. In particular, we have 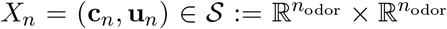 with its transition rules given by:

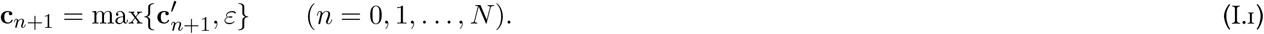

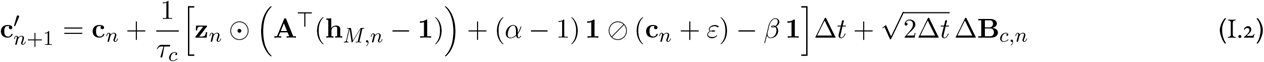

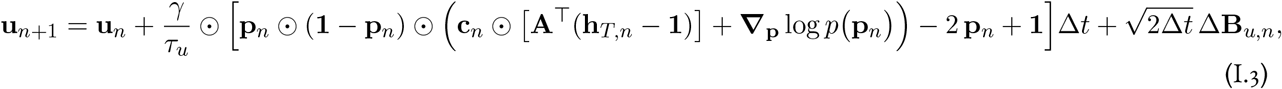

where

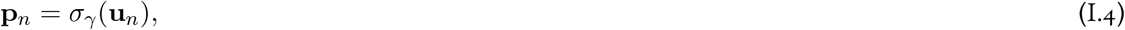

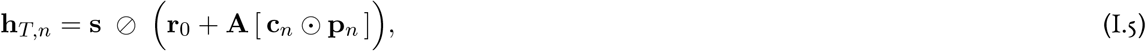

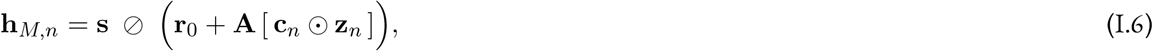

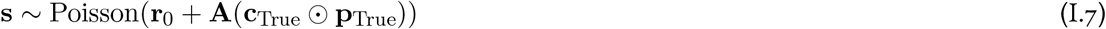

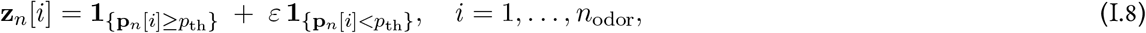

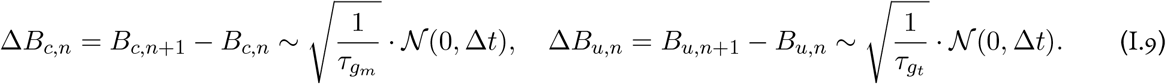

and

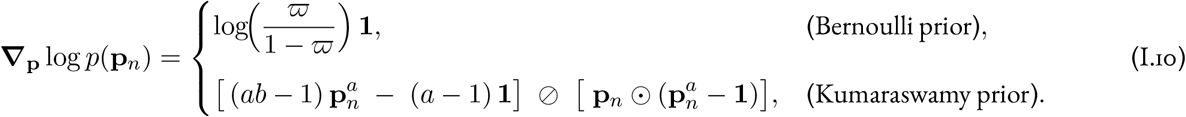

Here **h**_*T,n*_ (soft, tufted) and **h**_*M,n*_ (hard, mitral) are the prediction-error ratios introduced with the gradient derivation: **h**_*T,n*_ − **1** is the relative prediction error under the soft presence **p**_*n*_ and **h**_*M,n*_ − **1** its hard-gated counterpart, so the back-projected quantities **A**^*T*^(**h**_*T,n*_ − **1**) and **A**^*T*^(**h**_*M,n*_ − **1**) are the bottom-up evidence **e**_*n*_ and 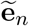 that drive the presence and concentration updates, respectively. The vector is **z**_*n*_ the hard-gated binary presence **z** = *f*(**p**) of (D.56): an odorant whose inferred presence reaches the threshold *p*_th_ is set to 1, while one below threshold is floored to *ε* (rather than exactly zero) for numerical stability, so odorants inferred absent are decoupled from the concentration update. In particular, we used the following hyperparameter settings: 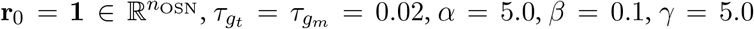 and *ε* = 10^*−*5^. For the Bernoulli prior, we set *ϖ* = 0.1 and *p*_*th*_ = 0.2; for the Kumaraswamy prior, we set *a* = 0.1, *b* = 0.5 and *p*_*th*_ = 0.5. The affinity matrix **A** is one of the three types of random matrix discussed in Appendix H. Since the Markov chain produced by the Euler-Maruyama method approximates both the trajectory and stationary distribution of the SDEs, its behavior—particularly empirical convergence and convergence rate—reflects the model’s capacity to infer the presence and concentration of odorants. In general, analyzing the behavior of this Markov chain suffices as a numerical approach for evaluating the model’s system of SDEs. Unless otherwise specified, the same numerical procedure was applied to variants of the basic model. Modifications specific to each variant are discussed in the corresponding sections. We summarize all parameter used in the scaling simulation in Table 2.

**Table 1:** Variable and parameter names.

| Variable name | Description | Type / Domain |
| --- | --- | --- |
| **Dimensions** |  |  |
| $n_{\text{odor}}$ | Number of potential odorants (Dictionary size) | $\mathbb{N}$ |
| $n_{\text{OSN}}$ | Number of sensors (Sensor repertoire size) | $\mathbb{N}$ |
| $n_P$ | Number of odorants present | $\mathbb{N}$ |
| **Variables** |  |  |
| $\mathbf{s}$ | OSN responses | $\mathbb{R}_+^{n_{\text{OSN}}}$ |
| $\mathbf{h}_T$ | Responses of soft-gated projection neurons | $\mathbb{R}_+^{n_{\text{OSN}}}$ |
| $\mathbf{h}_M$ | Responses of hard-gated projection neurons | $\mathbb{R}_+^{n_{\text{OSN}}}$ |
| $\mathbf{u}$ | Estimated presence in the dual space for all potential odorants | $\mathbb{R}^{n_{\text{odor}}}$ |
| $\mathbf{p}$ | Estimated presence in the primal space for all potential odorants | $[0, 1]^{n_{\text{odor}}}$ |
| $\mathbf{z}$ | Gated binary estimated presence for all potential odorants | $[0, 1]^{n_{\text{odor}}}$ |
| $\mathbf{c}$ | Estimated concentration for all potential odorants | $\mathbb{R}_+^{n_{\text{odor}}}$ |
| **Single-neuron parameters** |  |  |
| $r_0$ | Baseline firing rate of OSN | $\mathbb{R}_+^{n_{\text{OSN}}}$ |
| **Population parameters** |  |  |
| $\mathbf{A}$ | Affinity matrix | $\mathbb{R}_+^{n_{\text{OSN}} \times n_{\text{odor}}}$ |
| $\alpha$ | Parameter for the Gamma prior distribution $\text{Gamma}(\alpha, \beta)$ | $\mathbb{R}_+$ |
| $\beta$ | Parameter for the Gamma prior distribution $\text{Gamma}(\alpha, \beta)$ | $\mathbb{R}_+$ |
| $\gamma$ | Gain of the sigmoid mirror map $\text{Gamma}(\alpha, \beta)$ | $\mathbb{R}_+$ |
| **Notation** |  |  |
| $\odot$ | Element-wise (Hadamard) vector product | Notation |
| $\oslash$ | Element-wise (Hadamard) vector division | Notation |
| $\mathbf{B}$ | Brownian noise (Langevin) | Notation |
| $T$ | Time available for inference | Notation |
| $\mathbf{1}_{\{\cdot\}}$ | Indicator function | Notation |

**Table 2:** Hyperparameters used in simulations.

| Variable name | Description | Value | Unit |
| --- | --- | --- | --- |
| **Single Neuron parameters** |  |  |  |
| $r_0$ | Baseline firing rate of OSN | 1.0 | |
| $\tau_c$ | Time constant for concentration estimation | 0.03 for circuit impl.; 0.02 otherwise | second |
| $\tau_p$ | Time constant for presence estimation | 0.03 for circuit impl.; 0.02 otherwise | second |
| $\tau_f$ | Time constant for feedback dynamics | 0.02 for circuit impl. | |
| $\tau_h$ | Time constant for predictive error signal | 0.02 | second |
| **Prior distribution for concentration** |  |  |  |
| $\alpha$ | Parameter for Gamma($\alpha, \beta$) | 5.0 | |
| $\beta$ | Parameter for Gamma($\alpha, \beta$) | 0.1 | |
| **Prior distribution for presence** |  |  |  |
| $\varpi$ | Parameter for the continuous Bernoulli prior distribution | 0.01 | |
| $a$ | Parameter for truncated Kumaraswamy distribution | 0.055 | |
| $b$ | Parameter for truncated Kumaraswamy distribution | 0.422 | |
| $\epsilon$ | Truncation cutoff for truncated Kumaraswamy distribution | $5 \times 10^{-5}$ | |
| **Presence and $\mathbf{u}$ in dual space** |  |  |  |
| $p_{th}$ | Threshold used for binarizing the continuously relaxed presence | 0.2 | |
| $\gamma$ | Parameter in the sigmoid function that transform $\mathbf{p}$ into $\mathbf{u}$ | 5.0 | |
| **Non-separated model** |  |  |  |
| $\tau_E$ | Time constant for excitatory neurons | 0.02 | second |
| $\tau_I$ | Time constant for inhibitory neurons | 0.03 | second |
| $\lambda$ | Parameter for the $L_1$ exponential prior distribution | 1.0 | |
| **Simulation parameter** |  |  |  |
| $\Delta t$ | | $1 \times 10^{-5}$ | second |

#### I.2 Further computational implementation and computing resources

All scaling simulations were implemented in Python 3.11 using the PyTorch framework (version 2.6.0+computecanada) and executed on AMD EPYC 9655 CPU nodes of the Digital Research Alliance of Canada Fir cluster. Several runtime optimizations were applied to improve performance and ensure stability across different CPU types and under varying levels of cluster load. Because iterated Euler integration involves repeated multiplication of large matrices—an increasingly expensive operation as dimensionality grows—single-simulation runtime optimization focused on efficient matrix computation. We used PyTorch tensors instead of NumPy arrays for most matrix operations to leverage the multi-processing in PyTorch. Further speed-up was obtained by using PyTorch’s Just-In-Time (JIT) compilation for the core Euler-forward SDE integration kernels. In addition, we parallelize matrix computations across 4 CPU cores for the baseline model with OpenMP. All of these optimizations yield a 2-3x speedup over vanilla NumPy implementations in high-dimensional settings. At the experiment level, each sweep experiment was partitioned into multiple jobs and dispatched as independent tasks using SLURM’s array feature. File I/O was handled via the h5py library for efficient data storage and convenient retrieval. Additional algorithmic optimizations for scaling capacity experiments are discussed in Appendix I.4.4. Specific runtime varies across platform. In our case, one simulation with 5 ×10^4^ iteration steps at the highest dimensionality (16K possible odorants) took at most 1200s for SDEO model, and up to ~7000s for the baseline model. Thus, a significant proportion of the CPU hours was spent running the baseline comparison in Figures 8 and S10. Total compute time required to generate all the figures is ~12000 CPU hours.

#### I.3 Baseline model and metrics for evaluation

##### I.3.1 Baseline model

To demonstrate the improved capacity of our model through comparison, we use a model we proposed in [39] as a baseline. Specifically, we use the proposed model with one-to-one coding between neurons and odorants. We confirm that the performance of the baseline model we obtain here is consistent with the results in previous work. For validation of consistency, Figures S9 and S10 in this work can be indirectly compared with Figures 3 in [39]. We note that the displayed performance appear worse because we adopt stricter evaluation criteria and shorten available time for inference to ensure comparability with the high performance of the SDEO model. For example, we define correct estimate as one in which the estimated concentration falls within ±25% × True Concentration, while the previous work use ±50%.

##### I.3.2 Metrics for concentration estimation

We introduce the two metrics for evaluating the accuracy of the concentration inference. Recall that we denote the number of presented odors as *n*_present_, for any concentration estimates **ĉ** given the model, we have:

1. **Mean absolute error:** We compute the mean absolute error (MAE) between inferred and true concentrations for the presented odors: 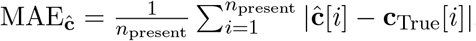. In Figures 7 and S9, MAE is plotted as curves in panel **a** and as heatmaps in panel **b**.
2. **Correct proportion:** We calculate the proportion of presented odors whose predicted concentrations fall within a ±*δ* neighborhood of the true value: 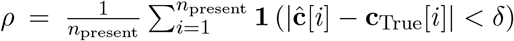. The correct proportion computed using *δ* = 10 is shown as contours in panel **b** of Figures 7 and S9. We note that since we set the true concentration value to be 40, the correct proportion with *δ* = 10 corresponds of the proportion of correct concentration estimates within a tolerance of ±25%×True concentration.

##### I.3.3 Metrics for presence estimation

To introduce metrics for the quality of the presence inference, we begin by noting that the detection of odorants is inherently a binary classification problem. We therefore adopt a metric, AUROC score, from receiver operating characteristic (ROC) analysis to quantify the probability that the presence of odorants can be decoded from based on the output of the model. In ROC analysis, the ROC curve is the function of the true positive rate (TPR) over the false positive rate (FPR), which can be plotting by interpolating the TPR versus FPR at all possible thresholds. The AUROC score (*AUC* ∈ [0, 1]) is defined as the area under the ROC curve. It can be computed given estimated value vector 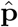 and a ground truth label vector **p**_True_. Importantly, its value is exactly the probability of responding correctly in the two-alternative forced-choice test based on a given data [105], and it reflects the discrimination between the representation of present and absent odorants in 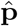. AUROC score can also be interpreted as the integration of the accuracy over all possible classifying thresholds. We briefly showed that this interpretation aligns with the standard definition of AUROC score. Let *θ* denote the threshold that downstream neurons use to classify **p**. Define:

- the true positive rate as *β*(*θ*) = *P* (*p* ≥ *θ*|present)
- the probability density function of presence *p* given the odorant is absent as 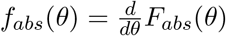 where *F*_*abs*_(*θ*) = *P* (*p* ≤ *θ*|absent).

We then have the probability of decoding correctly as:

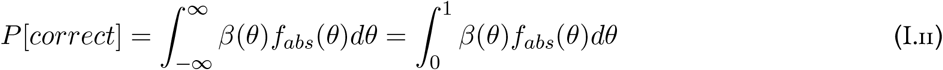

Now consider the false positive rate *α*(*θ*) = *P* (*p* ≥ *θ*|absent), notice that:

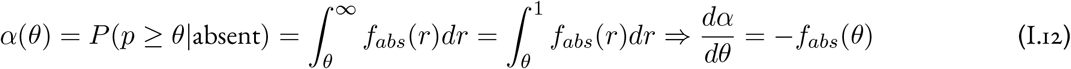

Substitute (I.12) into (I.11) we have:

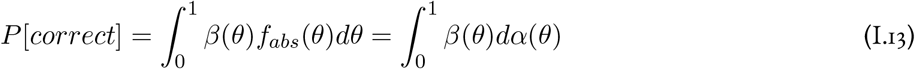

Given that the ROC curve is *β* = *ψ*(*α*), where *ψ* : [0, 1] ↦ [0, 1] maps the FPR *α* to the TPR *β*, the right hand side of (I.13) corresponds to the area under the ROC curve. Therefore, the AUROC score is indeed the probability of correct classification in a 2AFC task. As a direct consequence of its definition, AUROC score is threshold-independent. It also inherently incorporates both TPR and FPR in a single scalar value and reflects the discrimination between the two classes in the data we are performing the classification on. In comparison, most other metrics cannot account for TPR and FPR simultaneously, requiring additional metric as complementary indicator. Therefore, AUROC score provides a more comprehensive assessment of a classifier’s performance by considering the probability of correct classification than traditional threshold-based binary classification metrics. An AUROC score of 0.85 indicates that, for any randomly chosen pair consisting of a present odor and an absent odor, the model will correctly classify the present odor with a probability of 85%. Effectively, this means that, on average, 85% of all odors can be correctly classified, and in particular, 85% of the presented odors can be correctly classified as present. We can see that although the AUROC score captures more information about the model’s presence estimation performance, the two metrics have some extent of equivalency in their interpretation, and this allows a clear and direct comparison between the two models. Hence we use the AUROC score as the primary metric for evaluating the presence estimation performance of our model. In Figures 7 and S9, we plotted the AUROC score as curves in panel **c** and as heatmaps with contours overlaid in panel **d**.

##### I.3.4 Maximum detection capacity

Lastly, to investigate the scaling properties of the SDEO model, we introduce the *maximum detection capacity κ* to quantify the largest number of simultaneously presented odorants that the model can reliably detect given a certain number of sensors and dictionary size. For a fixed *n*_OSN_ and *n*_odor_, we first define *maximum detection capacity* assessed by presence estimates: *κ*_AUROC_. Specifically, we let it to be the largest *n*_present_ such that the AUROC score on model’s presence estimates 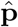 remains above a threshold *ϵ*_*T*_:

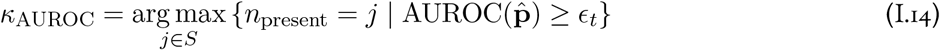

where *S* ⊂ N denotes the search space. In practice, the search space is chosen to balance computational cost and the resolution. Similarly, we define its counterpart that evaluates the concentration estimates as *κ*_MAE_. Notably, since larger MAE implies worse performance, *κ*_MAE_ is defined by MAE remaining less than some threshold *ϵ*_*T*_:

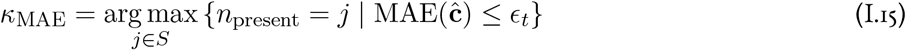

In Figures 8 and S12 (panel **f**), we plot the maximum detection capacity evaluated by the AUROC score as heatmaps with contours overlaid. The threshold used is the AUROC score = 0.85. In Figure S10, we plot the maximum detection capacity evaluated by the MAE as heatmaps with contours overlaid. The threshold used is 10 concentration unit. This represents a 25% relative error tolerance as the true concentration is 40.

#### I.4 Experimental setting and supplemental figures

##### I.4.1 Single simulation

Here we explain how a single simulation—the building block of all scaling experiment—is executed.

The first step of each simulation is to generate sensory scene, which includes generating ground truth time series, affinity matrix for OSNs, and multiplying it with ground truth time series to get OSNs responses. We considered the OSN responses **s** to be static throughout the inference, hence the values of **s** are drawn once from the Poisson distribution in (I.7). Since the affinity matrix is randomly generated in each trial, without loss of generality, we always pick the first *n*_present_ odorants in the dictionary to be the presented odors in the implementation level.

This is followed by the initialization of network. Each components of the concentration vector **c**_0_ is independently drawn from a Gamma distribution: **c**_0_[*i*] ~ Gamma(6, 4) for *i* = 1, …, *n*_odor_, while each components of the presence vector **p**_0_ was set such that 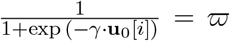. Consequently, the initial concentration states are randomly assigned, while the initial presence values correspond to their prior probability *ϖ*. In our simulations, the prior probability *ϖ* was set to 0.01 for all odorants. However, these prior need not be identical—they can be assigned heterogeneously or updated adaptively based through a learning process. In fact, non-uniform and adaptive prior may better reflect the implementation in biological system. Further exploration on the dynamical prior and OSN responses were left for the future work.

After initialization, we run the RNN through the forward integration method discussed in Appendix I.1, and record data at some chosen sample rate into a tensor, where the further analysis and metrics are computed.

##### I.4.2 Dynamics demonstration

In Figures S1 and S2, we show the dynamics of the model during the process of estimating a single group of odorants, under sparse binary and dense gamma sensing matrices respectively.

We set the the number of presented odors *n*_present_ = 5, the dictionary size *n*_odor_ = 500, and the number of sensor *n*_OSN_ = 300. Other parameters remain the same as in Table 2 except we use *γ* = 1 in dynamics simulation.

We set the appearance of odorants to be 0.25 sec after the network starts running. Then the odorants are appears with concentration 40 for a duration of 0.5 seconds, while the network runs for another 0.25 sec after odorants disappear. This design allows us to illustrate the baseline steady state of the network right after initialization and after odorants disappearing. We record the activities of neurons in the network with a sample rate of 1000 Hz.

In Figure 3, we show the dynamics of the model during the process of estimation two groups of odorants, under sparse binary sensing matrices. The simulation procedure remains the same, except that we generate a sensory scene that involves independent groups of odorants with slow-changing concentration. We plot the output concentration estimate **c** *T***p** to illustrate the final output of our model.

We set the number of presented odors *n*_present_ = 6, arranged in two groups of three odorants. We maintain all other setting the same as in the single group dynamics simulation above.

##### I.4.3 Simultaneous and rapid detection capacity

In Figures 7 and S9, we assess the model’s ability to infer the presence and concentration of multiple odorants simultaneously presented.

We vary the number of presented odors *n*_present_ from 1 to 100, while fixing the total number of sensors *n*_OSN_ = 300 and the dictionary size. For simulation using the sparse binary sensing matrix, we let *n*_odor_ = 1000 (Figures 7 and S1). For simulation using the dense Gamma sensing matrix, we set *n*_odor_ = 500 (Figure S9). This is because using a sparse binary matrix improves the capacity of the model.

For each 1 ≤*n*_present_ ≤100, we run 40 independent simulations. In each simulation, we present *n*_present_ odorants to the model for an interval of length *T* = 0.75*s* and record the models’ estimates with a sample rate of 100 Hz. Under this setting, a slice of the recorded tensor at some timepoint *T* represent the estimates produced by the model when given an available time window of *T*s. To assess the quality of the estimates, we compute three metrics score from these slices: mean absolute error (MAE), correct proportion with tolerance *δ*=10, and AUROC score (defined in Appendix I.3.2). We then take the averaged scores throughout the 40 trials as the final results.

Results of this simulation are shown in Figures 7 and S9. We first focus on evaluating the concentration estimates. The averaged MAE for each 1≤ *n*_present_≤ 100 over the time course of *T* = 0.75*s* is shown as heatmaps in panel **b**. As a complement, smoothed contours of equal correct proportion are overlaid on top of the heatmaps. While MAE shows the quantitative measure of the accuracy, the correct proportion scores give more qualitative indication of the model performance. For visualization purposes, we smoothed the contours with a Gaussian filter of standard deviation *σ* = 1. Across heatmaps, we ensured the color axis are the same so the fair visual comparison across different conditions can be made.

Furthermore, we plot vertical slices of the heatmaps at *T*_1_ = 100ms and *T*_2_ = 600ms separately in panel **a**. They shows as MAE as a function of *n*_present_ at that two timepoint. The shaded area around the curves represents ±1.96 ×*SEM*, indicating 95% confidence interval of the mean.

When evaluating the presence estimates, similar procedure is repeated. The averaged AUROC score for each 1≤ *n*_present_ ≤ 100 over the time course of *T* = 0.75*s* is shown as heatmaps in panel **d**. Contours of equal AUROC score are overlaid. Vertical slices of the AUROC score heatmaps at *T*_1_ = 100ms and *T*_2_ = 600 are also plotted as curves with shaded region indicating 95%C.I. in panel **c**.

##### I.4.4 Scaling capacity

In Figures 8 and S10, we aim to investigate how the number of receptor types *n*_OSN_ required to effectively detect a fix number of presented odors *n*_present_ scales with the dictionary size *n*_odor_. Thus, we vary *n*_OSN_ linearly from 100 to 800 with an increment of 50 and sampled *n*_odor_ over 16 evenly spaced points in log scale ranging from 1000 to 16000. This created a 2D grid of (*ñ*_OSN_, *ñ*_odor_) pairs.

For each pair of (*ñ*_OSN_, *ñ*_odor_), we perform three independent runs of binary search to find the *maximumsimultaneous detection capacity κ* defined in Appendix I.3.4, where we set the search space *S* = {*n* |*n* = 5*i, i* = 1, …, 20}. Particularly, in Figure 8, we use maximum detection capacity assessed by presence estimates *κ*_AUROC_ with a threshold *ϵ*_*T*_ = 0.85; in Figure S10, we use its concentration counterpart *κ*_MAE_ with a threshold of 10, while the true concentration is 40. In each binary search, we ran simulations under the condition *n*_OSN_ = *ñ*_OSN_, *n*_odor_ = *ñ*_odor_, and *n*_present_ = *j*, where *j* ∈ *S* is taken from the sequence (search path) 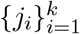 produced by the binary search algorithm. The available time for estimation *T* is 0.2 sec in Figure 8 and 0.5 sec in Figure S10. The concentration is harder to estimate so we increase the time available.

We then used the averaged results of the three independent run, 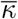, as the final estimate of the maximum simultaneous detection capacity. We repeated this process across the 15 × 16 pairs of (*ñ*_OSN_, *ñ*_odor_), and visualized the resulting grid of 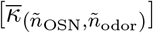 as a heatmap. Contours of equal AUROC score or MAE is further overlaid onto the heatmaps. All heatmaps within the same figures share the same color axis.

The use of binary search design significantly reduces the computational cost, without whom the experiment would be computationally infeasible, as binary search guarantees we can find the *κ* using at most ⌈log_2_(|*S*|) ⌉ = 5 simulations, whereas a naive linear search would easily require more than 10 simulations.

Using this efficient pipeline, we are not only able to directly compare the scaling capacity of our SDEO model with the non-separated baseline model, but also investigate the effect of the affinity matrix and presence prior on the scaling capacity. We tested 9 combinations between three types of affinity matrix (Appendix H) and three types of model variants (non-separated, SDEO with Bernoulli prior and SDEO with Kumaraswamy prior).

### J Fang et al.’s approach to sampling with *L*_0_ priors in rate networks

As noted in the Discussion, the factorization *p*(*c, z*) = *p*(*c*| *z*)*p*(*z*) is not the only way one can design a rate network to sample with an *L*_0_ prior. In particular, Fang *et al*. [51] proposed a Langevin sampling algorithm for sparse coding with an *L*_0_ prior. Here, we detail this model as applied to the olfactory sensing problem, following the discussion in Appendix E of our previous work [39]. Our starting point is a spike-and-slab prior on odor concentrations:

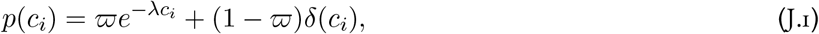

where for simplicity we will assume that *a priori* each odor is present with the same probability *ϖ*, and given that it is present its concentration is drawn from an exponential distribution of rate *λ*. Now, define an auxiliary variable **u** that is mapped to concentration estimates **c** via element-wise soft thresholding:

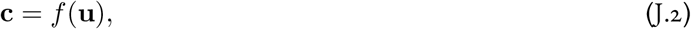

where

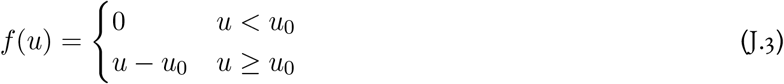

is the soft-thresholding function for threshold

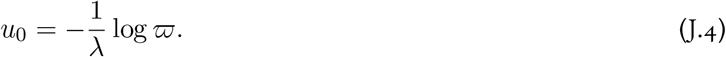

Given an observation **s**, we then run the following unconstrained Langevin dynamics for **u**:

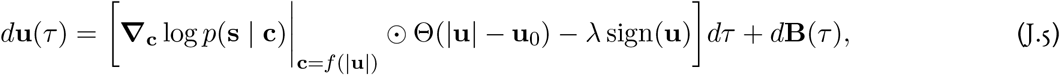

where **u**_0_ = *u*_0_**1**. For a Poisson likelihood as used elsewhere, we have

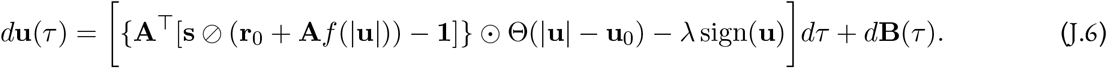

Like our model, this is a gated RNN, though of a different form.

**Figure S1:**
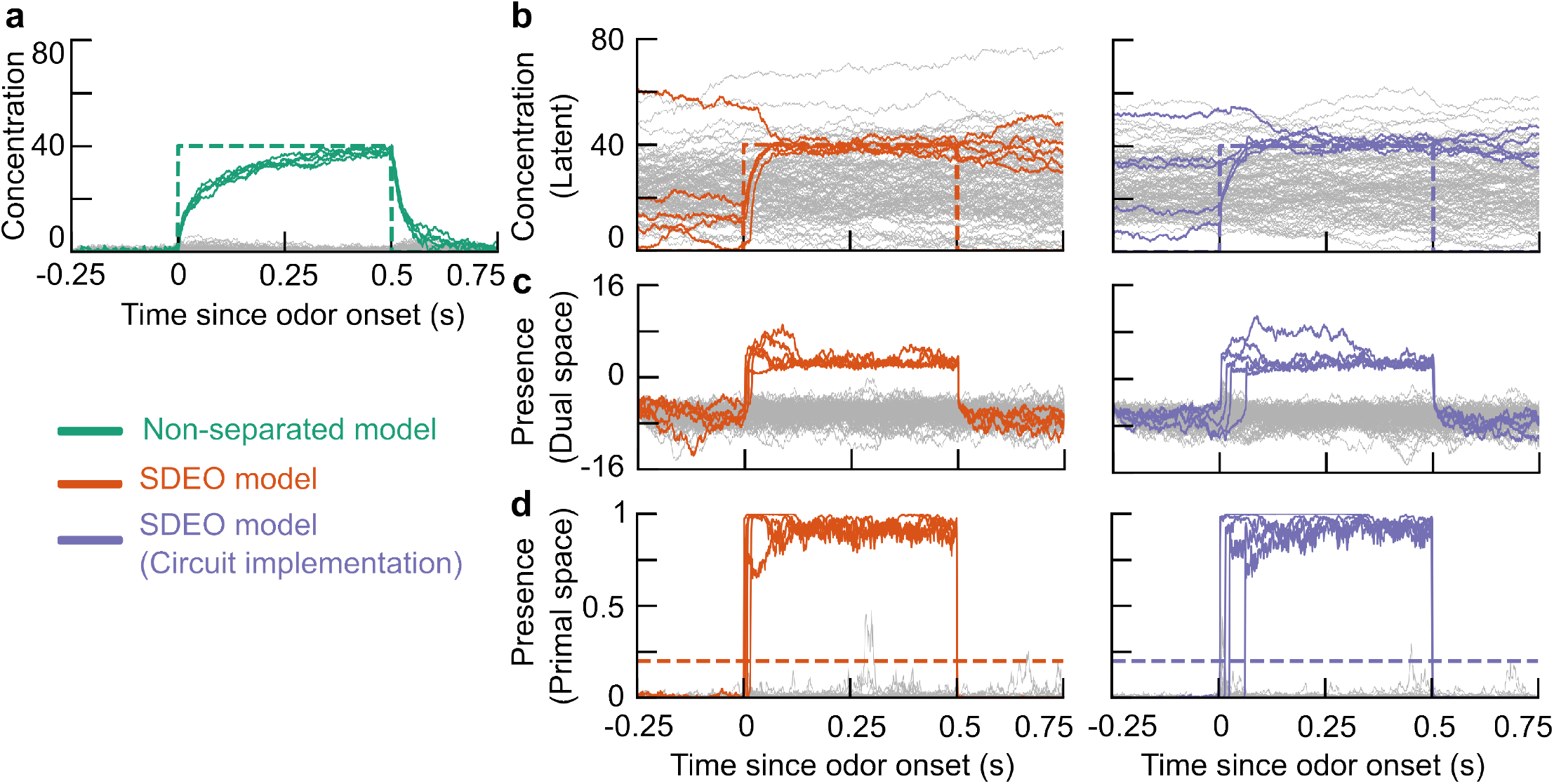
Dynamics of non-separated and SDEO models during the estimation process under sparse binary sensing matrices. We test three models— non-separated (as in [39]), SDEO, and SDEO with circuit implementation—on a simple estimation task to give qualitative illustration of the model dynamics during olfactory sensing. In the task, a randomly selected set of 5 out of 500 odorants appears at concentration 40 for a duration of 0.5 s. The three columns illustrate the dynamics of three models respectively, and the three rows show different quantities estimated. In each plot, the colored lines denote the values for the presented odorants, while the gray lines represent those for the background (non-presented) odorants. **a**. Estimated concentration. The dashed line traces true concentration over time. **b**. Estimated presence in the dual space. **c**. Estimated presence in the [0, 1]-bounded primal space. The dashed line marks the threshold used to binarize the presence variable during inference. Here we used sparse binary sensing matrices with sparsity 0.1 (defined in Appendix H). We ran the same simulation using dense Gamma sensing matrices and showed the results in Figure S2. For details of implementations, see Appendix I.4.2.

**Figure S2:**
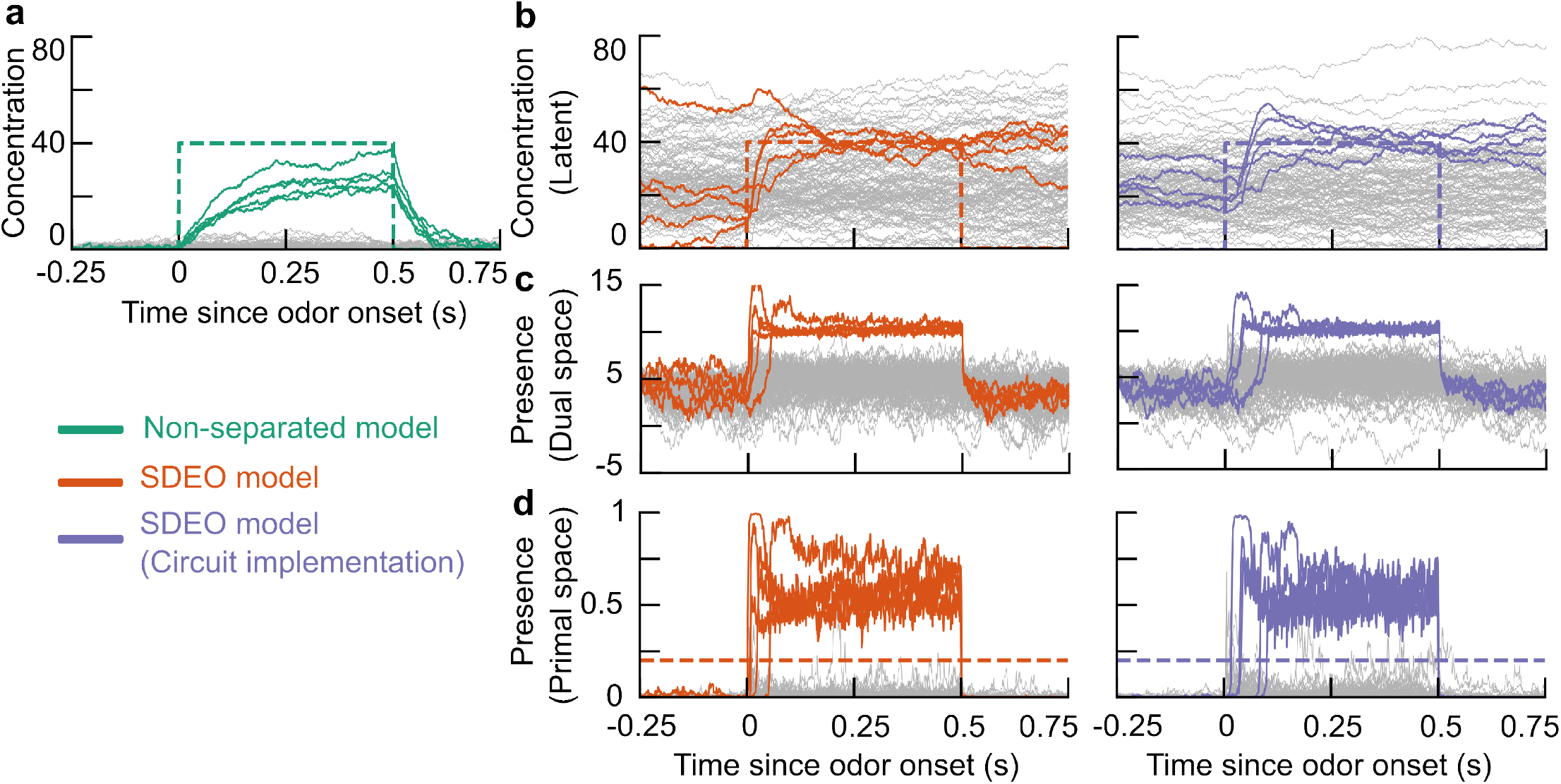
Dynamics of non-separated and SDEO models during the estimation process under dense Gamma sensing matrices. Here we re-ran the same simulation as in Figure S1 but used dense gamma affinity matrices instead. Hence, for details of the experiment, see the caption under Figure S1. **a**. Estimated concentration. The dashed line traces true concentration over time. **b**. Estimated presence in the mirror space. **c**. Estimated presence in the [0, 1]-bounded space. The dashed line marks the threshold used to binarize the presence variable during inference. For details of implementations, see Appendix I.4.2.

**Figure S3:**
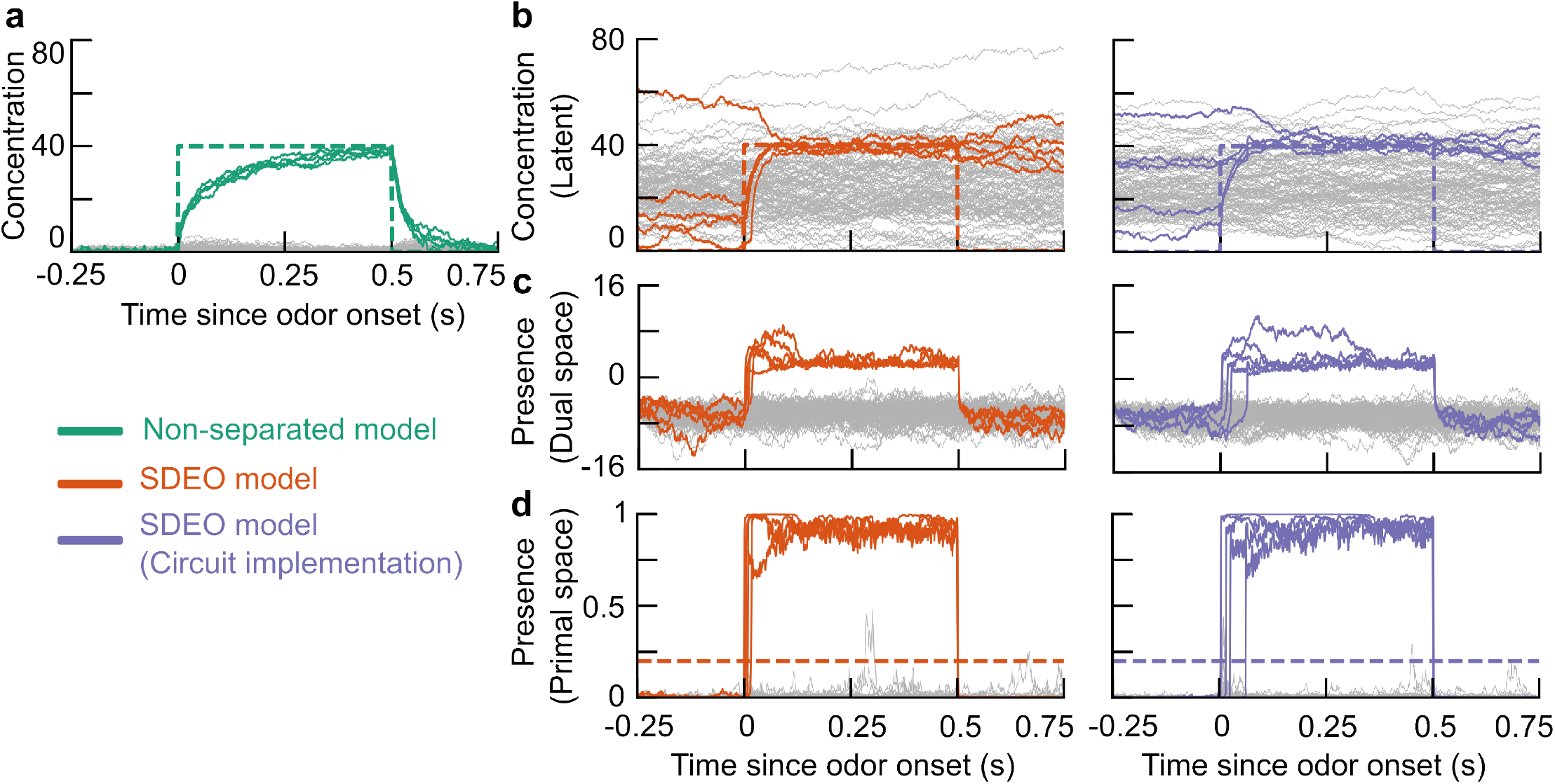
Dynamics of SDEO models using presence-dependent concentration priors with sparse binary and dense gamma affinity matrices. We set the rate of the exponential prior *λ* as 2.5. Once the odors disappear at 0.5s, the presence estimate falls below the threshold, and the exponential prior gradually suppresses the concentration estimate toward 0.

**Figure S4:**
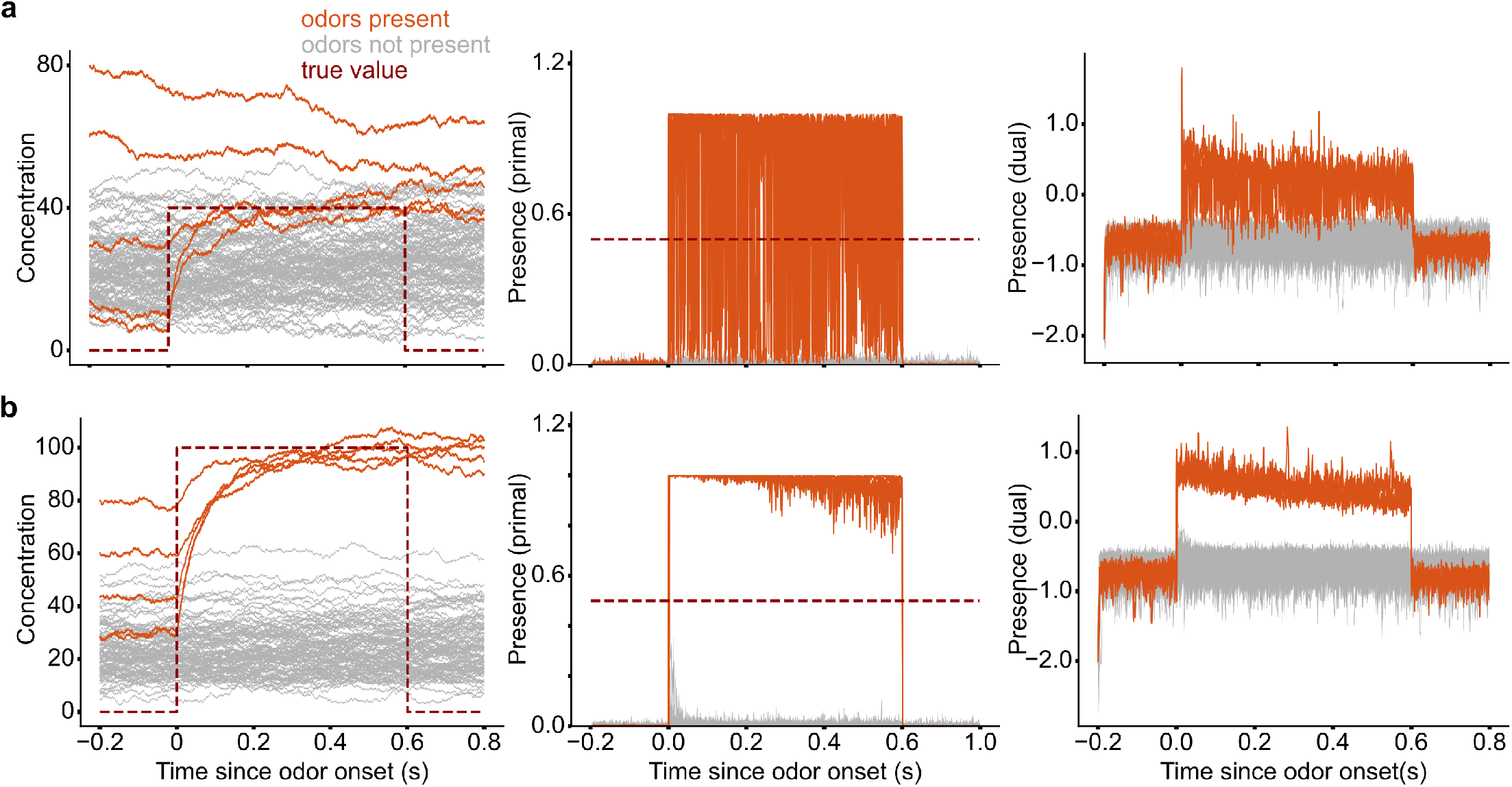
An ablation study was conducted by removing the hard-gated prediction-error term from concentration inference. In this variant, both concentration and presence inference relied on the same soft-gated prediction-error term. The model performed well when the true odor concentration was substantially higher than the prior expectation. However, when the true concentration was lower than, or close to, the prior, the model failed to infer the concentration and presence accurately.

**Figure S5:**
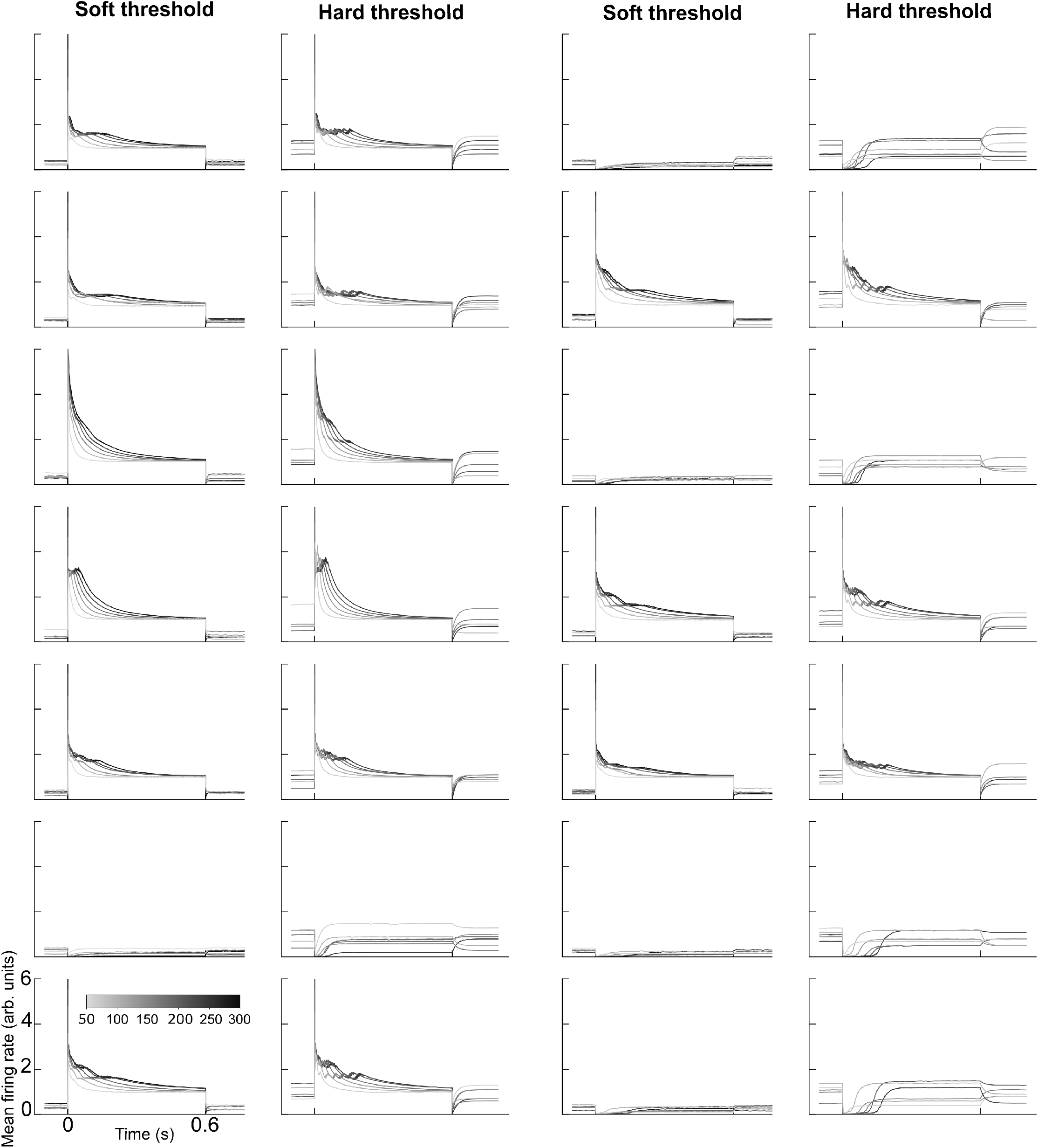
Responses of single projection neurons under different thresholding schemes. Each pair of panels depicts the corresponding neurons simulated with soft and hard gating, allowing direct comparison of their effects on firing dynamics. A set of fixed odor stimuli is present at 0s and withdrawn at 0.6s. Darker color indicates stronger stimulation.

**Figure S6:**
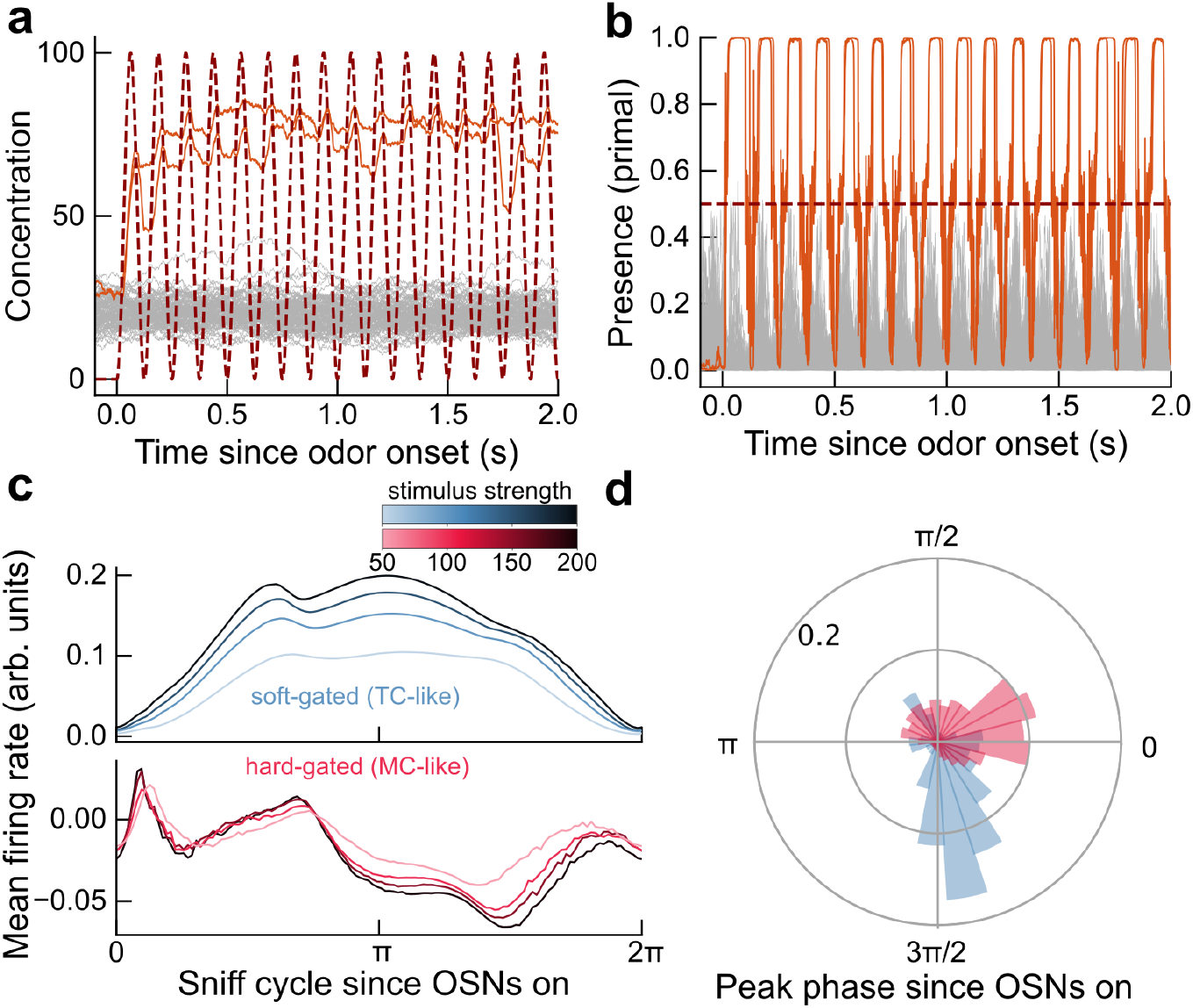
Sniff-coupled dynamics of the SDEO model. We repeated the same simulation shown in Figure 6 using a higher sniff frequency. External odor stimuli are held fixed, but the input concentration received by OSNs is modulated by sniffing, modeled here as a sinusoidal oscillation. **a**. Example concentration estimates across odors. Orange traces indicate present odors, gray traces indicate absent odors, and the dark-red dashed trace shows the sniff-modulated input concentration received by OSNs. The inferred concentration approximately tracks the true input over a few successive sniff cycles. **b**. Inferred odor presence for the same simulation. Present odors rapidly cross the decision threshold, the dark red dashed line, at an early phase of the sniff cycle, whereas absent odors remain mostly below threshold. **c**. Baseline-subtracted mean firing rate in a sniff cycle of hard-gated projection neurons (red) and soft-gated projection neurons (blue) over concentration. The soft-gated neurons exhibits a clear monotonic trend with stimulus strength. While the activity of hard-gated neurons are distributed more broadly without a clear mode. **d**. Distribution of single cell peak response phases for soft- and hard-gated projection neurons, showing distinct phase preferences between the two types.

**Figure S7:**
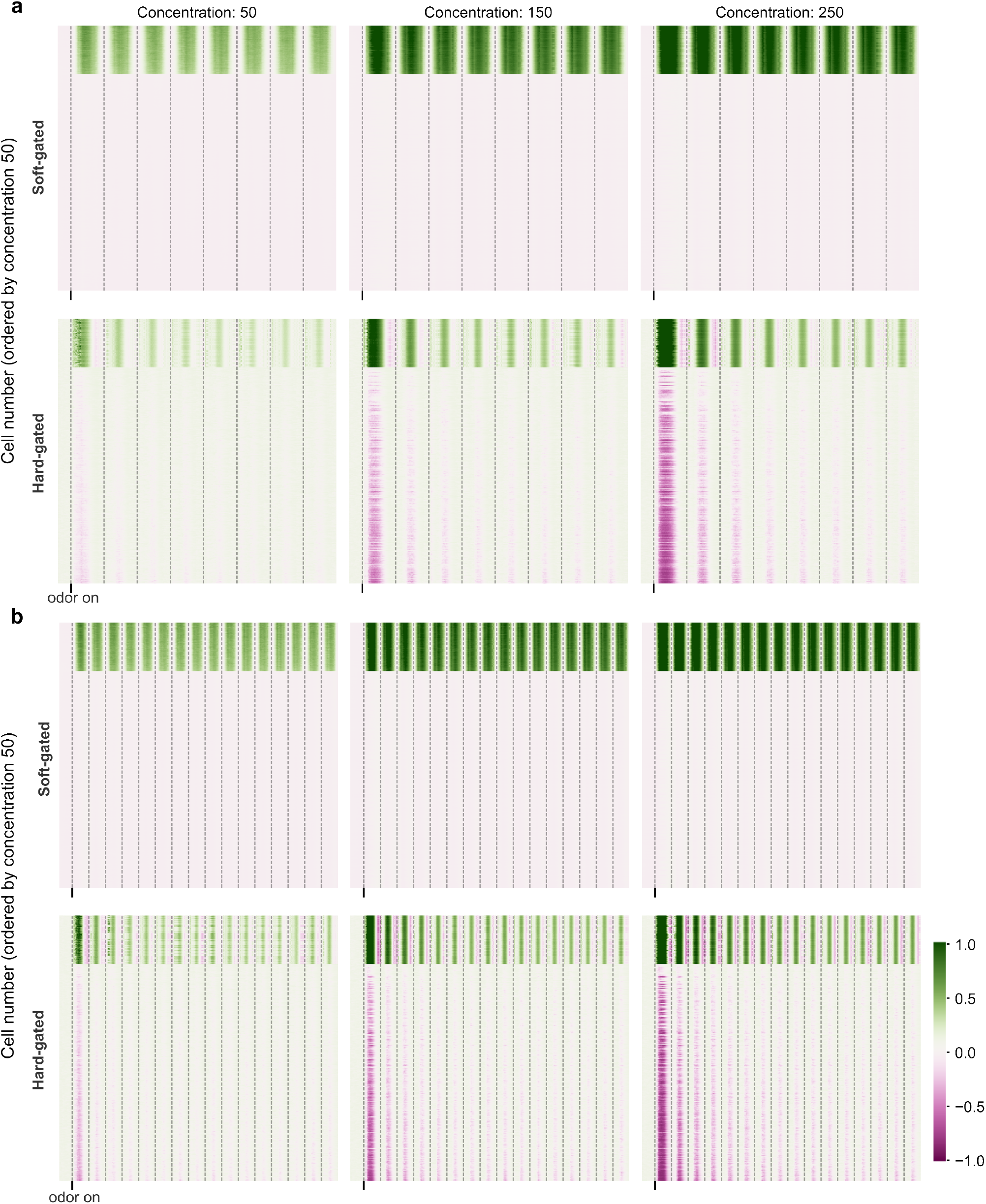
Baseline-subtracted activity of 300 soft-gated neurons and 300 hard-gated neurons, ordered by the mean response at concentration 50. The odor stimulus remains constant following onset. The dashed line indicates the onset of the sniff cycle. The three columns correspond to 3 different stimulus strengths, and we show examples at low sniff frequency (**a**) and high frequency (**b**). Similar to the single-cell activity shown in Figure 5, soft-gated neurons exhibit relatively more stable responses among non-activated neurons.

**Figure S8:**
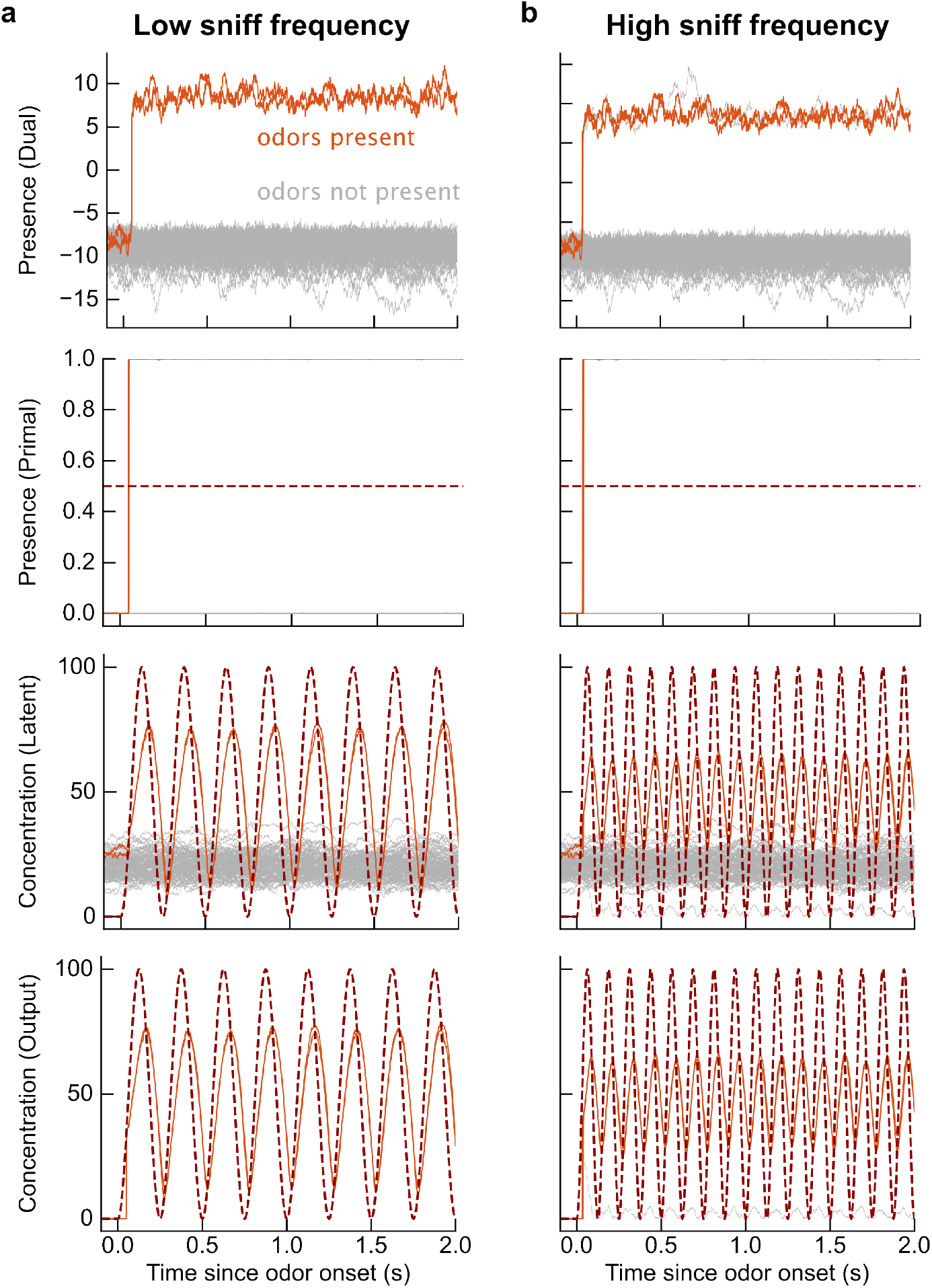
Dynamics of SDEO model using adaptive presence prior. Compared with static presence priors, such as Bernoulli distribution and Kumaraswamy distribution, the adaptive presence prior yields a more stable estimate of odor presence and allows the concentration estimate to more closely track the oscillations. This is particularly important in natural odor sensing, where odor concentration is highly fluctuating. We show examples at low sniff frequency (**a**) and high sniff frequency (**b**)

**Figure S9:**
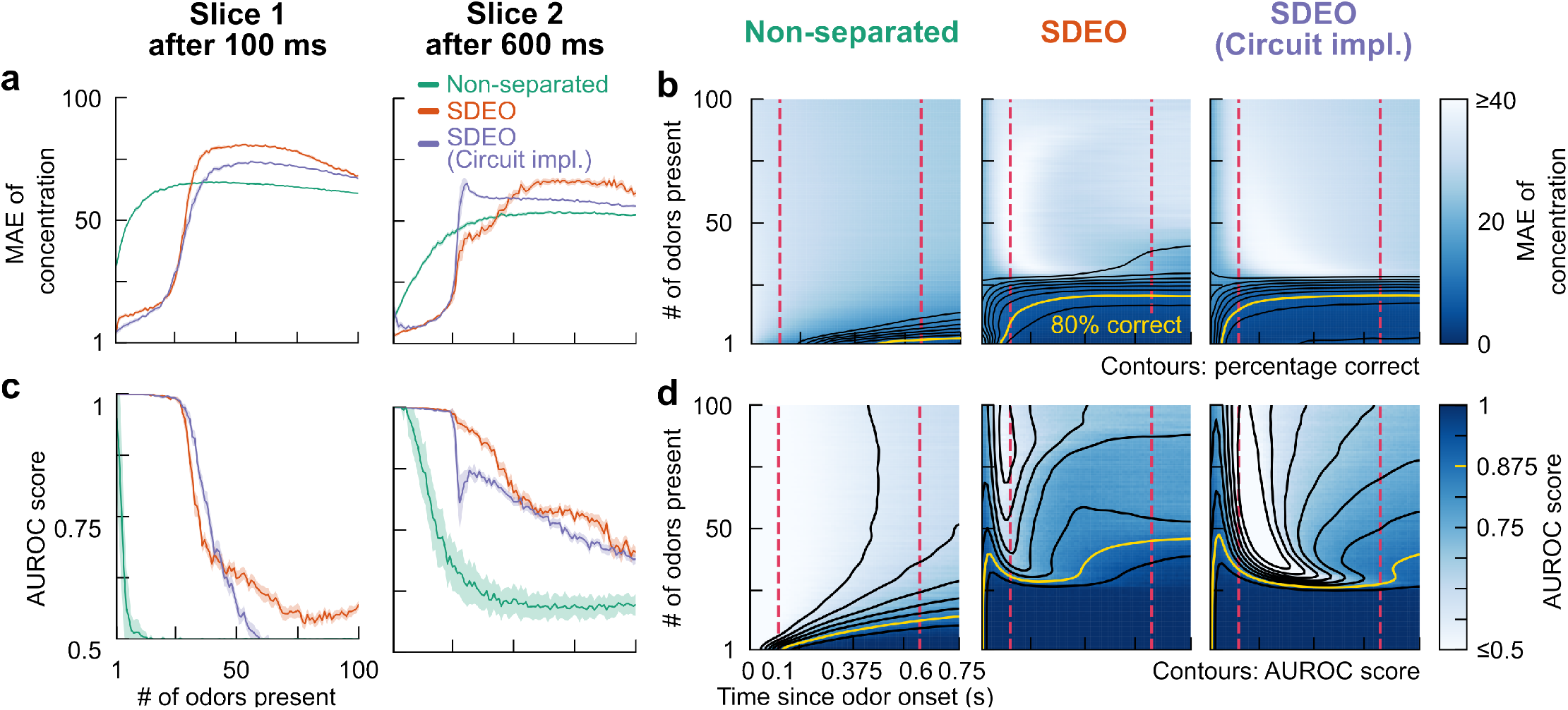
Improvement in fast detection of multiple odorants through separation of inference. We repeat the same simulation as in Figure 7 but using dense Gamma affinity matrices instead of sparse binary affinity matrices. We evaluate the same three models as in Figure 3 in a series of simulation where increasing numbers of odorants are simultaneously presented. In each run, in a set of 1000 odorants, a number of them are randomly selected and presented to the model for a duration of 0.75 s. We increase number of presented odors from 1 to 100, while repeat each setting for 40 times, compute the metrics and then take the average as final results. The shaded areas in **a** and **c** show *±*1.96 *·*SEM (representing 95% C.I) over realizations throughout. For numerical stability, here we use *α* = 1 for *α* in the Gamma prior distribution, while other hyperparameters remains unchanged as in Table 2. Row 1 assesses the models’ performance in odorants concentration estimation using mean absolute error. **a**. Mean absolute error of estimated concentration as a function of the number of odorants present at two timepoints after odor onset. **b**. Heatmap of mean absolute error over inference time and number of presented odorants, with smoothed contours of correct detection fraction overlaid. Row 2 assesses the models’ performance in odorants presence estimation. For the non-separated model, we convert the concentration estimation into presence estimation by binarizing the estimated concentrations based on whether they exceeds half of the true odorant concentration. Since presence estimation is a binary classification task, we use AUROC score as the performance metric. **c**. AUROC score as a function of the number of odors present at two timepoints after odor onset. **d**. Heatmap of AUROC score over inference time and number of presented odors, with smoothed contours overlaid. For details of implementation, see Appendix I.4.3.

**Figure S10:**
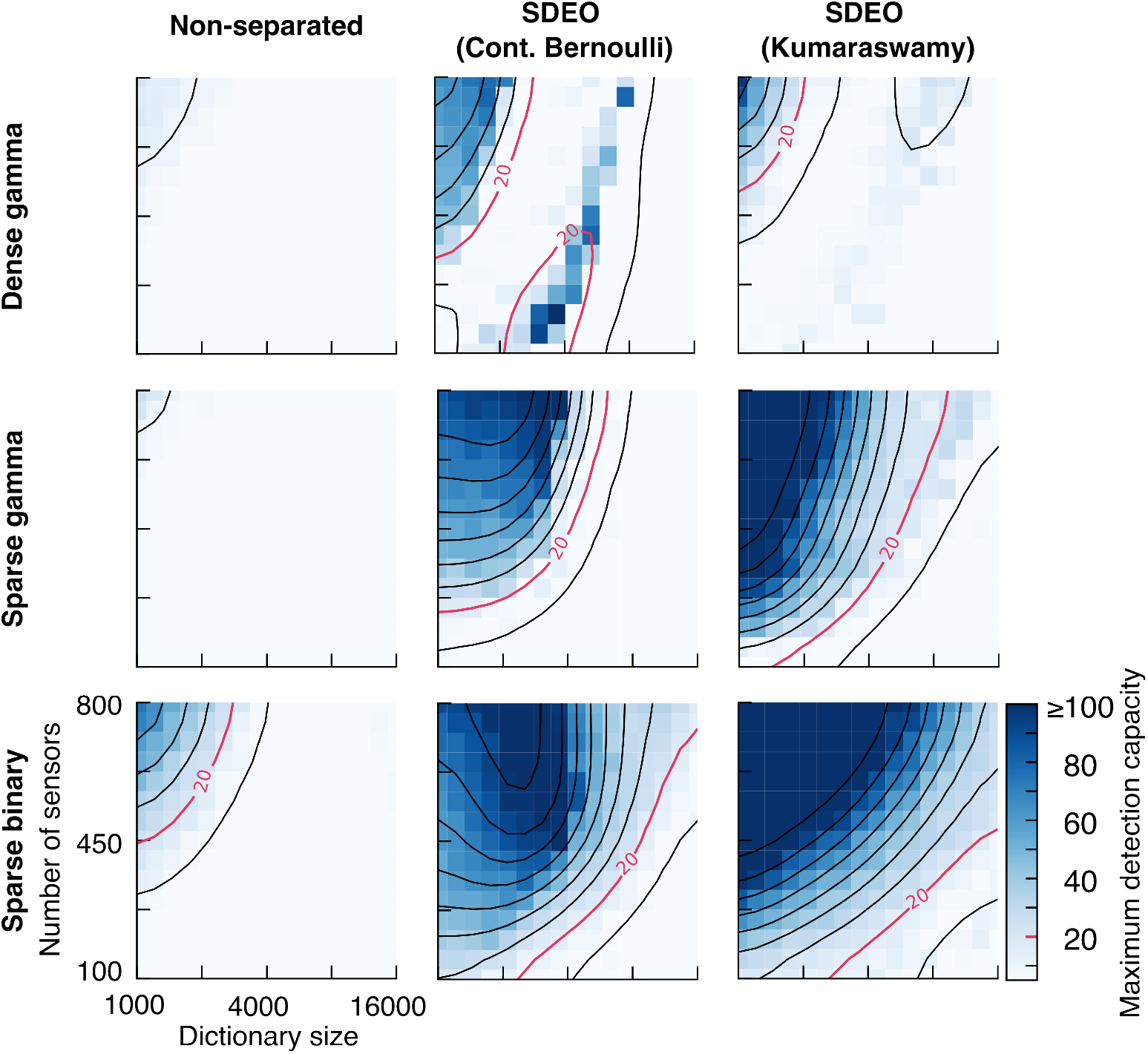
Scaling of detection capacity with dictionary size and sensor repertoire for different priors and sensing matrix models. We run the same simulation as in Figure 8, but showing maximum detection capacity assessed by concentration instead of presence estimates. The three columns correspond to three models—non-separated, SDEO, and SDEO with truncated KS prior on the soft presence—and the three rows correspond to three types of affinity matrices. These are: dense Gamma, whose entries are i.i.d. random variable following Gamma(0.37, 0.36); sparse Gamma, obtained by applying a 0.1 sparsity mask to a dense Gamma matrix; and sparse binary, whose entries are i.i.d. random variable following Bernoulli(0.1). Each heatmap shows the maximum detection capacity assessed by concentration estimates for combinations of sensors counts (from 100 to 800, equally spaced linearly) and dictionary size (1000 to 16000 equally spaced on a log scale). The maximum detection capacity *κ*_MAE_ is defined as the largest number of simultaneously presented number of odorants that the model can detect with a mean absolute error *≤*10, while *c*_*True*_ = 40. Smoothed contours are overlaid and can be interpreted as the required number of sensors to maintain a certain capacity as a function of dictionary size. The total inference time duration is 0.5 s for all runs, and the value in each cell of the heatmap is the average of three independent runs. For details of implementation, see Appendix I.4.4.

**Figure S11:**
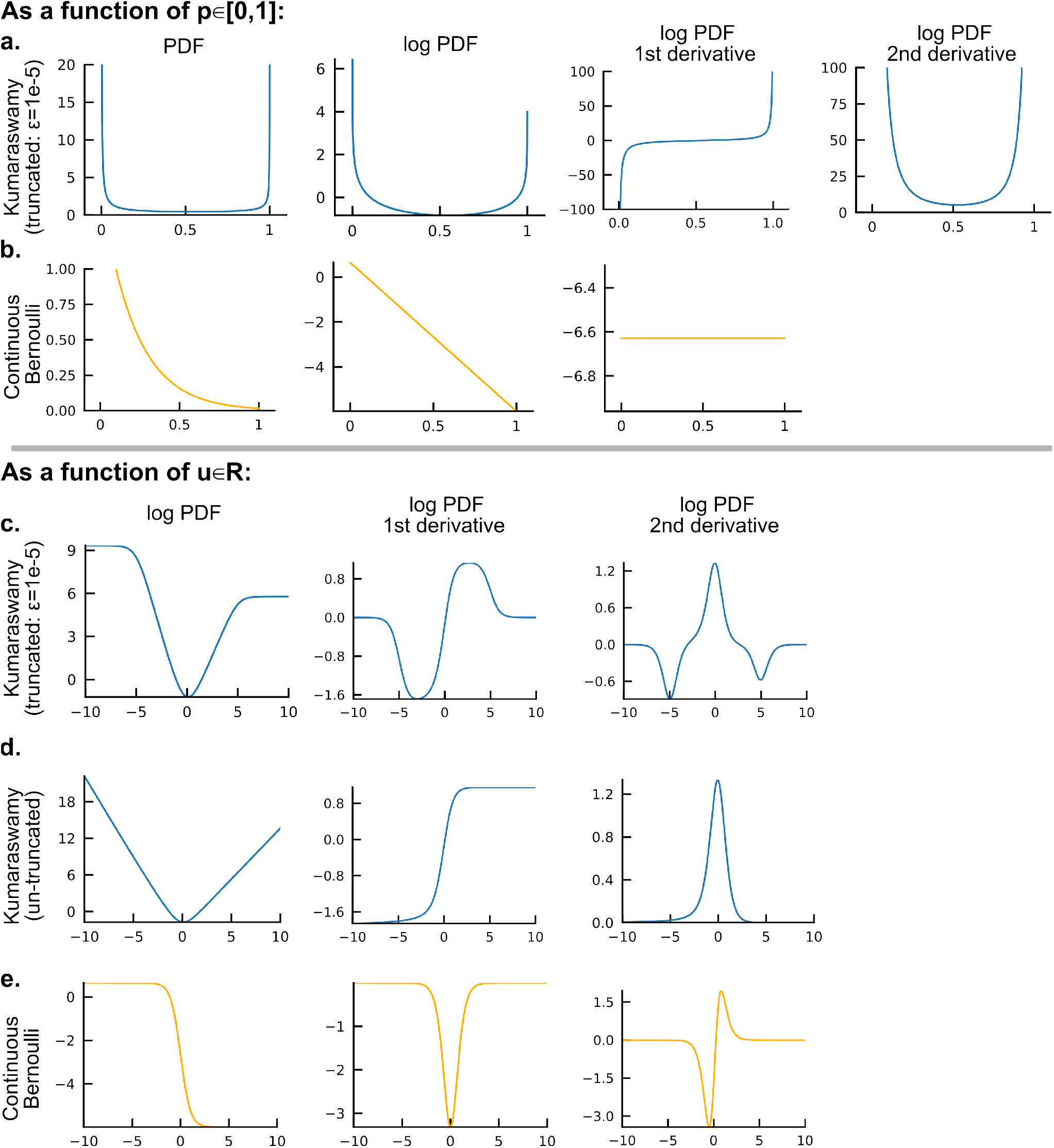
In **a**. and **b**., we plot the Truncated Kumaraswamy distribution and continuous Bernoulli distribution on [0, 1]. **a**. Truncated Kumaraswamy distribution with parameters *a* = 0.055, *b* = 0.422, *ε* = 1*e −*5. The four columns from left to right respectively are: probability density function (PDF), log PDF, the first derivative of log PDF, and the second derivative of log PDF. **b**. Continuous Bernoulli distribution with *ϖ* = 0.01. The three columns from left to right respectively are: probability density function (PDF), log PDF, and the first derivative of log PDF. We didn’t show the second derivative of log PDF because it vanishes. In **c**., **d**. and **e**. we plot three distributions with respect to *u ∈* ℝ such that 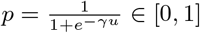. This is to illustrate the score function in the dual space. The three columns from left to right respectively are the log PDF, the first derivative of the log PDF, and the second derivative of the PDF. **c**. Truncated Kumaraswamy distribution with the same parameters as in **a. d**. Un-truncated Kumaraswamy distribution with parameters *a* = 0.055, *b* = 0.422, *ε* = 0. **e**. Continuous Bernoulli distribution with *ϖ* = 0.01.

**Figure S12:**
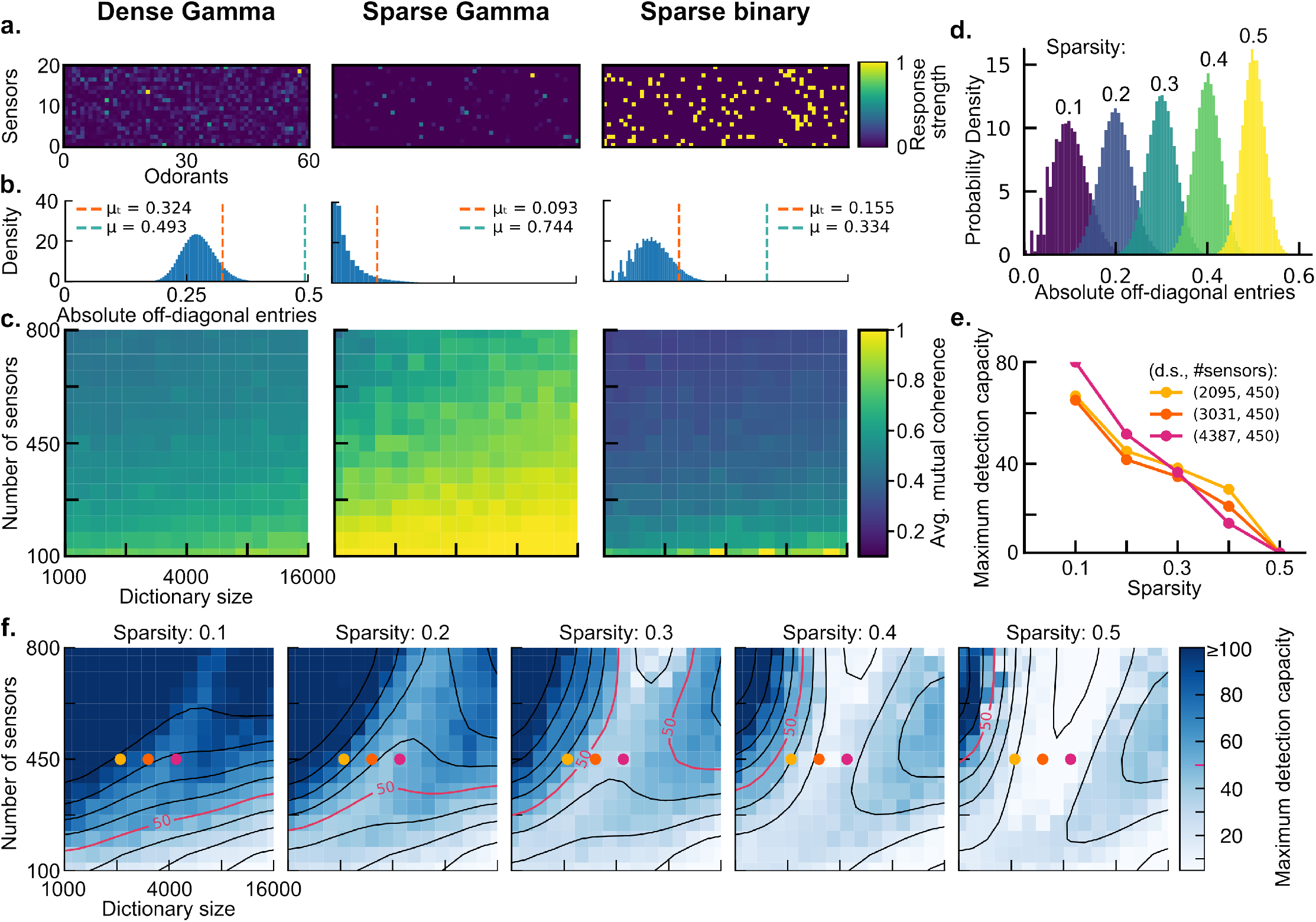
**a**. Zoomed in example of the three types of sensing matrix—dense Gamma with Gamma(0.36, 0.37), sparse Gamma with Gamma(0.36, 0.37) and sparsity 0.1, and sparse Binary with sparsity 0.1. **b**. Histograms showing the distribution of absolute off-diagonal entries of the Gram matrix of the sensing matrices, which is equivalent to column-wise correlation in the sensing matrix. Three shows distinct sensing matrix types matching the column titles. Here we use a system with 600 sensors and 5000 dictionary size. The worst case correlation *µ*, namely the mutual coherence, is indicated by the green vertical dashed line overlaid to the histogram, and also printed out in the legend. The average of the top 20% largest correlation *µt*, is indicated by the orange vertical dashed line overlaid to the histogram, and also printed out in the legend. **c**. Mutual coherence of the sensing matrices under different dimensionality, averaged across 5 random trials. **d**. Histogram showing the probability density of absolute off-diagonal entries of sparse binary sensing matrices with different sparsity. **e**. Maximum detection capacity as a function of sparsity of the sparse binary sensing matrix, under different combination of dictionary size and sensor repertoire. The dimensionality is indicated by dots with corresponding color in panel **f. f**. Heatmap of maximum detection capacity under different dictionary size and sensor repertoire when using sparse binary sensing matrices with increasing sparsity (s=0.1 to 0.5)

## Footnotes

1 FN denotes false negatives (missed detections), and FP denotes false positives (false alarms).

